# Enhanced compound-protein binding affinity prediction by representing protein multimodal information via a coevolutionary strategy

**DOI:** 10.1101/2022.04.06.487274

**Authors:** Binjie Guo, Hanyu Zheng, Haohan Jiang, Xiaodan Li, Naiyu Guan, Yanming Zuo, Yicheng Zhang, Hengfu Yang, Xuhua Wang

**Affiliations:** Department of Neurobiology and Department of Rehabilitation Medicine, First Affiliated Hospital, Zhejiang University School of Medicine, Hangzhou, Zhejiang Province 310058, China; Liangzhu Laboratory, MOE Frontier Science Center for Brain Science and Brain-machine Integration, State Key Laboratory of Brain-machine Intelligence, Zhejiang University, 1369 West Wenyi Road, Hangzhou 311121, China; NHC and CAMS Key Laboratory of Medical Neurobiology, Zhejiang University, Hangzhou 310058, China; Co-innovation Center of Neuroregeneration, Nantong University, Nantong, 226001 Jiangsu, China; School of Computer Science, Hunan First Normal University, Changsha, 410205 Hunan, China

**Keywords:** Compound-protein binding affinity prediction, coevolutionary strategy, protein multimodal information, Protein 3D structure, Computational model, Artificial intelligence

## Abstract

Due to the lack of a method to efficiently represent the multimodal information of a protein, including its structure and sequence information, predicting compound-protein binding affinity (CPA) still suffers from low accuracy when applying machine learning methods. To overcome this limitation, in a novel end-to-end architecture (named FeatNN), we develop a coevolutionary strategy to jointly represent the structure and sequence features of proteins and ultimately optimize the mathematical models for predicting CPA. Furthermore, from the perspective of data-driven approach, we proposed a rational method that can utilize both high- and low-quality databases to optimize the accuracy and generalization ability of FeatNN in CPA prediction tasks. Notably, we visually interpret the feature interaction process between sequence and structure in the rationally designed architecture. As a result, FeatNN considerably outperforms the state-of-the-art (SOTA) baseline in virtual drug screening tasks, indicating the feasibility of this approach for practical use. FeatNN provides an outstanding method for higher CPA prediction accuracy and better generalization ability by efficiently representing multimodal information of proteins via a coevolutionary strategy.

## Introduction

Since it is time and resource consuming to experimentally assess compounds and target protein binding affinities during drug discovery and development, effective virtual screening approaches using computational methods could greatly accelerate the drug candidate identification process by learning the abstract binding information between drug and target and accurately predicting compound-protein binding affinities (CPA) [1, 2], especially in cases where great numbers of sources for compound and protein interaction data are available through open source databases. For instance, BindingDB [3] currently provides a comprehensive collection of experimentally measured binding affinity data including more than 1 million protein–ligand complexes in the Protein Data Bank (PDB) [4], which substantially increases the potential for *in silico* CPA prediction. However, even with these abundant data, accurately predicting CPA is still the fundamental challenge preventing this method from being used in practical drug candidate screening applications due to the lack of a method to efficiently extract features from the data. To increase the accuracy of CPA prediction, the development of computational methods has proceeded with a variety of protein information embedding and representation strategies [5–8]. Despite substantial advancements, these strategies have met challenges with respect to further increasing the accuracy of CPA prediction.

Initially, researchers tended to represent protein features only using the protein sequence information, namely, the target (protein) is regarded as a sequence of residues. In these models, a pairwise array with the residue features of the protein as its column (or row) and the SMILES sequence information of the compound as its row (or column) is often utilized as the attention matrix to learn the potential interaction between a protein and a compound [9]. Typically, these models rely on the sequence information of the compounds and proteins of interest to learn their interactions via pairwise matrices, with the aim of predicting the binding affinities between them [9–13]. For example, multilayer 1-dimensional convolutional neural networks (1D-CNNs) are utilized to extract the features from the residue sequences of proteins, and the obtained vectors are used to represent the features of proteins, predict the CPAs and intensively study the noncovalent interaction between the ligand and binding target [14–16]. However, in addition to a protein’s sequence of residues, the 3D structure of a protein also contributes significantly to its features [17, 18]. Therefore, neglecting the 3D spatial structure information of the protein may prevent the full realization of the potential of computational modeling in CPA prediction.

In this scenario, the approaches of representing and embedding protein structure information have been tentatively proposed to improve the accuracy in CPA prediction. To do so, molecular docking simulation methods [19, 20] based on background molecular dynamics knowledge and structure-based machine learning methods [8, 21] have been proposed. Relying on the knowledge of biophysics, the docking method computationally simulates the potential binding sites and 3D structures of compound-protein complexes, so it heavily depends on high-quality 3D protein structure data during CPA prediction [22, 23]. Despite a few successful stories, this method is severely limited due to the scarcity of high-quality 3D structure data of proteins (the precise position of each atom in a protein) [24]. By contrast, machine learning algorithm-based approaches can use 3D protein structure data with either high or low resolutions (the positions of key atoms in a protein). These models are fed with the spatial 3D information of the proteins in order to attain a superior ability to predict CPA [25–27]. For instance, the structural features of proteins were extracted through 3D atomic representations in voxel space by applying 3D CNNs [8]. However, the performance of these models was not significantly improved by introducing the structural information of the proteins [6, 8]. We hypothesized that this was due to the lack of the comprehensive consideration of the multimodal information (both sequence and structure information) of the protein by these methods. To address this problem, we sought to develop a method that can rationally incorporate the multimodal information of protein into CPA prediction models in order to improve CPA prediction performance.

Inspired by the multi-feature fusion tactics via coevolution [28], we designed an end-to-end neural network architecture (Fig. 1), named the fast evolutional aggregating and thoroughgoing graph neural network (FeatNN). Through the coevolutionary strategy, FeatNN efficiently represented the multimodal information (containing both structure and sequence information) of proteins and thus overcame the multimodal protein information representation challenge. Upon the IC_50_ and KIKD datasets generated from PDBbind [29], FeatNN outperforms the SOTA method (MONN) in CPA prediction tasks by 21.33% and 17.07% with respect to the R^2^ metric, 6.16% and 2.98% in terms of the root mean square error (RMSE), and 7.00% and 5.45% in the Pearson coefficients, respectively (Fig. 2).

**Fig. 1.**
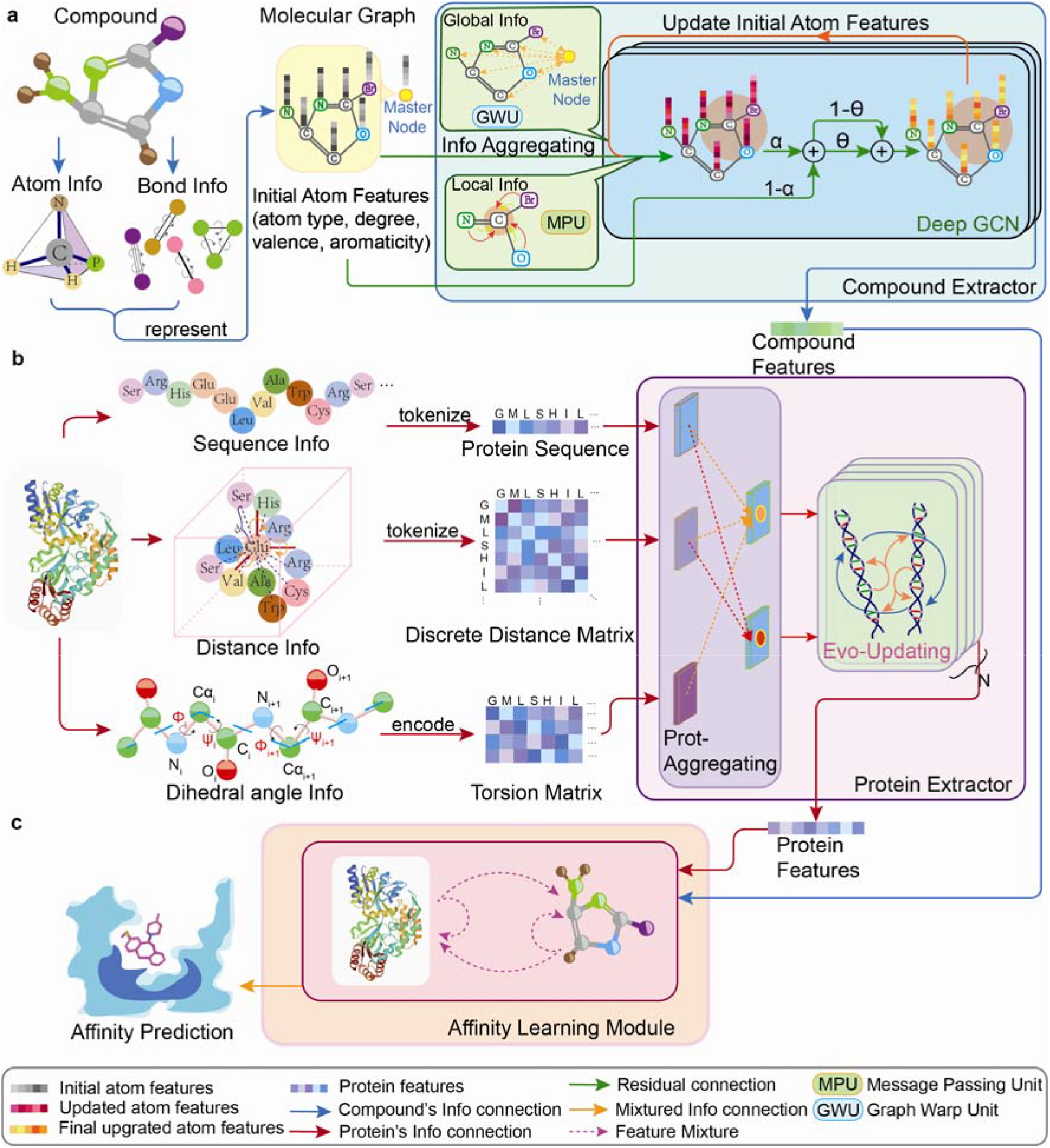
Architecture overview of FeatNN. **a.** The atom and bond information of a given compound is encoded into a molecular graph, which acts as the input for the compound extractor module to distill its features. The compound extractor includes a deep GCN block (Supplementary Fig. 12) and multihead attention blocks (Supplementary Fig. 14). **b.** The features of a protein are embedded with matrices and vectors as inputs to the Prot-Aggregation module (Supplementary Fig. 17), whose outputs are then fed to the Evo-Updating module (Supplementary Fig. 18), which co-evolutionarily updates the structure and sequence features. Both the Prot-Aggregation module and the Evo-Updating module form the protein extractor block. **c.** The extracted atom and residue features are processed by the affinity learning module (Supplementary Fig. 20), which also enables FeatNN to learn the potential interaction features between the atoms of the compound and the residues of the protein. Additionally, the sets of information derived from the atom features and residue features are integrated through the affinity learning module to predict the CPA. The parameter settings of FeatNN are shown in Supplementary Table 2.

**Fig. 2.**
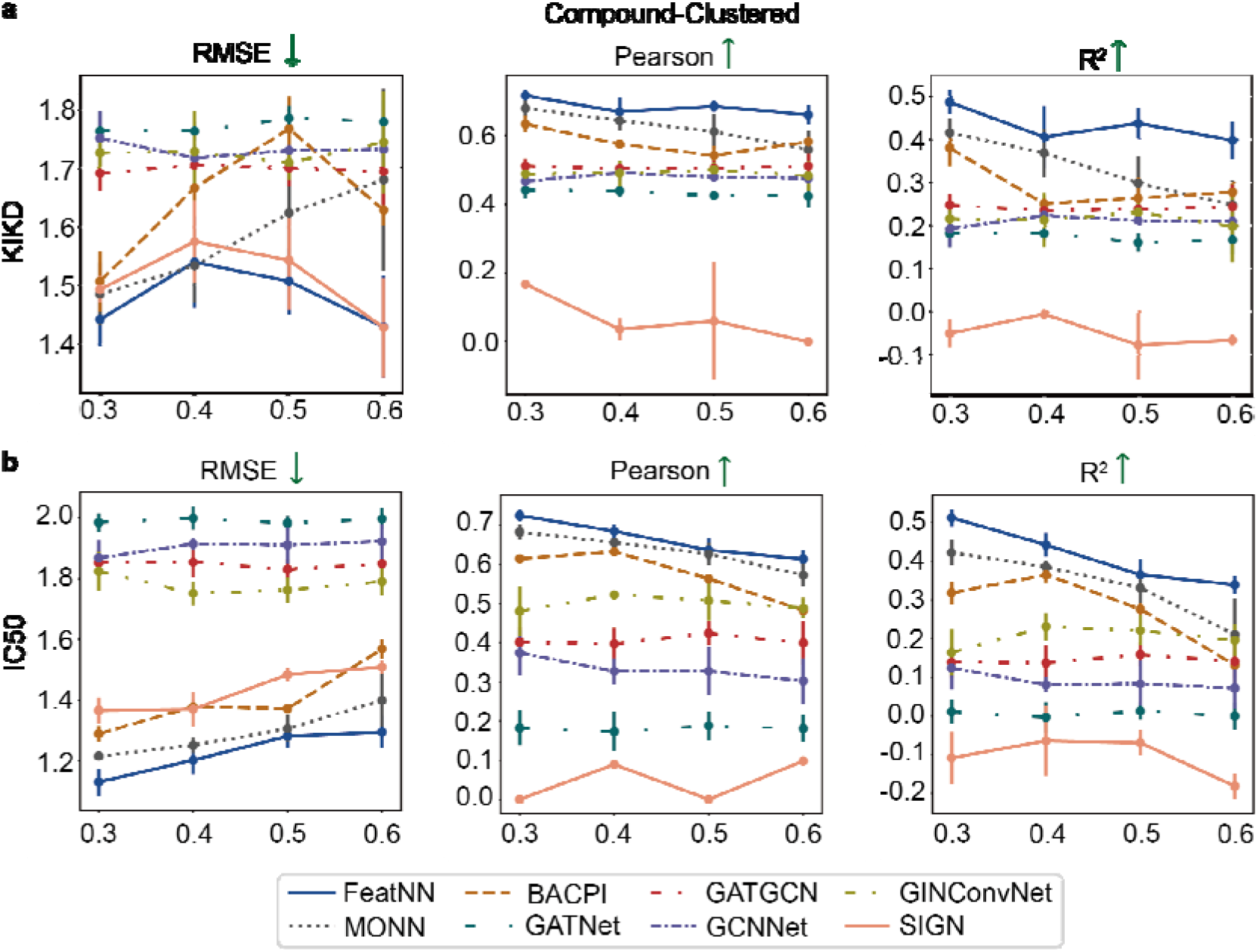
Evaluation of FeatNN, BACPI, SIGN, GraphDTA (GATNet, GATGCN, GCNNet, GINConvNet) and MONN. Performance evaluated on compound-clustered strategy datasets with similarity thresholds of 0.3, 0.4, 0.5 and 0.6 constructed from PDBbind with KIKD and IC_50_ measurement, respectively. The benchmark dataset is generated from PDBbind (version 2020, the general set) and contains 12,699 compound-protein pairs. Performance results are plotted as the mean values and standard deviations (SD) by 5-fold cross-validation strategy with 10 independent experiments. Each point represents the independent experimental group mean with error bars indicating SD. We choose the three indicators (the RMSE, Pearson coefficient, and R^2^) that can best evaluate the prediction performances of the methods in terms of the continuous values (CPA) they predicted. **a.** Performances evaluated on the dataset generated from PDBbind with KIKD measurement. **b.** Performances evaluated on the dataset generated from PDBbind with IC_50_ measurement. Please note that the results of SIGN present here were different from the results reported by the original literature [26], possibly because we use PDBbind-v2020 as our benchmark database instead of PDBbind-v2016 used in their study. In addition, considering the biology means behind the data, we split the dataset into two parts (“IC_50_” and “KIKD” [9]) instead of simply mixing the affinity measured with “IC_50_”, “K_i_”, and “K_d_” together in their study. Moreover, we applied compound-cluster and protein-cluster strategies in our study to avoid data leakage caused by the biology-correlated knowledge (similarity structure or sequence in protein or compound). In most case, MONN achieved the best performances in baselines; therefore, we consider MONN as the SOTA baseline in our paper.

The major technical advances of FeatNN are listed as follows.

1. An Evo-Updating block is employed in the protein encoding module to interactively update the sequence and structure information of proteins so that the high-quality features of proteins are extracted and presented, enabling FeatNN to outperform the SOTA model by great margins exceeding 21.33% in R^2^.
2. In FeatNN, the distance matrices of protein residues are discretized into one dimension, and the word embedding strategy is applied to encode protein structure information, so that the network could effectively represent the multimodal protein information and lower the computational cost simultaneously.
3. With respect to the extraction of compound features, a specific residual connection is applied to represent the molecular graph, in which the features of the initial nodes are added onto each layer of the GCN [30], such that the graph features representation limitation caused by the notorious oversmoothing problem in traditional deep GCNs is solved.
4. With the pretraining and fine-tuning strategy, the R^2^ performance of the optimized model, FeatNN^optm^, further increases by 3.29% on average compared to that of FeatNN.
5. FeatNN has excellent generalization in the affinity prediction task, which is vital and pivotal in the drug screening domain. Targeting severe acute respiratory syndrome coronavirus 2 (SARS-CoV-2) 3-chymotrypsin (3C)-like protease and Akt-1, the generalization of FeatNN vastly outperforms the SOTA baseline in the affinity value prediction task.
6. The prediction results of FeatNN with different conformations of the same protein are robust when 3D structure information is directly introduced in the model while neglecting the molecular dynamics of the protein.

## Materials

### Dataset Construction

Even though PDBbind [31], BindingDB [3] and Binding MOAD [32–34] databases (Supplementary Fig. 2 and Supplementary Table. 1) contain paired information of protein-ligand complexes with structural data and the corresponding binding affinities, it was necessary to eliminate some data to comply with the quality standards of our model and baselines. The exclusion criteria included protein PDB file defects, and sequence information inconsistency in UniProt and PDB. Based on these criteria, we constructed a benchmark dataset based on PDBbind (version 2020, the general set) [29] that contains 12,699 compound-protein pairs. Meanwhile, a refined dataset [31] with higher quality of structural information has also been constructed from PDBbind (version 2020, the refined set, see Supplementary Fig. 2f). Additionally, we generated another dataset based on BindingDB (version Feb 6, 2022; the general set) [3] that is rich in data on compound-protein paired complexes but poor in protein diversity. The complex structure information in such dataset is not strictly paired and remains low-quality, because not all complexes in BindingDB have strictly paired 3D structure conformations, and most of these complexes correspond to multiple protein conformations with different PDB entries. Therefore, we preferentially chose the ligand-free or high-resolution PDB file for these complexes without strict correspondence between protein and compound. This generated dataset contains more than 210 thousand compound-protein pairs (Supplementary Table 1). To test the generalization ability of the models, we constructed new datasets from the Binding MOAD (see in Supplementary Table 1) database and excluded the complexes that appear in the datasets (train, validation, and test datasets) constructed from PDBbind (Supplementary Fig. Table 1). An affinity value of a certain measurement type (i.e., K_i_, K_d_, or IC_50_.) for each complex was provided, and “KIKD” was used to refer to the combination of K_i_-measured data and K_d_-measured data due to their high homogeneity. More details about the dataset construction process are available in the Supplementary Methods 3.3.

**Table 1.** Performance evaluation of different prediction approaches on the dataset generated from BindingDB. We apply RMSE, Pearson and R^2^ to evaluate the CPA prediction performances. The results of each group were counted with 10 independent experiments. The mean value (and SD) of each independent experimental group are shown in the table. Note: The SIGN is highly dependent on the structure information of the complex and binding pockets while most structure information recorded in BindingDB is redundant and low-quality (lack of the information of pocket and binding site to represent the complex graph as the input training data), it is difficult to process the data before training the SIGN. Therefore, we did not train the SIGN on BindingDB.

| **Model** | **$R^2$ ↑** | **RMSE ↓** | **Pearson ↑** |
| --- | --- | --- | --- |
| **FeatNN** | 0.719 (0.003) | 0.765 (0.004) | 0.850 (0.001) |
| **MONN** | 0.706 (0.004) | 0.783 (0.005) | 0.844 (0.002) |
| **BACPI** | 0.577 (0.005) | 0.935 (0.006) | 0.769 (0.002) |
| **GATGCN** | 0.543 (0.015) | 0.992 (0.016) | 0.742 (0.012) |
| **GCNNet** | 0.510 (0.023) | 1.030 (0.023) | 0.717 (0.015) |
| **GINConvNet** | 0.451 (0.124) | 1.080 (0.119) | 0.669 (0.094) |
| **GATConvNet** | 0.327 (0.027) | 1.200 (0.024) | 0.585 (0.001) |

### Training Data Generation

The PDBbind-based (both the general and refined datasets) training dataset generation process included three key steps. 1) Before performing data cleaning, we first assessed whether the regression labels (CPA values) in both PDBbind and BindingDB followed normal distributions to avoid the potential prediction deviation problem (Supplementary Fig. 2); 2) We then clustered the input compound and protein information according to a certain threshold (0.3, 0.4, 0.5, and 0.6) [9] to avoid the potential data leakage problem that can occur due to data similarities. In this evaluation, we assessed the similarity of the proteins using their multi-sequence alignment (MSA) scores and calculated the similarity of the compounds based on their fingerprints. Then, the same kinds of compounds or proteins with a certain threshold were divided into the same dataset; the details of this process are provided in Supplementary Methods 3.4 and 3.5; 3) Finally, we used a 5-fold cross-validation strategy [35] to generate training datasets to alleviate the potential overfitting problem. Then, the dataset was randomly shuffled with a training-validation-testing splitting ratio of approximately 7:1:2. For the generation of the BindingDB-based training dataset, we directly shuffled and split the dataset with the same training-validation-testing splitting ratio. The datasets generated from Binding MOAD were only used for testing the models’ generalization ability and transferability.

### Baseline Methods

To assess the performance of FeatNN, we chose to represent the SOTA algorithm architecture with the multiobjective neural network (MONN) [9], the structure-aware interactive graph neural network (SIGN) [26] and chose two classic methods, the drug-target binding affinity graph neural network (GraphDTA) [36], the bidirectional attention neural network for compound-protein interaction (BACPI) [37] as our baseline models. We followed the same experimental settings as those used in in the original studies that reported these baseline models.

- MONN applies a GCN block [30] to extract compound features and a 1D-CNN block to extract protein features and then constructs a pairwise matrix from the features of compounds and proteins to describe noncovalent interactions and predict CPA.
- GraphDTA comprises four models: the graph attention network (GATNet), graph convolutional network (GCNNet), the combined GAT and GCN (GATGCN) and graph isomorphism network (GINConvNet), all of which utilize architectures with a GCN block and an attention mechanism to extract protein and compound features and finally predict CPA through several dense layers that aggregate the features of compounds and proteins.
- BACPI serves as a bidirectional attention neural network and uses a 1D-CNN block to extract protein features from residue sequences and a graph attention network to extract compound features. CPA is predicted through several dense layers; this is similar to the GraphDTA approach.
- SIGN is as a structure-based method that converts the protein-ligand complex into a complex interaction graph and extract its features from such graph. The training data for this model must strictly contain the pair data (both protein and compound) in a complex with high-quality structure information.

## Results

### The Design of FeatNN with Input Protein Sequence and Structure Information

Given that the structure-based models that only consider the structure information of a protein might not well represent the protein’s multimodal information, namely the sequence and structure information, we hypothesized that introducing the multimodal information of protein with a rational strategy in the CPA prediction model may further improve its CPA prediction performance.

To test this hypothesis, in an end-to-end neural network architecture, we first developed a method to represent the protein structure information (including the Euclidean distances between the residues of proteins in 3D space, the dihedral angles (Φ and ψ) on the backbones of proteins. Then we co-evolutionally updated this structure information with the residue sequences information of proteins, with the aim to comprehensively and efficiently represent their multimodal information. The general workflow of this model, FeatNN, is depicted in Fig. 1. FeatNN was designed based on a dexterous architecture that can process amino acid sequences and atom sequence with any lengths; thus, the whole set of information about proteins and compounds can be characterized. More specifically, the compound information proceeds through the compound extractor module (Fig. 1a and Supplementary Fig. 13) that consists of a multihead vertex representation (Fig. 1a and Supplementary Fig. 14) and deep GCN blocks (Fig. 1a and Supplementary Fig. 9). Notably, the deep GCN block is applied to prevent the oversmoothing problem during training process [38] of the compound extractor (the oversmoothing problem is described in more detail in Supplementary Note 1.1). To allow the remote atoms to communicate with a certain node, a master node is employed to simultaneously capture both local and global features so that FeatNN can learn comprehensive compound features from both global and local views at the same time.

Meanwhile, for the representation of protein structure information, the distance matrix of protein residues is discretized into one dimension, and the strategy of word embedding is applied to encode structure information regarding the Euclidean distances between protein residues as a discrete distance matrix (DDM), which greatly reduces the computational cost of obtaining structure information while still allowing the model to effectively represent the structure information of proteins. After that, the protein features are generally learned by the protein extractor module (Fig. 1b and Supplementary Fig. 15). In the protein extractor module, a Prot-Aggregation block (Fig. 1b and Supplementary Fig. 17) first converts the residue sequence of the given protein, the DDM, and the torsion matrix into two variables: a new matrix representing the residue sequence of the protein and a new distance matrix encoded with the structure information of the protein. The two outputs generated from the Prot-Aggregation block are then fed into the Evo-Updating block (Fig. 1b and Supplementary Fig. 18), which serves as the vital component in the protein encoder module (Fig. 1b and Supplementary Fig. 16). In this way, the structure and sequence information are interactively aggregated through a coevolutionary strategy in the Evo-Updating block, which ensures that FeatNN can learn preeminent features from multimodal protein information.

Finally, the learned representations of compound features and protein features are input into the affinity learning module (Fig. 1c and Supplementary Fig. 20). The detailed designs of the compound extraction module, protein extraction module and affinity learning module are described in the Methods and Supplementary sections.

### FeatNN Outperformed the SOTA Model in CPA Prediction

To assess the performance of FeatNN, seven kinds of models mentioned above were trained on the dataset generated from the general PDBbind set, and their CPA prediction performances were compared (Fig. 2 and Supplementary Fig. 3). In addition to our model (FeatNN), the baseline models were BACPI [37], SIGN [26], MONN [9] and four variants of GraphDTA (i.e., GATGCN, GCNNet, GATNet and GINConvNet)[36]. Because some compounds and proteins tend to be highly similar and homologous, we followed the clustering strategy (for details, see Supplementary Methods 3.4 and 3.5) proposed in previous studies to prevent information leakage from the test set data during the model training process [9, 39]. Four different clustering thresholds were used to split and cluster the similarity data into training, valid and test sets in the control group experiment. They were 0.3, 0.4, 0.5 and 0.6, indicating the minimum distance between each similar class. For example, a 0.3 clustering threshold meant that any compounds from two different sets (training, valid, or test set) were at least 30% different in terms of their respective structures. In terms of the compound-clustered test group, FeatNN^general^ outperformed the SOTA baseline^general^ (MONN) by 21.33% in the R^2^ metric under IC_50_ (Fig. 2b and Supplementary Table 3) and 17.07% under KIKD (Fig. 2a and Supplementary Table 3). In addition, the evaluation results of the protein-clustered test group can be found in Supplementary Fig. 3. FeatNN^general^ also surpassed the baseline models in most cases (Supplementary Fig. 3 and Supplementary Table 3). However, as shown by Supplementary Fig. 3a, the SIGN model achieved the best performance in RMSE but the worst in Pearson and R^2^ on the “KIKD” dataset constructed from the general set of PDBbind-v2020, possibly because the SIGN model efficiently learned the absolute error (RMSE) between the prediction affinity and the real ones, but unable to learn their correlation (Pearson, R^2^). Even though the similarity of the data (protein or compound) in the same dataset (training, validation, or test datasets) decreases with increasing threshold, the CPA prediction correlation performances of FeatNN^general^ remained consistent and it outperformed the baselines, indicating the robustness and outstanding performance of FeatNN in comparison with the baseline models. Furthermore, we trained FeatNN^refine^ on the refined datasets of PDBbind [31] to assess whether a high-quality structural dataset can enhance its CPA prediction performances. Interestingly, we found that the Pearson performances of FeatNN^refine^ and SOTA baseline^refine^ were respectively elevated by 2.65% and 5.45% compared to the corresponding methods trained on general datasets of PDBBind with the compound-clustered method (with the threshold of 0.3, details in Supplementary Fig. 4a, Supplementary Fig. 5a, Supplementary Table. 4, Supplementary Table 5). However, R^2^ and Pearson values of FeatNN^refine^ and SOTA baseline^refine^ were found to be somewhat lower when applying the protein-clustered method, indicating that the accuracy and generalization of models were affected, possibly due to the limited number of high-quality data in the refined dataset of PDBbind-v2020 (Supplementary Fig. 4b, Supplementary Fig. 5b, Supplementary Table. 4, Supplementary Table 5). According to the statistic result (Supplementary Table 1), we found the protein diversity is poor in the refined dataset. Such a negative effect is observed possibly because the diversification of protein data is crucial for the performance of a computational model in CPA prediction tasks [40].

### Performances of FeatNN on the BindingDB Dataset

Even though the PDBbind database has rich protein diversity, the amount of paired information in this database is limited (12,699 records). By contrast, the BindingDB database is much larger (218,615 records), but the quality of the structural data in this database is not very high, and it is also poor in protein diversity and provides limited structure information for the compound and protein complexes. To comprehensively evaluate the performances of FeatNN, we first tested FeatNN and baseline models on BindingDB with a large-scale compound-protein interaction dataset. To do so, on the dataset generated from BindingDB with 218,615 compound-protein pairs, FeatNN and the baseline models were evaluated with 153,031 training samples, 21861 validation samples and 43,723 test samples[3]. To conduct a fair comparison, we evaluated the CPA prediction performance of the models by averaging the prediction results obtained over approximately 10 independent training processes on the dataset generated from BindingDB database. In contrast to the computer vision and natural language processing fields, the data in the biotechnology field are more flexible. The diversity of data in different datasets and the composition of data pairs may greatly change the performance of the model. As shown in Table 1, FeatNN outperformed the SOTA baseline with the best RMSE (0.765), Pearson correlation coefficient (0.850) and R^2^ value (0.719).

### Applying Pretraining Strategy Enhanced the Performances of FeatNN

First, to assess the generalization ability of FeatNN (Details in Supplementary Methods 3.6), we set up an independent third database named Binding MOAD with high-quality paired information data (the details for the generation of this dataset are provided in Supplementary Table 1). As shown in Supplementary Fig. 6, we found that the generalization ability of FeatNN was strongly depended on the amount of paired information in the training datasets. When trained on the general PDBbind dataset, FeatNN^general^ showed superior generalization performance, outperforming the SOTA baseline^general^ by 4.57% and 5.72% for the evaluation of the Pearson coefficient tested on IC_50_ and KIKD measurement datasets constructed from Binding MOAD (Supplementary Fig. 6, Supplementary Fig. 7, Supplementary Table 6, Supplementary Table 8,). However, when trained on the refined datasets of PDBbind even with higher data quality, the models (both FeatNN^refine^ and the SOTA baseline^refine^) trained on the refined dataset of PDBbind showed considerably lower generalization ability compared to the corresponding models (FeatNN^general^ and the SOTA baseline^general^) trained on the general PDBbind dataset (Supplementary Fig. 6, Supplementary Table 6), with decreases by 62.95% and 93.10% in R^2^ evaluation for FeatNN and SOTA baseline, respectively, possibly due to the limited amount of paired information used in the training process.

To further enhance the performance of FeatNN, FeatNN^optm^ was tentatively trained by applying a pretraining strategy [41] to warm FeatNN up on the dataset with relatively low-quality structure data generated from BindingDB (Fig. 3a, Supplementary Methods 3.7). Considering that CPA prediction on PDBbind and BindingDB served as the same type of task, the parameters of the compound extractor learned from the two datasets could be highly generalized and portable. To test this hypothesis, we attempted to assess whether the performance of FeatNN on the PDBbind dataset could be improved by this parameter transfer strategy. To do so, the compound extractor parameters learned from BindingDB were frozen at first. The next steps were to fine-tune the protein extractor and affinity learning module, take the ‘knowledge’ learned from BindingDB as the initial parameters of the protein extractor and affinity learning module. In this way, we fine-tuned these two modules on the datasets generated from PDBbind, that is, to conduct multiple rounds of training and thus obtain FeatNN^optm^ (Fig. 3a). As a result, the RMSE, Pearson coefficient, and R^2^ of FeatNN^optm^ for the PDBBind test dataset were increased by 3.29%, 1.93% and 5.47% (Fig. 3b and Supplementary Table 7) respectively, suggesting the excellent transferability of FeatNN to different datasets. Interestingly, the generalization ability of FeatNN^optm^ is further enhanced by 2.04% and 5.79% for Pearson and R^2^ compared with FeatNN directly trained on the PDBbind (Supplementary Fig. 7, Supplementary Table 8).

**Fig. 3.**
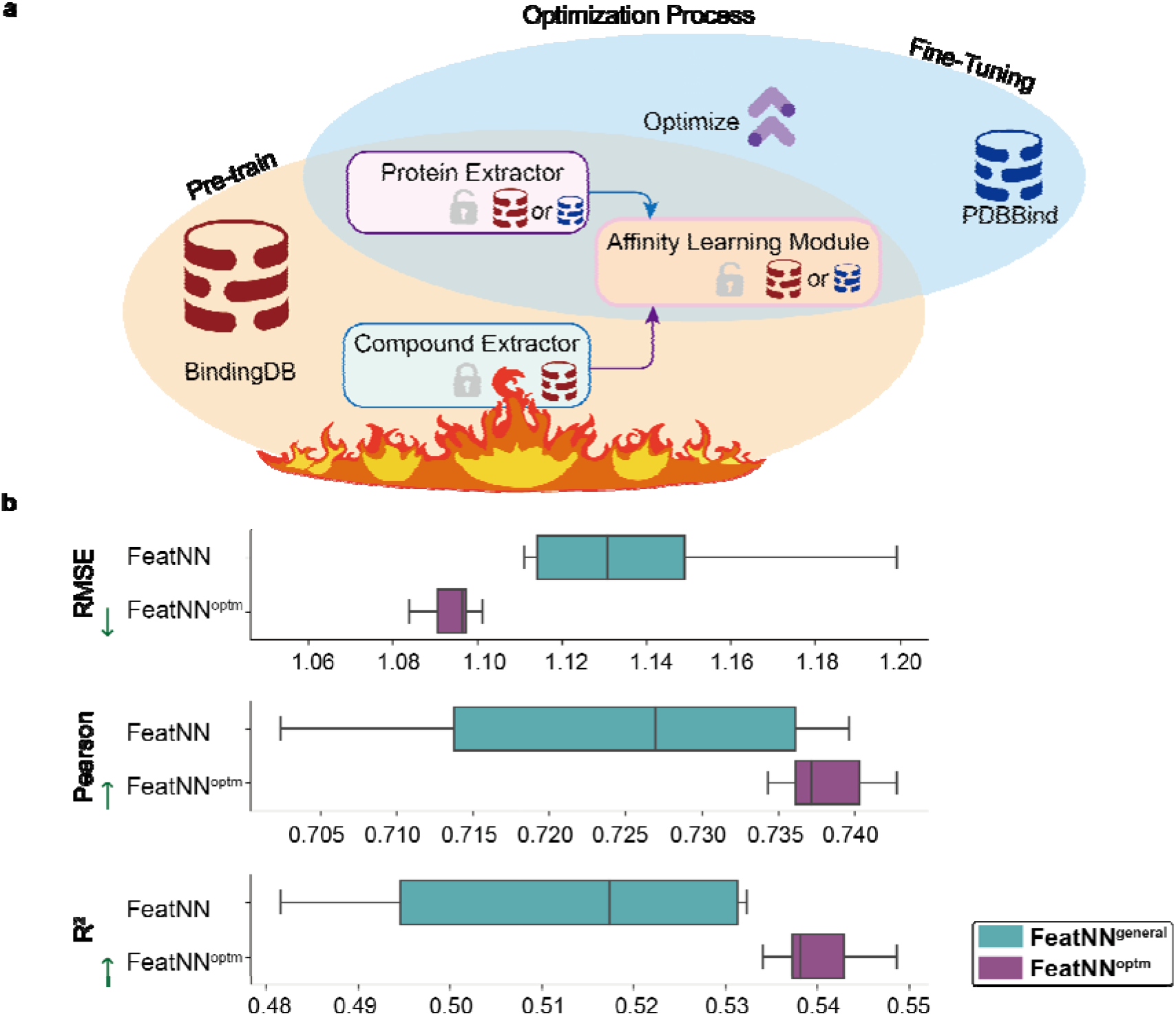
The performance of FeatNN is greatly improved after optimization with fine-tuning strategy. **a**. To optimize the performance of FeatNN, the parameters of the compound extractor obtained from the warm-up (pretraining) strategy on BindingDB are frozen, and then the protein extractor module and affinity learning module are fine-tuned on PDBbind to obtain FeatNN^optm^. **b.** The RMSE, Pearson coefficient, and R^2^ of FeatNN with the fine-tuning strategy (FeatNN^optm^) were increased by 3.29%, 1.93% and 5.47% compared with that of the FeatNN version directly trained on PDBbind-v2020. FeatNN: original FeatNN trained on PDBbind. FeatNN^optm^: FeatNN optimized with a fine-tuning strategy. The results of each group were counted with 10 independent experiments by 5-fold cross-validation strategy. The mean value, upper and lower quartiles, and SD of each independent experiment group are clearly shown in Fig. 3b. Box plots; boxes depict the upper and lower quartiles of the data, and the vertical line in the box indicates the median of the statistical value of the group.

### The Functionality-based Interpretation of the FeatNN Module

To elucidate the function of each block in FeatNN, we sought to assess the performance of FeatNN by ablating the blocks (Supplementary Methods 3.8) that were specifically designed to elevate its performance (for details, see Methods). The results shown in Fig. 4 demonstrate that a variety of components contribute significantly to the accuracy of FeatNN in CPA prediction. For instance, the robustness and prediction accuracy of FeatNN declined by approximately 14.34% in terms of the RMSE, 11.60% in the Pearson coefficient and 31.25% in R^2^ without Evo-Updating, emphasizing the significance of the coevolutionary strategy in the protein extractor. Strikingly, the prediction accuracy decreased by approximately 15.22% in the RMSE, 15.61% in the Pearson coefficient and 36.33% in R^2^ without addressing the oversmoothing problem via the deep GCN block. In addition, the master node in the deep GCN block, which represented the global information of each compound and communicated with the remote graph node through the graph warp unit (Fig. 4 and Supplementary Table 9), also contributed significantly to the accuracy of CPA prediction, highlighting the importance of interactively updating the global and local features and the importance of addressing the oversmoothing problem when representing the information of compounds. More importantly, the performance of the FeatNN versions that only used protein sequence information or structure information (DDM and torsion matrix) declined markedly by approximately 36.52% and 69.34%, respectively, in R^2^ compared with the intact FeatNN baseline (Fig. 4 and Supplementary Table 9), emphasizing the importance of introducing the coevolutionary strategy to jointly aggregate and update the sequence and structure information of proteins. We ablated the compound-protein interactive matrix in the affinity learning module, which could help FeatNN to represent and learn the interaction information between compound and protein, and found that the R^2^ performance declined by 38.09% (Fig. 4), indicating the rationality of learning effective interaction features by compound-protein interactive matrix. In addition, we ablated the torsion-related architecture and found that the performances declined by 13.48% in R^2^ (Fig. 4 and Supplementary Table 9), highlighting the necessity of introducing the torsion information into FeatNN.

**Fig. 4.**
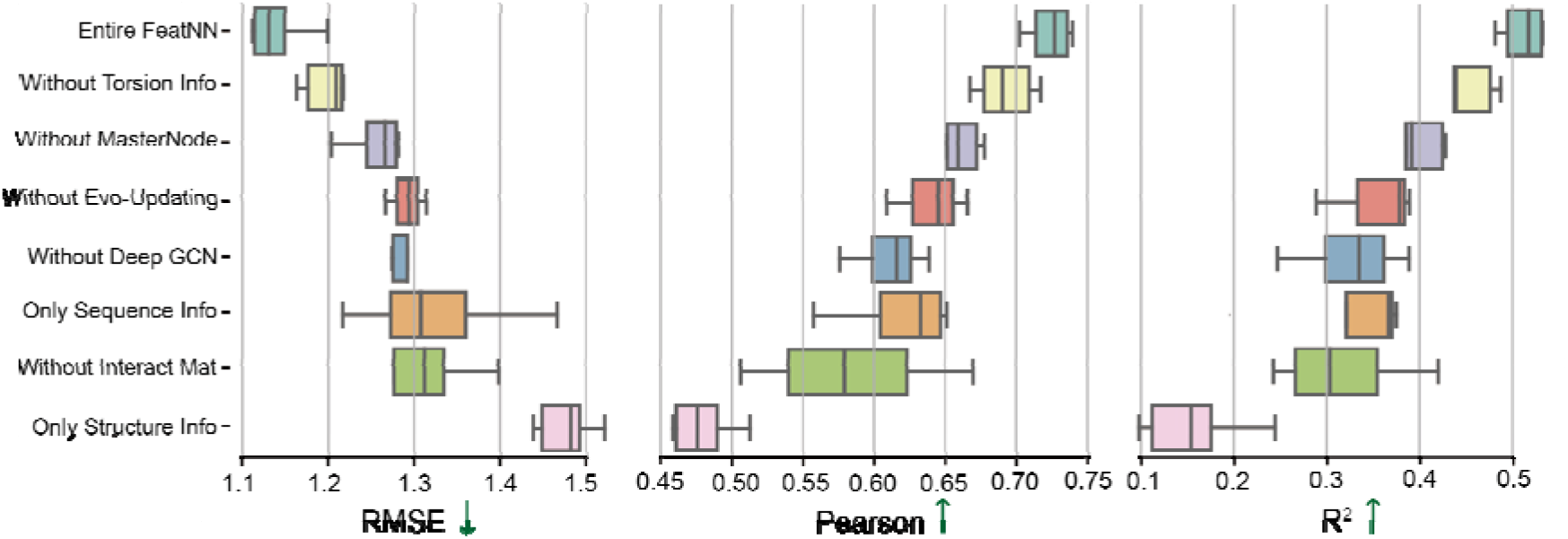
Essential block ablation results of FeatNN. Ablation results of FeatNN on the dataset generated from PDBbind, emphasizing the functionality of the essential blocks of FeatNN. The accuracy and robustness of FeatNN in terms of CPA prediction dramatically decline without the Evo-Updating block or torsion information, which functions as the core in protein feature extraction. Addressing the oversmoothing problem in the deep GCN block also remarkably increases the ability of the compound extractor to extract features from compounds, which in turn enhances the CPA prediction accuracy of the overall model. In addition, introducing the master node into the network to learn the global information of compounds is also important. The performances of the FeatNN version that only uses protein sequence information or structure information also remarkably decline compared with the entire FeatNN baseline, suggesting the importance of applying the coevolutionary strategy to interactively represent and update features of both sequence and 3D protein structure information. Furthermore, with ablation of the compound-protein interactive matrix, significant decline is observed in performances of the FeatNN, indicating the importance of learning the interaction features between protein and compound. The results of each group were counted with 10 independent experiments by 5-fold cross-validation strategy. The mean value, upper and lower quartiles, and SD of each independent experimental group are clearly depicted in Fig. 4. Box plots; boxes depict the upper and lower quartiles of the data, and the vertical line in the box indicates the median of the statistical value of the group. Abbreviations: Info: information.

### The Interpretation of Information Flows in FeatNN

To understand how information flows in the deep GCN, Evo-Updating, and affinity learning module, we visualized the original features in the intermediate layers of FeatNN (Supplementary Fig. 8). Because it is difficult to show the information transformation process in the original features directly, we applied t-distributed stochastic neighbor embedding (t-SNE) [42], a compression algorithm for high-dimensional data, to obtain a limpid data distribution in two dimension view (Fig. 5). As shown in Fig. 5, the atom features became more aggregated as the GCN layers deepened. This phenomenon dynamically explained why the node information flows in the layers and aggregates the features of neighbor nodes through the message passing mechanism [43] in the deep GCN block (Fig. 5a). In the Evo-Updating block, embedded sequence features and structure features were obtained from the Prot-Aggregation block, and then the sequence features and structure features were partially updated on each other, and part of their own information was integrated into the Evo-Updating block (Fig. 5b). When the Evo-Updating layers deepened, the difference between the sequence features and structural features gradually lessened, and the layers fused more multimodal information into themselves. Additionally, we extract the compound and protein features, which are learned from the deep GCN block and Evo-Updating block, respectively, in each layer for dimension reduction analysis (Fig. 5c, 5d). The distributions of compound features learned in the deep GCN block of each layer are clearly illustrated (Fig. 5c, 5d). We found that the features aggregated by the first three layers of the block have a certain degree of similarity, whereas the distribution of compound features tends to be more distinguishable in deep layers of GCN block (Fig. 5c), which might enable FeatNN to learn the precise features of the compound and address the notorious oversmoothing problem (Fig. 4 and Supplementary Fig. 11a-c). In the Evo-Updating block, we showed that the eigenspace distance between protein structural features and sequence features that are learned in the same layer remains adjacent (Fig. 5d). More interestingly, we found that both the sequence and structural features learned in the deep layer of the block are updated along the same direction (evolution) through this coevolutionary strategy, which efficiently represents the multimodal information of proteins and ultimately benefits the CPA prediction accuracy (Fig. 4).

**Fig. 5.**
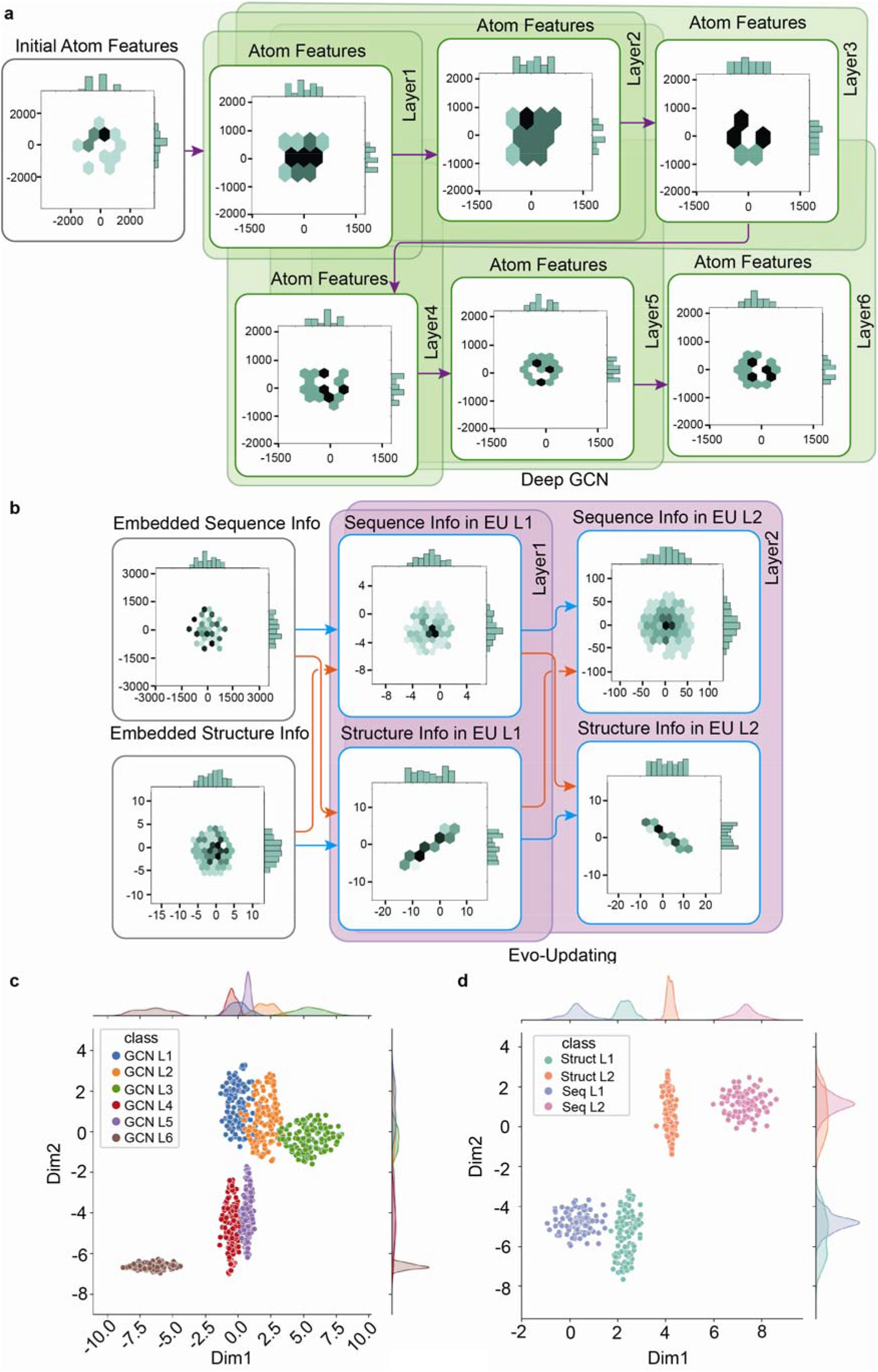
Information flows in FeatNN’s deep GCN and Evo-Updating blocks. **a**. Visualization of the compound information aggregation process in the deep GCN block. **b**. Visualization of the coevolutionary process between the protein sequence and structure information in the Evo-Updating block. **c**. t-SNE dimensionality reduction analysis of deep GCN block (6 layers). **d**. t-SNE dimensionality reduction analysis of Evo-Updating block (2 layers). Abbreviations: EU L1 or L2: Evo-Updating Layer1 or Layer2. GCN L1 or L2: GCN block Layer1 or Layer2. Struct L1 or L2: Structure features in EU L1 or L2. Seq L1 or L2: Sequence features in EU L1 or L2. Embedded Sequence Info: sequence features obtained from the Prot-Aggregation block. Embedded Structure Info: structure features obtained from the Prot-Aggregation block. Initial atom features: atom features obtained from the graph embedding.

### FeatNN Outperformed the SOTA Baseline in Virtual Drug Screening Tasks

To verify the feasibility of the use of FeatNN in virtual drug screening tasks [37, 44], we initially selected “SARS-CoV-2 3C-like protease” as the drug target (receptor), which is a verified target for developing drugs to cure SARS-CoV-2 [45]. We unbiasedly selected 28 bioactive small molecules [45–59] (listed in Supplementary Tab. 10, note: these molecules related to the target did not exist in PDBbind nor BindingDB) from publication research and the DrugBank database. The process of receptor-based affinity value prediction by applying FeatNN is shown in Fig. 6a. In addition, we selected a ligand-free protein structure of SARS-CoV-2 3C-like protease with the identity number of 7CWC in PDB. Strikingly, we found that the Pearson coefficient reached a value of 0.612 (Fig. 6b) in a CPA prediction task. Compared with the SOTA baseline (MONN) that obtained a Pearson coefficient of 0.402 (Fig. 6c), this was suggestive of the outstanding performance of FeatNN in searching for potential drug candidates from a massive database.

**Fig. 6.**
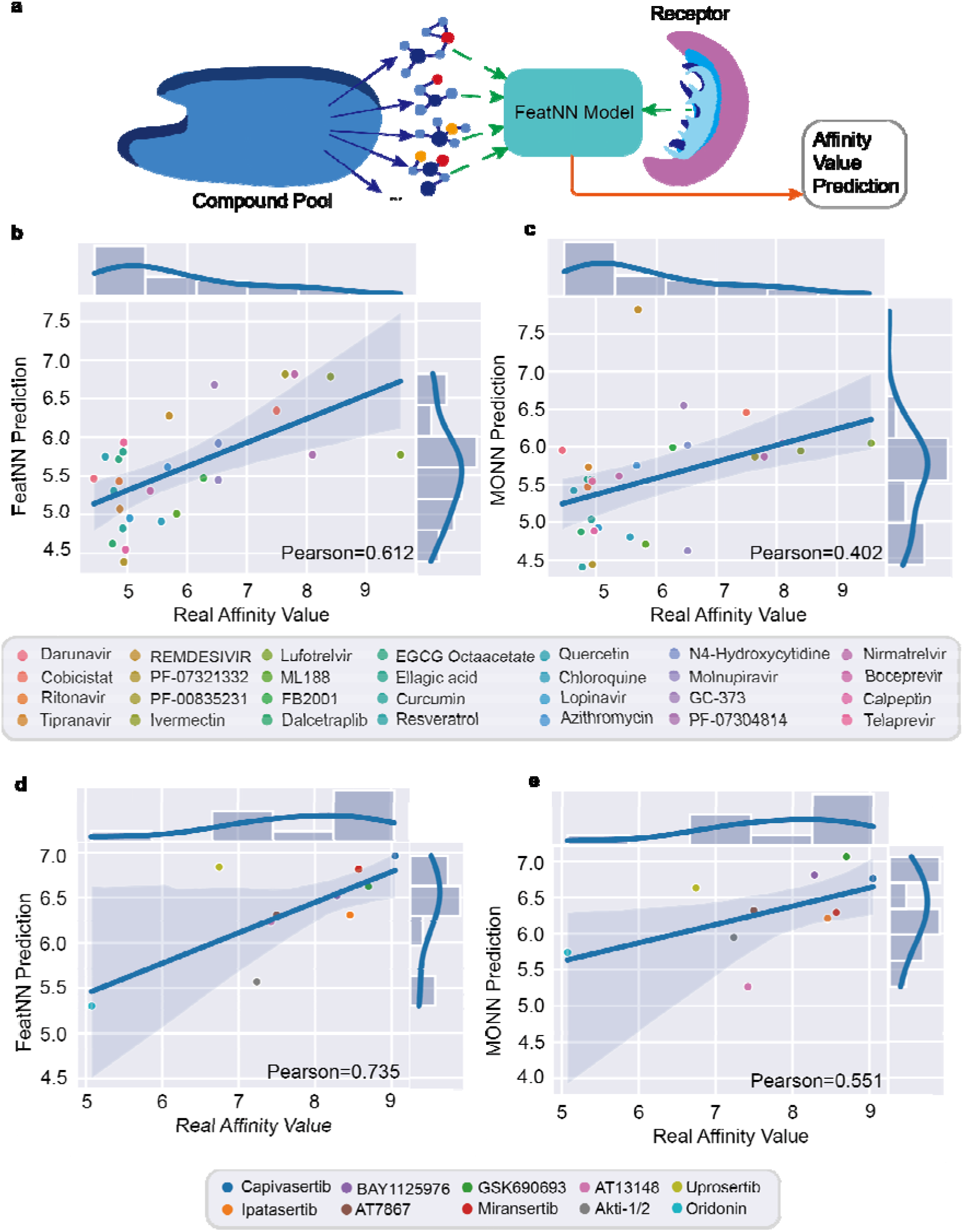
Affinity prediction results of FeatNN and the SOTA baseline in practice. **a**. Receptor-based virtual screening tasks: targeting both receptors of the SARS-CoV-2 3C-like protease and Akt-1, related bioactive compounds were unbiasedly selected (Supplementary Table 10 and 11) from published research and the DrugBank database to test the affinity prediction precision and generalization ability of FeatNN. Targeting 3CL protease, **b**. the affinity prediction of 28 validated bioactive compounds by FeatNN result in a Pearson coefficient of 0.612. **c**. The affinity prediction of 28 validated bioactive compounds by MONN result in a Pearson coefficient of 0.402. Targeting Akt-1, **d**. the affinity prediction of 10 validated bioactive compounds by FeatNN results in a Pearson coefficient of 0.735. **e**. The affinity prediction of 10 validated bioactive compounds by MONN results in a Pearson coefficient of 0.551. Note: From the above experiments, it can be seen that MONN serves as the SOTA baseline in both datasets that generated from PDBbind and BindingDB databases, which is the reason that we only used MONN as a representative baseline model for testing. Both structure conformations of 3CL protease and Akt-1 are extracted from the PDB file with the PDB id of 7CWC and 3O96. Each point was obtained by the average of 15 independent experiments.

In addition, to verify the robustness of FeatNN, we repeated the prediction task many times and analyzed the results statistically (Fig. 6b). Nonetheless, a concern remained regarding the multimodality-based model of FeatNN: the prediction results obtained with different 3D protein structure conformations might have been variable. To assess this possibility, we selected the ligand-free protein conformations from 3 PDB files (recorded with PDB-ids of 7CWC, 7CWB and 7BAJ in the PDB Database, Supplementary Fig. 9a) of SARS-CoV-2 3C-like proteases as receptors for CPA prediction with FeatNN (Fig. 6b and Supplementary Figs. 9b-c). Remarkably, the CPA prediction task among 28 validated compounds still achieved robustness and exhibited excellent results with Pearson coefficients of 0.606 and 0.607, indicating that the prediction results obtained with FeatNN do not exhibit unstable changes in different target conformations (Fig. 6b and Supplementary Figs. 9b-c). To verify the feasibility of the use of FeatNN on different targets, we additionally chose a target named Akt-1 (PDB-id: 3O96) that is a critical receptor for the transmission of growth-promoting signals and resisting cancer [51]. In this experiment, 10 previously reported drugs (Supplementary Table. 11) that target Akt-1 [60–69] were selected for this virtual screening task, and FeatNN showed a better Pearson performance of 0.735 in the CPA prediction task compared with the SOTA baseline. Using different Akt-1 conformations (PDB-ids of 6HHJ, 3MV5, 3CQW and the ligand-free conformation predicted by AlphaFold2 [28]), the Pearson performance also remained stable (Fig. 6d, Supplementary Fig. 10 and Supplementary Table 11), indicating the robustness and reliable prediction ability of FeatNN in various virtual screening tasks with different targets.

## Discussion

The FeatNN model proposed in this study introduced a coevolutionary strategy to effectively represent multimodal protein features. Through a t-SNE visualization analysis and a module ablation study, from the perspective of interpretation, we showed that the information between protein sequences and structure features was jointly updated and aggregated, which ultimately benefited the CPA prediction accuracy of our approach. In this study, we found that the Evo-Updating block and deep GCN block in FeatNN function as the key components for aggregating and updating the features of both proteins and compounds (Fig. 4), emphasizing the significance of applying the coevolutionary strategy in protein feature extraction. Altogether, FeatNN learns efficiently from a limited data resource but is still able to cope with the complexity of structure data and achieve outstanding performance.

Although it is theoretically appealing to introduce the structural information of proteins in a CPA prediction model, we overcame numerous obstacles in the development of FeatNN. First, we elegantly overcame the oversmoothing problem[38] by introducing a specific residual connection in each layer of the GCN, which could add part of the initial information of the molecular graph into the current layers [70, 71]; therefore, the extraction ability of the model with respect to compound features was enhanced when the layers deepened (Supplementary Fig. 11a-c). Second, in the deep GCN block, a master node was employed to learn the global features during the training process, thus facilitating communication among remote nodes. Third, the protein distance matrix was discretely encoded to overcome the overwhelming information problem of the traditional continuous distance matrix. As a result, FeatNN greatly outperformed the SOTA model in tasks involving generalization ability on an independent database and targeting the “SARS-CoV-2 3C-like protease” and “Akt-1” affinity value prediction, indicating that FeatNN can be a powerful tool for advancing the drug development process.

Nevertheless, due to the scarcity of precise noncovalent interaction binding site data between the ligand and the binding pocket, and the data imbalance problem in the distribution of the few positive and predominantly negative data of binding sites, FeatNN faces difficulties in interpreting the CPA prediction results at the interaction level at current stage. Traditional methods such as upsampling and gradient penalty still cannot address such a dilemma (imbalance problem) without enough data for binding interactions [72]. Possibly, docking simulation combined with AI may be able to interpret the results predicted by the AI models at the interaction level [20], which may be a new research direction in the further development of FeatNN in our future study. Moreover, 3D structural information is not only relevant to proteins but also to other compounds [25, 73]. In this study, we only introduced the protein structural information, and experiments to additionally introduce compound geometry information are ongoing [74]. Theoretically, the strategy developed for protein feature extraction in our model could also be utilized to extract the geometric information of compounds. It could be appealing to introduce both protein and compound structure features in our model to further enhance its performance, given that the application of only the protein structure features in this study has already achieved a remarkable result. Other protein properties, such as the residue types of binding ligands, secondary structures and physicochemical characteristics, are also very important features. Incorporating these features into our model might further improve its performance. However, the challenge is how to represent these features with a rational method or provide an interpretable architecture, which is left to be addressed in future studies.

### Limitations

1) The training of the deep learning model depends strongly on the training data. In practice, if compounds or proteins are encountered with fairly different similarities that are very different from the data in the training set, the confidence in the prediction results will be greatly reduced. 2) Furthermore, because the architecture of FeatNN highly depends on the 3D structure of the protein, some protein data cannot be characterized due to the residue continuity defect of PDB files, so they must be discarded. Therefore, the number of training data will be decreased, but this will not significantly affect the performance of FeatNN. 3) Even though FeatNN can achieve improved precision and generalization ability in CPA prediction while ignoring the information regarding the binding pose between the ligand and the binding pocket, it is difficult for FeatNN to interpret the CPA prediction results at the interaction level, because of the scarcity and data imbalance problems of precise noncovalent interaction data between the ligand and the binding pocket.

## Conclusion

The proposed FeatNN model introduces a torsion matrix and a distance matrix in its protein extractor module, and it utilizes the deep GCN block with the master node in the compound extractor module to predict the affinity of a given compound-protein pair. The experimental results of our study showed that FeatNN outperformed the SOTA baseline by a significant margin, and the accessibility of FeatNN applied in lead compound screening was also verified; this approach demonstrates great potential for reducing the considerable time and expense involved in drug candidate screening experiments, and provides an interpretable architecture based on biology databases.

## Key Points

- We apply both 3D protein structure and sequence information with a coevolutionary strategy.
- We addressed the oversmoothing problem in graph representation of compounds.
- FeatNN achieved highly enhanced affinity prediction on well-known databases compared with the state-of-the-art methods.
- Generalization ability and feasibility of FeatNN are superior to the SOTA baseline both on the datasets generated from the Binding MOAD database and the virtual screening task targeting the receptor of the SARS-CoV-2 3CL protease and Akt-1.

## Acknowledgments

This study was supported by the Scientific and Technological Innovation 2030 Program of China - major projects (2021ZD0200408 to X.W.), the National Natural Science Foundation of China (81971866 to X.W.), the Natural Science Foundation of Zhejiang Province (LR20H090002 to X.W.), the Leading Innovative and Entrepreneur Team Introduction Program of Zhejiang (2019R01007 to X.W.), and the Fundamental Research Funds for the Central Universities (K20210195 to X.W.). We would like to thank professor Jie Yang for his feedback and advice on writing this paper.

## Author Contributions

B.G., H.Z. and H.J. contributed equally to this work. X.W., B.G. and H.Z. conceptualized and designed the study. B.G., H.Z. and H.J. conducted the experiments and collected the data. B.G., H.Z., H.J., X.W., H.Y., X.L., N.G. and Y. Z. analyzed and interpreted the data. B.G., H.Z. and X.W. drafted the paper. All authors critically revised the manuscript and approved the final version for submission.

## Data Availability

The data that support the findings of this study are included in the paper, and further data are available from the corresponding author upon reasonable request.

