## Supplemental Tables and Figures for "Enhanced compound-protein binding affinity prediction by representing protein multimodal information via a coevolutionary strategy"

Affiliations:

### 1. Supplementary Notes

#### 1.1. The Oversmoothing Issue in GCNs

Deep graph convolutional networks (GCNs) have been very popular since 2017, when Kipf and Welling achieved great success by obtaining SOTA performance on a semisupervised classification task[1]. This method can also be used in biological research to represent compound features and optimize compound property predictions[2, 3]. However, this method always encounters an oversmoothing issue due to the limitation of depth[4]. In other words, the performance of the GCN becomes worse when the number of layers increases because the representations of the nodes in the GCN converge to approximately the same values. Applying the residual network (ResNet)[5] and appending residual connections in GCN models can hardly solve this problem, while oversmoothing in a GCN is a type of Laplacian smoothing. To circumvent this issue, inspired by GCNII[6], a specific residual connection with the initial features of each node in the molecular graph is applied to extract compound features in our work; this strategy increases the number of layers from 2 to 4, enabling the model to extract more information. We mathematically interpret the oversmoothing issue in a traditional GCN as follows.

First, we define a simple and connected undirected graph $G$ (Supplementary Fig. 1a) with $n$ nodes and $m$ edges. We use $A$ as the adjacency matrix and $D$ as the degree matrix of graph $G$, where $d(v_{i})$ is the degree of node $v_{i}$. Let $\tilde{A}$ and $\tilde{D}$ be the adjacency and degree matrices of graph G augmented with self-loops. The normalized graph Laplacian matrix is defined as$L = I- \tilde{P}= I- \tilde{D}^{-1/2}\tilde{A}\tilde{D}^{-1/2}$, and time proceeds in unit steps: $t = 1,2,\ldots n$. At each time t, the walk stays at some node $v_{i} \in V$, and at time $t+1$, based on the transition matrix $P$, as $P=AD^{-1}$, the walk randomly chooses one of $v_{i}$’s neighbors to move to (Supplementary Fig. 1b); this is described as a random walk. A lazy random walk is a modified version of the original random walk. In a lazy random walk, at time t, the walker stays at the current vertex with the probability of $\frac{1}{2}$ and takes a step as in the original random walk with the probability of $1/2$ (Supplementary Fig. 1c).

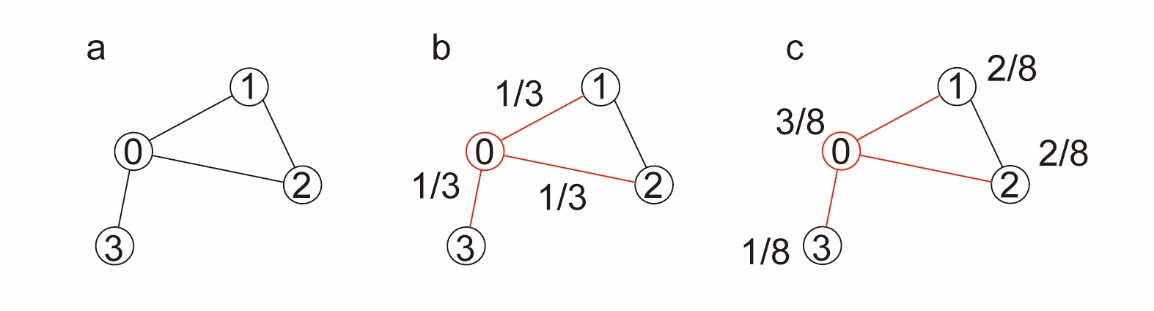

**Supplementary Fig. 1 Graph representation.** Figs. 1a-c. represent three iterations in a graph.

We define a probability vector $\pi$ that corresponds to the stationary distribution of the random walk. At time $t , \pi^{t+1}=P\cdot\pi^{t}= AD^{-1}\cdot\pi^{t}$, and $\pi\left( v_{i} \right)=\frac{d_{v_{i}}}{2m}$. This breaks the periodicity of the random walk and forgets the initial graph information.

Because a deep GCN faces the oversmoothing problem, we first consider a multilayer GCN:

$$H^{(l+1)}=\tilde{P}\cdots\sigma(\tilde{P}\sigma\left( \tilde{P}XW^{\left( 0 \right)} \right)W^{\left( 1 \right)})\cdots W^{\left( l \right)}$$

$W^{\left( l \right)}$is a layer-specific trainable weight matrix, $H^{(l)}$is the matrix of activations in the $l$th layer, $H(0) = X$, and $\sigma\left( \cdot\right)$ denotes an activation function. First, ignoring $\sigma\left( \cdot\right)$, we can describe the matrix as $H^{(K)}=\tilde{P}^{K}XW$,$\tilde{P}= \tilde{D}^{-1/2}\tilde{A}\tilde{D}^{-1/2}$, and then expand the calculation; we obtain

$$\tilde{P}^{K}= \tilde{D}^{-1/2}\tilde{A}\tilde{D}^{-1}\tilde{A}\tilde{D}^{-1}\cdots\tilde{A}\tilde{D}^{-1}\tilde{A}\tilde{D}^{-1/2}$$

$$= \tilde{D}^{-1/2}(\tilde{A}\tilde{D}^{-1})(\tilde{A}\tilde{D}^{-1})\cdots(\tilde{A}\tilde{D}^{-1})\tilde{A}\tilde{D}^{-1/2}\cdot\tilde{D}^{-1/2}\cdot\tilde{D}^{1/2}$$

$$= \tilde{D}^{-1/2}{(\tilde{A}\tilde{D}^{-1})}^{K}\tilde{D}^{1/2}$$

This demonstrates that as the number of layers increases, the nodes in the GCN converge to certain values; this convergence makes the initial information indistinguishable, degrading the performance of the GCN.

**Supplementary Table 1** Overall statistics of the datasets extracted from PDBbind, BindingDB and Binding MOAD. The refined set of PDBbind only contains the measurements with K_i_ and K_d_. BindingDB is rich in measured IC_50_ values (more than 500 thousand data, due to the defects of the PDB file and some compounds could not apply graph representation (atoms with more than 6 adjacent nodes), we only obtained 218,615 compound-protein paired data, while the collections of the measured values obtained based on K_i_ and K_d_ are significantly smaller (40 thousand K_i_ measurements and 28 thousand K_d_ measurements are recorded). In this paper, to construct large datasets from BindingDB, we only select the measured IC_50_ values to generate training data. Note: Although the data of compound-protein pairs are fairly rich in BindingDB, the diversity of proteins remains very low compared with the data in PDBbind [7]. To test the generalization ability of the models, we constructed new datasets from the Binding MOAD database and excluded the complexes that appeared in the datasets (training, validation, and test datasets) constructed from PDBbind. For a fair comparison of the generalization ability, we limit the datasets constructed from Binding MOAD with the measurement of IC_50_ and KIKD to the same number of compounds. Thus, we constructed the dataset using the results of the IC_50_ and KIKD measurements from the “all of Binding MOAD” and “nonredundant MOAD” sets in the Binding MOAD database.

| **Database** | **Measurement** | **Quantity** | **Compound**  **Amount** | **PDB**  **Entries** | **Max Affinity** | **Min Affinity** |
| --- | --- | --- | --- | --- | --- | --- |
| **PDBbind**  **(general)** | **KIKD** | 7,156 | 5,691 | 7,156 | 15.2218 | 0.3979 |
|  | **IC_50_** | 5,543 | 5,243 | 5,543 | 11.5229 | 0.4498 |
| **PDBbind**  **(refined)** | **KIKD** | 2768 | 2475 | 2768 | 11.9208 | 2.0 |
| **BindingDB** | **IC_50_** | 218,615 | 183,584 | 2,248 | 11.0458 | 2.3468 |
| **Binding MOAD** | **IC_50_** | 1963 | 1862 | 1952 | 11.5003 | 0.4226 |
|  | **KIKD** | 1915 | 1285 | 1884 | 13.9586 | 0.0773 |

**Supplementary Table 2** Parameter settings of FeatNN training on the datasets generated from the PDBbind (both refined and general sets) and BindingDB databases. Note: FeatNN^optm^ follows the same settings.

| **parameter name** | **value** |
| --- | --- |
| Hidden size (in the entire architecture) | 128 |
| Dropout probability | 0.1 |
| Number of attention heads in the deep GCN block | 4 |
| Number of attention heads in the Evo-Updating block | 4 |
| Layers of deep GCN blocks | 6 |
| Layers of Evo-Updating blocks | 2 |
| α in the deep GCN block | 0.2 |
| λ in the deep GCN block | 0.5 |
| Maximum number of neighbors for each atom node | 6 |
| DDM word embedding size | 40 |
| Torsion size (both the sine and cosine values of Φ and ψ on the backbone) | 4 |
| Kernel size in all CNN layers | 11 |
| Padding size in all CNN layers | 5 |
| Stride in all CNN layers | 1 |

**Supplementary Table 3** Model performance comparisons for the compound-clustered group and protein-clustered group. The models are ordered by their performance on the compound-clustered test group in terms of the R^2^ for IC_50_. FeatNN outperforms the other models by significant margins in all metrics and on both affinity measurements. Each performance result is shown as the mean value and standard deviation (SD) by 5-fold cross-validations with 10 independent experiments. The mean value (and SD) of each independent experimental group is shown in the table.

| **Type** | **Threshold** | **Model** | R^2^ | | **RMSE** | | **Pearson** | | **Spearman** | |
| --- | --- | --- | --- | --- | --- | --- | --- | --- | --- | --- |
|  |  |  | IC_50_ | **KIKD** | IC_50_ | **KIKD** | IC_50_ | **KIKD** | **IC_50_** | **KIKD** |
| Compound-Cluster | 0.3 | FeatNN | 0.512(0.022) | 0.487(0.027) | 1.130(0.045) | 1.442(0.046) | 0.724(0.015) | 0.716(0.015) | 0.697(0.018) | 0.714(0.019) |
|  |  | MONN | 0.422(0.033) | 0.416(0.033) | 1.215(0.015) | 1.485(0.054) | 0.682(0.019) | 0.679(0.024) | 0.661(0.02) | 0.679(0.031) |
|  |  | BACPI | 0.318(0.029) | 0.381(0.043) | 1.289(0.027) | 1.507(0.051) | 0.614(0.010) | 0.633(0.024) | 0.592(0.007) | 0.636(0.025) |
|  |  | GATNet | 0.011(0.031) | 0.182(0.032) | 1.986(0.031) | 1.764(0.034) | 0.182(0.044) | 0.441(0.026) | 0.182(0.038) | 0.385(0.03) |
|  |  | GATGCN | 0.139(0.040) | 0.248(0.027) | 1.853(0.044) | 1.692(0.030) | 0.401(0.040) | 0.511(0.021) | 0.582(0.024) | 0.480(0.029) |
|  |  | GCNNet | 0.124(0.055) | 0.193(0.043) | 1.869(0.058) | 1.752(0.047) | 0.374(0.057) | 0.467(0.026) | 0.361(0.054) | 0.429(0.022) |
|  |  | GINConvNet | 0.164(0.059) | 0.216(0.036) | 1.825(0.064) | 1.727(0.040) | 0.480(0.063) | 0.488(0.026) | 0.517(0.071) | 0.483(0.022) |
|  |  | SIGN | -0.108(0.067) | -0.05(0.032) | 1.366(0.042) | 1.493(0.023) | 0(0) | 0.167(0) | 0(0) | 0.189(0) |
|  | 0.4 | FeatNN | 0.442(0.031) | 0.406(0.073) | 1.202(0.047) | 1.540(0.079) | 0.684(0.018) | 0.669(0.040) | 0.669(0.02) | 0.677(0.041) |
|  |  | MONN | 0.385(0.014) | 0.369(0.057) | 1.251(0.025) | 1.534(0.064) | 0.655(0.010) | 0.643(0.030) | 0.631(0.008) | 0.641(0.039) |
|  |  | BACPI | 0.365(0.020) | 0.251(0.020) | 1.378(0.022) | 1.667(0.022) | 0.632(0.009) | 0.575(0.008) | 0.610(0.006) | 0.600(0.006) |
|  |  | GATNet | -0.003(0.039) | 0.182(0.016) | 2.000(0.039) | 1.764(0.017) | 0.173(0.051) | 0.439(0.019) | 0.173(0.053) | 0.390(0.026) |
|  |  | GATGCN | 0.137(0.047) | 0.235(0.015) | 1.855(0.051) | 1.706(0.017) | 0.397(0.040) | 0.503(0.012) | 0.379(0.043) | 0.473(0.015) |
|  |  | GCNNet | 0.081(0.019) | 0.224(0.021) | 1.915(0.019) | 1.718(0.023) | 0.327(0.034) | 0.491(0.021) | 0.313(0.038) | 0.457(0.029) |
|  |  | GINConvNet | 0.231(0.036) | 0.213(0.063) | 1.752(0.040) | 1.729(0.069) | 0.522(0.010) | 0.490(0.037) | 0.560(0.006) | 0.497(0.024) |
|  |  | SIGN | -0.064(0.09) | -0.005(0.003) | 1.37(0.057) | 1.575(0.07) | 0.089(0) | 0.036(0.033) | 0.103(0) | 0.042(0.038) |
|  | 0.5 | FeatNN | 0.365(0.039) | 0.438(0.036) | 1.281(0.039) | 1.507(0.056) | 0.636(0.028) | 0.685(0.016) | 0.608(0.027) | 0.674(0.023) |
|  |  | MONN | 0.331(0.045) | 0.299(0.061) | 1.306(0.045) | 1.624(0.109) | 0.626(0.028) | 0.611(0.051) | 0.603(0.024) | 0.605(0.054) |
|  |  | BACPI | 0.276(0.017) | 0.264(0.048) | 1.372(0.016) | 1.768(0.056) | 0.563(0.007) | 0.542(0.027) | 0.533(0.005) | 0.514(0.024) |
|  |  | GATNet | 0.013(0.023) | 0.161(0.021) | 1.984(0.023) | 1.786(0.022) | 0.188(0.036) | 0.425(0.011) | 0.194(0.034) | 0.366(0.017) |
|  |  | GATGCN | 0.159(0.030) | 0.239(0.029) | 1.831(0.033) | 1.701(0.032) | 0.424(0.031) | 0.504(0.022) | 0.409(0.021) | 0.467(0.029) |
|  |  | GCNNet | 0.083(0.060) | 0.212(0.008) | 1.912(0.062) | 1.731(0.009) | 0.327(0.062) | 0.479(0.007) | 0.315(0.061) | 0.438(0.012) |
|  |  | GINConvNet | 0.221(0.038) | 0.231(0.026) | 1.763(0.042) | 1.710(0.029) | 0.507(0.051) | 0.500(0.022) | 0.540(0.057) | 0.499(0.016) |
|  |  | SIGN | -0.069(0.034) | -0.077(0.081) | 1.484(0.024) | 1.543(0.084) | 0(0) | 0.060(0.171) | 0(0) | 0.066(0.208) |
|  | 0.6 | FeatNN | 0.339(0.023) | 0.398(0.043) | 1.295(0.053) | 1.429(0.088) | 0.613(0.021) | 0.660(0.028) | 0.59(0.022) | 0.634(0.022) |
|  |  | MONN | 0.210(0.093) | 0.248(0.057) | 1.399(0.092) | 1.681(0.155) | 0.572(0.030) | 0.559(0.057) | 0.554(0.029) | 0.544(0.032) |
|  |  | BACPI | 0.132(0.033) | 0.278(0.024) | 1.569(0.030) | 1.629(0.027) | 0.482(0.009) | 0.582(0.011) | 0.467(0.005) | 0.561(0.009) |
|  |  | GATNet | 0(0.035) | 0.167(0.034) | 1.998(0.035) | 1.780(0.036) | 0.181(0.035) | 0.423(0.033) | 0.176(0.038) | 0.369(0.05) |
|  |  | GATGCN | 0.141(0.060) | 0.244(0.051) | 1.850(0.064) | 1.695(0.057) | 0.399(0.055) | 0.511(0.036) | 0.382(0.064) | 0.482(0.042) |
|  |  | GCNNet | 0.072(0.054) | 0.210(0.019) | 1.923(0.056) | 1.733(0.021) | 0.302(0.059) | 0.475(0.012) | 0.293(0.06) | 0.433(0.019) |
|  |  | GINConvNet | 0.196(0.042) | 0.198(0.083) | 1.791(0.046) | 1.745(0.087) | 0.488(0.026) | 0.483(0.045) | 0.513(0.031) | 0.490(0.014) |
|  |  | SIGN | -0.181(0.032) | -0.066(0.012) | 1.509(0.021) | 1.428(0.085) | 0.098(0) | 0(0) | 0.115(0) | 0(0) |

| **Type** | **Threshold** | **Model** | **R^2^** | | **RMSE** | | **Pearson** | | **Spearman** | |
| --- | --- | --- | --- | --- | --- | --- | --- | --- | --- | --- |
|  |  |  | **IC_50_** | **KIKD** | **IC_50_** | **KIKD** | **IC_50_** | **KIKD** | **IC_50_** | **KIKD** |
| Protein-  Cluster | 0.3 | FeatNN | 0.285(0.039) | 0.326(0.050) | 1.371(0.068) | 1.647(0.067) | 0.552(0.027) | 0.586(0.036) | 0.538(0.024) | 0.577(0.039) |
|  |  | MONN | 0.247(0.058) | 0.306(0.063) | 1.383(0.046) | 1.642(0.049) | 0.537(0.042) | 0.579(0.044) | 0.515(0.048) | 0.572(0.038) |
|  |  | BACPI | 0.154(0.015) | 0.276(0.034) | 1.446(0.013) | 1.771(0.041) | 0.491(0.009) | 0.558(0.020) | 0.475(0.01) | 0.557(0.021) |
|  |  | GATNet | 0.012(0.015) | 0.161(0.009) | 1.985(0.016) | 1.786(0.010) | 0.169(0.061) | 0.423(0.006) | 0.177(0.062) | 0.366(0.006) |
|  |  | GATGCN | 0.211(0.037) | 0.252(0.015) | 1.774(0.042) | 1.687(0.017) | 0.468(0.034) | 0.519(0.009) | 0.453(0.037) | 0.486(0.009) |
|  |  | GCNNet | 0.062(0.045) | 0.224(0.021) | 1.934(0.046) | 1.718(0.023) | 0.279(0.078) | 0.487(0.017) | 0.276(0.076) | 0.444(0.031) |
|  |  | GINConvNet | 0.234(0.023) | 0.244(0.018) | 1.748(0.027) | 1.696(0.020) | 0.525(0.009) | 0.512(0.013) | 0.562(0.009) | 0.500(0.008) |
|  |  | SIGN | -0.066(0.087) | 0.047(0.125) | 1.464(0.06) | 1.433(0.098) | 0.091(0) | 0.232(0.264) | 0.112(0) | 0.266(0.301) |
|  | 0.4 | FeatNN | 0.292(0.045) | 0.324(0.029) | 1.364(0.045) | 1.643(0.039) | 0.559(0.035) | 0.586(0.028) | 0.535(0.049) | 0.572(0.018) |
|  |  | MONN | 0.244(0.048) | 0.289(0.027) | 1.399(0.060) | 1.671(0.050) | 0.551(0.035) | 0.568(0.021) | 0.528(0.04) | 0.561(0.021) |
|  |  | BACPI | 0.128(0.026) | 0.279(0.015) | 1.529(0.023) | 1.758(0.019) | 0.476(0.005) | 0.566(0.008) | 0.458(0.007) | 0.554(0.008) |
|  |  | GATNet | 0.014(0.037) | 0.176(0.014) | 1.983(0.037) | 1.770(0.015) | 0.210(0.036) | 0.433(0.017) | 0.215(0.046) | 0.382(0.021) |
|  |  | GATGCN | 0.131(0.056) | 0.240(0.018) | 1.861(0.060) | 1.701(0.020) | 0.397(0.048) | 0.507(0.013) | 0.381(0.052) | 0.476(0.018) |
|  |  | GCNNet | 0.114(0.047) | 0.214(0.045) | 1.880(0.049) | 1.729(0.049) | 0.366(0.046) | 0.478(0.028) | 0.35(0.047) | 0.436(0.031) |
|  |  | GINConvNet | 0.202(0.053) | 0.223(0.065) | 1.784(0.059) | 1.718(0.072) | 0.512(0.029) | 0.497(0.048) | 0.552(0.028) | 0.484(0.031) |
|  |  | SIGN | -0.019(0.029) | -0.015(0.007) | 1.394(0.019) | 1.446(0.005) | 0.143(0) | 0.162(0) | 0.179(0) | 0.185(0) |
|  | 0.5 | FeatNN | 0.283(0.041) | 0.307(0.021) | 1.378(0.045) | 1.659(0.057) | 0.552(0.021) | 0.570(0.020) | 0.538(0.017) | 0.562(0.017) |
|  |  | MONN | 0.249(0.065) | 0.288(0.028) | 1.426(0.027) | 1.662(0.055) | 0.556(0.050) | 0.566(0.020) | 0.536(0.062) | 0.559(0.023) |
|  |  | BACPI | 0.081(0.029) | 0.304(0.016) | 1.464(0.023) | 1.727(0.02) | 0.461(0.008) | 0.575(0.007) | 0.444(0.006) | 0.560(0.005) |
|  |  | GATNet | 0.027(0.042) | 0.174(0.031) | 1.970(0.042) | 1.772(0.033) | 0.192(0.098) | 0.432(0.029) | 0.191(0.1) | 0.375(0.051) |
|  |  | GATGCN | 0.153(0.035) | 0.245(0.022) | 1.838(0.038) | 1.694(0.025) | 0.410(0.035) | 0.508(0.020) | 0.395(0.039) | 0.48(0.027) |
|  |  | GCNNet | 0.096(0.052) | 0.233(0.033) | 1.899(0.055) | 1.708(0.037) | 0.341(0.057) | 0.493(0.035) | 0.331(0.063) | 0.453(0.049) |
|  |  | GINConvNet | 0.212(0.044) | 0.201(0.044) | 1.772(0.049) | 1.743(0.048) | 0.493(0.051) | 0.480(0.044) | 0.529(0.059) | 0.481(0.03) |
|  |  | SIGN | -0.033(0.025) | -0.058(0.059) | 1.389(0.048) | 1.515(0.041) | 0.156(0) | 0(0) | 0.186(0) | 0(0) |
|  | 0.6 | FeatNN | 0.285(0.032) | 0.343(0.050) | 1.366(0.054) | 1.640(0.045) | 0.555(0.015) | 0.598(0.032) | 0.532(0.023) | 0.583(0.027) |
|  |  | MONN | 0.204(0.059) | 0.288(0.043) | 1.440(0.045) | 1.676(0.053) | 0.514(0.030) | 0.570(0.030) | 0.500(0.03) | 0.558(0.037) |
|  |  | BACPI | 0.012(0.036) | 0.286(0.021) | 1.575(0.029) | 1.700(0.025) | 0.412(0.014) | 0.558(0.013) | 0.385(0.015) | 0.563(0.014) |
|  |  | GATNet | 0.024(0.033) | 0.184(0.023) | 1.973(0.034) | 1.761(0.025) | 0.220(0.040) | 0.444(0.020) | 0.225(0.038) | 0.393(0.034) |
|  |  | GATGCN | 0.174(0.035) | 0.225(0.040) | 1.815(0.038) | 1.716(0.045) | 0.435(0.027) | 0.499(0.018) | 0.418(0.031) | 0.467(0.016) |
|  |  | GCNNet | 0.090(0.041) | 0.232(0.013) | 1.905(0.042) | 1.710(0.014) | 0.341(0.029) | 0.491(0.012) | 0.328(0.034) | 0.454(0.018) |
|  |  | GINConvNet | 0.186(0.057) | 0.226(0.070) | 1.801(0.061) | 1.714(0.077) | 0.485(0.058) | 0.503(0.036) | 0.527(0.061) | 0.511(0.024) |
|  |  | SIGN | -0.033(0.079) | -0.015(0.013) | 1.409(0.055) | 1.436(0.009) | 0.074(0.166) | 0.084(0) | 0.088(0.198) | 0.098(0) |

**Supplementary Table 4** Comparison of the performances of FeatNN on the datasets generated from the general set and from refined set of PDBbind with the compound-clustered and protein-clustered strategy. The results of each group were obtained with 5 independent experiments by 5-fold cross-validation strategy. Details are provided in Supplementary Fig. 4.

| **FeatNN** | **Threshold** | **RMSE** | | **Pearson** | | **Spearman** | | **R^2^** | |
| --- | --- | --- | --- | --- | --- | --- | --- | --- | --- |
| **Type** |  | **refined** | **general** | **refined** | **general** | **refined** | **general** | **refined** | **general** |
| Compound-  Clustered | 0.3 | 1.38(0.071) | 1.442(0.046) | 0.735(0.029) | 0.716(0.015) | 0.729(0.032) | 0.714(0.019) | 0.512(0.055) | 0.487(0.027) |
|  | 0.4 | 1.469(0.063) | 1.54(0.079) | 0.698(0.029) | 0.669(0.04) | 0.700(0.03) | 0.677(0.041) | 0.448(0.054) | 0.406(0.073) |
|  | 0.5 | 1.448(0.09) | 1.507(0.056) | 0.699(0.061) | 0.685(0.016) | 0.690(0.064) | 0.674(0.023) | 0.440(0.113) | 0.438(0.036) |
|  | 0.6 | 1.442(0.148) | 1.429(0.088) | 0.672(0.019) | 0.66(0.028) | 0.636(0.029) | 0.634(0.022) | 0.421(0.034) | 0.398(0.043) |
| Protein-  Clustered | 0.3 | 1.642(0.104) | 1.647(0.067) | 0.558(0.125) | 0.586(0.036) | 0.552(0.127) | 0.577(0.039) | 0.278(0.18) | 0.326(0.05) |
|  | 0.4 | 1.617(0.062) | 1.643(0.039) | 0.578(0.143) | 0.586(0.028) | 0.579(0.143) | 0.572(0.018) | 0.305(0.201) | 0.324(0.029) |
|  | 0.5 | 1.615(0.111) | 1.659(0.057) | 0.579(0.066) | 0.570(0.02) | 0.577(0.081) | 0.562(0.017) | 0.305(0.074) | 0.307(0.021) |
|  | 0.6 | 1.654(0.052) | 1.64(0.045) | 0.549(0.073) | 0.598(0.032) | 0.553(0.082) | 0.583(0.027) | 0.277(0.072) | 0.343(0.05) |

**Supplementary Table 5** Comparison of the performances of the SOTA baseline (MONN) on the datasets generated from the general set and refined set of PDBbind with the compound-clustered and protein-clustered strategy. The results of each group were obtained with 5 independent experiments by 5-fold cross-validation strategy. Details are provided in Supplementary Fig. 5.

| **MONN** | **Threshold** | **RMSE** | | **Pearson** | | **Spearman** | | **R^2^** | |
| --- | --- | --- | --- | --- | --- | --- | --- | --- | --- |
| **Type** |  | **refined** | **general** | **refined** | **general** | **refined** | **general** | **refined** | **general** |
| Compound-  Clustered | 0.3 | 1.438(0.075) | 1.485(0.054) | 0.716(0.021) | 0.679(0.024) | 0.71(0.022) | 0.679(0.031) | 0.481(0.024) | 0.416(0.033) |
|  | 0.4 | 1.514(0.172) | 1.534(0.064) | 0.684(0.021) | 0.643(0.03) | 0.683(0.028) | 0.641(0.039) | 0.391(0.084) | 0.369(0.057) |
|  | 0.5 | 1.516(0.083) | 1.624(0.109) | 0.668(0.029) | 0.611(0.051) | 0.664(0.032) | 0.605(0.054) | 0.403(0.053) | 0.299(0.061) |
|  | 0.6 | 1.496(0.132) | 1.681(0.155) | 0.638(0.055) | 0.559(0.057) | 0.617(0.051) | 0.544(0.032) | 0.36(0.094) | 0.248(0.057) |
| Protein-  Clustered | 0.3 | 1.702(0.075) | 1.642(0.049) | 0.539(0.079) | 0.579(0.044) | 0.537(0.089) | 0.572(0.038) | 0.236(0.112) | 0.306(0.063) |
|  | 0.4 | 1.651(0.072) | 1.671(0.05) | 0.546(0.069) | 0.568(0.021) | 0.544(0.066) | 0.561(0.021) | 0.251(0.098) | 0.289(0.027) |
|  | 0.5 | 1.646(0.089) | 1.662(0.055) | 0.552(0.069) | 0.566(0.02) | 0.557(0.079) | 0.559(0.023) | 0.281(0.08) | 0.288(0.028) |
|  | 0.6 | 1.68(0.123) | 1.676(0.053) | 0.502(0.058) | 0.57(0.03) | 0.500(0.054) | 0.558(0.037) | 0.176(0.075) | 0.288(0.043) |

**Supplementary Table 6** Comparison of FeatNN and the SOTA baseline (MONN) with regard to the generalization ability on the datasets generated from the general and refined sets of PDBbind. The generalization abilities of FeatNN^refine^ and the SOTA baseline^refine^ decrease compared with the corresponding methods trained on the general set of PDBbind, possibly because the amount of the data affects the training process. The results of each group were tested on the dataset constructed from Binding MOAD with at least 15 independent models. Considering that the refined set only contains the measurement of K_i_ and K_d_, we use the FeatNN^general^ trained on the KIKD dataset constructed from PDBbind as the control group. Therefore, the test dataset constructed from Binding MOAD in this part is based on the measurement of K_i_ and K_d_.

| **Model** | **RMSE** | | **Pearson** | | **Spearman** | | **R^2^** | |
| --- | --- | --- | --- | --- | --- | --- | --- | --- |
|  | **general** | **refined** | **general** | **refined** | **general** | **refined** | **general** | **refined** |
| **FeatNN** | 1.656(0.033) | 1.925(0.046) | 0.647(0.017) | 0.47(0.024) | 0.656(0.02) | 0.465(0.023) | 0.359(0.025) | 0.133(0.042) |
| **SOTA Baseline** | 1.668(0.082) | 2.042(0.072) | 0.612(0.052) | 0.378(0.025) | 0.592(0.055) | 0.358(0.019) | 0.348(0.067) | 0.024(0.07) |

**Supplementary Table 7** Pretraining and Fine-tuning Results of FeatNN. This process was applied to the datasets constructed from the general set of PDBbind using IC_50_ measurement results. FeatNN^general^ is also trained on the general set of PDBbind with the measurement of IC_50_. The results of each group were obtained from 10 independent experiments by 5-fold cross-validation strategy.

| **Model** | **RMSE** | **Pearson** | **Spearman** | **R^2^** |
| --- | --- | --- | --- | --- |
| **FeatNN^general^** | 1.130(0.045) | 0.724(0.015) | 0.697(0.018) | 0.512(0.022) |
| **FeatNN^optm^** | 1.094(0.006) | 0.738(0.003) | 0.712(0.004) | 0.540(0.005) |

**Supplementary Table 8** Comparison of FeatNN^general^, FeatNN^optm^ and the SOTA baseline (MONN which is trained on the dataset constructed from the general set of PDBbind) with regard to the generalization ability. Because FeatNN^optm^ is pretrained on BindingDB only with IC_50_ measurements, all of these models are tested on the datasets constructed from Binding MOAD using the IC_50_ measurement results. Thus, both the FeatNN^general^ and SOTA baseline^general^ are trained on the general set of PDBbind using the measured IC_50_ values. The results for each group were obtained from at least 10 independent experiments.

| **Model** | **RMSE** | **Pearson** | **Spearman** | **R^2^** |
| --- | --- | --- | --- | --- |
| **SOTA Baseline^general^** | 1.339(0.044) | 0.657(0.014) | 0.621(0.016) | 0.385(0.041) |
| **FeatNN^general^** | 1.267(0.036) | 0.687(0.019) | 0.660(0.018) | 0.449(0.032) |
| **FeatNN^optm^** | 1.238(0.019) | 0.701(0.011) | 0.683(0.008) | 0.475(0.016) |

**Supplementary Table 9** Ablation study for module deletion in FeatNN. “Entire FeatNN” refers to the full proposed FeatNN model. Here, “Only Sequence” or “Only Structure” indicate only the protein sequence information or structure information being used when representing the features of protein by protein extractor. “Without Interact Mat” indicates the ablation of the compound-protein interactive matrix in the affinity learning module, which could help FeatNN to learn the interaction information between compound and protein possibly. The performances are sorted by the R^2^ values of the respective variant models. The results of each group were obtained from 10 independent experiments by 5-fold cross-validation strategy. The mean value (and SD) of each independent experimental group is shown in the table.

| **Name** | **R^2^** | **RMSE** | **Pearson** | **Spearman** |
| --- | --- | --- | --- | --- |
| Entire FeatNN | **0.512(0.022)** | **1.130(0.045)** | **0.724(0.015)** | **0.697(0.018)** |
| Without Torsion Info | 0.443(0.045) | 1.196(0.025) | 0.692(0.021) | 0.672(0.027) |
| Without MasterNode | 0.388(0.048) | 1.255(0.032) | 0.653(0.028) | 0.636(0.026) |
| Without Evo-Updating | 0.352(0.055) | 1.292(0.024) | 0.640(0.029) | 0.624(0.027) |
| Without Deep GCN | 0.326(0.055) | 1.302(0.042) | 0.611(0.024) | 0.591(0.036) |
| Only Sequence | 0.325(0.089) | 1.325(0.105) | 0.619(0.043) | 0.592(0.05) |
| Without Interact Mat | 0.317(0.078) | 1.299(0.093) | 0.584(0.071) | 0.56(0.077) |
| Only Structure | 0.157(0.058) | 1.477(0.034) | 0.480(0.022) | 0.451(0.021) |

**Supplementary Table 10** Targeting SARS-CoV-2 3C-like protease with the conformation constructed from the PDB file with PDB-id of 7CWC, we applied FeatNN and MONN (SOTA baseline) to predict the listed 28 validated bioactive compounds to test the CPAs prediction precision of FeatNN. Each result was obtained by the average of 15 independent experiments. Real affinity values were collected from published papers and are listed in the references.

| **Compound Name** | **Real Affinity Value** | **FeatNN Prediction Value** | **MONN Prediction Value** |
| --- | --- | --- | --- |
| Darunavir [8] | 4.442 | 5.467(0.282) | 5.955(0.285) |
| Cobicistat [9] | 7.495 | 6.346(0.557) | 6.457(0.583) |
| Ritonavir[10] | 4.863 | 5.430(0.326) | 5.469(0.219) |
| Tipranavir [11] | 4.875 | 5.074(0.091) | 5.733(0.264) |
| Ivermectin [12] | 5.699 | 6.279(0.405) | 7.821(0.397) |
| REMDESIVIR [11] | 4.943 | 4.388(0.274) | 4.443(0.113) |
| PF-07321332 [13] | 7.638 | 6.816(0.348) | 5.872(0.551) |
| PF-00835231 [14] | 8.398 | 6.785(0.254) | 5.947(1.406) |
| Lufotrelvir [15] | 8.097 | 5.775(0.253) | 6.049(0.572) |
| ML188 [16] | 5.824 | 5.011(0.219) | 4.709(0.236) |
| FB2001 [17] | 6.276 | 5.473(0.751) | 5.995(0.515) |
| Dalcetrapib [18] | 4.752 | 4.622(0.127) | 4.875(0.421) |
| EGCG Octaacetate [19] | 4.857 | 5.715(0.517) | 5.571(0.378) |
| Ellagic acid [19] | 4.928 | 5.810(0.511) | 5.571(0.683) |
| Curcumin [19] | 4.924 | 4.821(0.193) | 5.040(0.468) |
| Resveratrol [19] | 4.772 | 5.307(0.248) | 4.408(0.931) |
| Quercetin [19] | 4.631 | 5.750(0.291) | 5.422(0.319) |
| Chloroquine [20] | 5.567 | 4.910(0.155) | 4.805(0.210) |
| Lopinavir [11] | 5.040 | 4.954(0.172) | 4.927(0.283) |
| Azithromycin [11] | 5.674 | 5.614(0.519) | 5.753(0.510) |
| N4-Hydroxycytidine [11] | 6.523 | 5.921(0.658) | 6.023(0.707) |
| Molnupiravir [17] | 6.523 | 5.447(0.825) | 4.622(0.874) |
| GC-373 [15] | 6.456 | 6.681(0.277) | 6.549(0.465) |
| PF-07304814 [15] | 8.097 | 5.775(0.253) | 6.049(0.572) |
| Nirmatrelvir [21] | 7.796 | 6.816(0.348) | 5.872(0.551) |
| Boceprevir [11, 15] | 5.384 | 5.307(0.268) | 5.614(0.775) |
| Calpeptin [15] | 4.971 | 4.548(0.309) | 4.885(0.283) |
| Telaprevir [15] | 4.940 | 5.933(0.417) | 5.544(0.682) |

**Supplementary Table 11** Targeting Akt-1 protease with the conformation constructed from the PDB file with PDB-id of 3O96, we applied FeatNN and MONN (SOTA baseline) to predict the listed 10 validated bioactive compounds to test the CPA prediction precision of FeatNN. Each result was obtained by the average of 15 independent experiments. Real affinity values were collected from published papers and are listed in the references.

| **Compound Name** | **Real Affinity Value** | **FeatNN Prediction Value** | **MONN Prediction Value** |
| --- | --- | --- | --- |
| Capivasertib [22] | 9.046 | 6.963(0.108) | 6.763(0.809) |
| Ipatasertib [23] | 8.456 | 6.308(0.031) | 6.213(0.703) |
| GSK690693 [24] | 8.699 | 6.623(0.260) | 7.065(0.356) |
| Miransertib [25] | 8.569 | 6.815(0.874) | 6.29(0.262) |
| BAY1125976 [26] | 8.284 | 6.521(0.632) | 6.813(0.572) |
| AT7867 [27] | 7.495 | 6.311(0.788) | 6.322(0.073) |
| AT13148 [28] | 7.420 | 6.234(0.963) | 5.266(0.087) |
| Akti-1/2 [29] | 7.237 | 5.569(0.213) | 5.949(0.122) |
| Uprosertib [30] | 6.745 | 6.840(0.680) | 6.634(0.671) |
| Oridonin [31] | 5.076 | 5.302(0.433) | 5.743(0.403) |

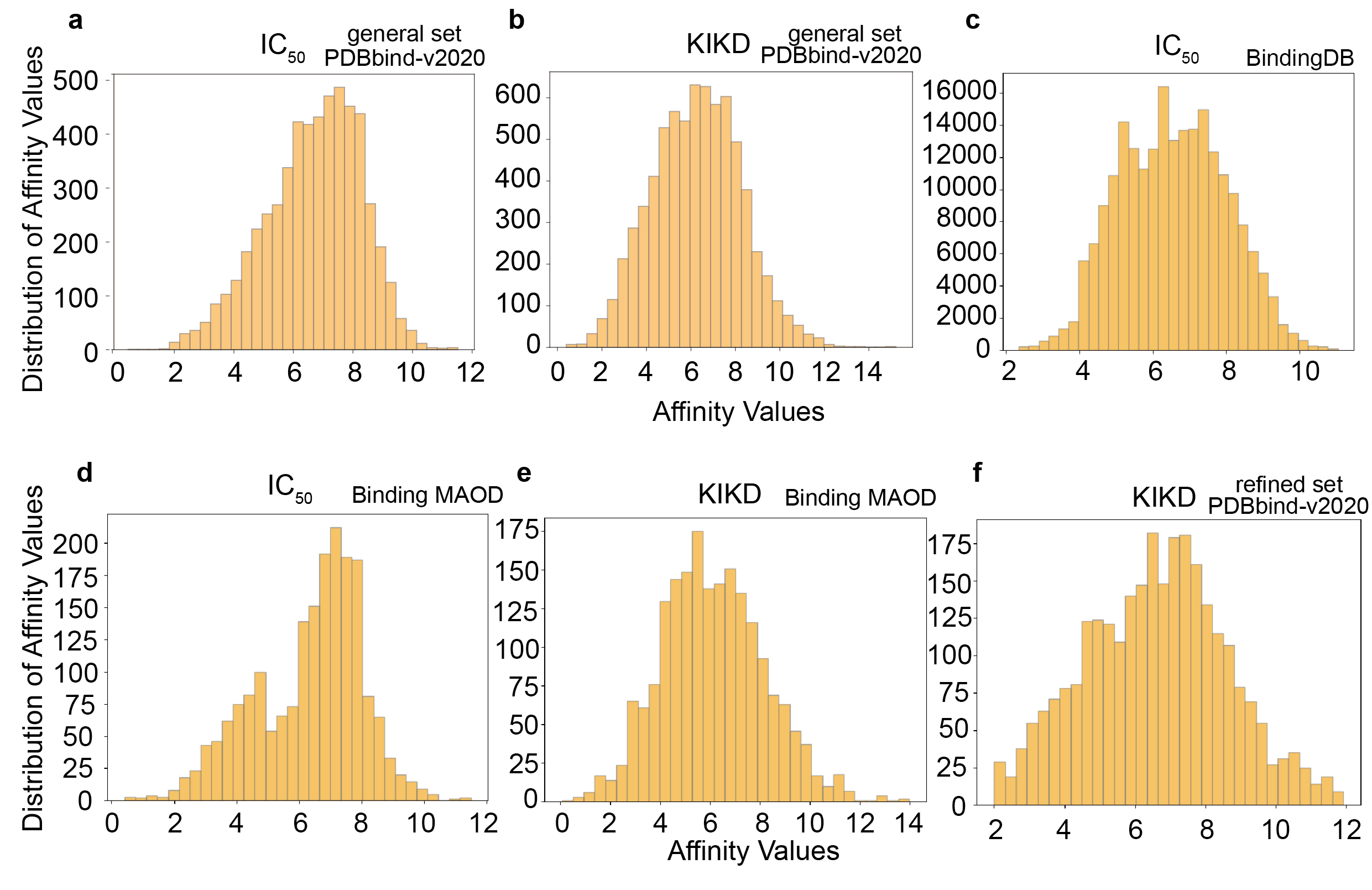

**Supplementary Fig. 2** The overall distributions of the affinity values in the PDBbind-v2020 dataset (**a.** IC_50_ and **b.** KIKD, general set) and **c.** BindingDB dataset (IC_50_). For a fair comparison of the generalization ability, we limit the datasets constructed from Binding MOAD with the measurements of **d.** IC_50_ and **e.** KIKD to the same amount of data. Thus, we constructed the dataset with IC_50_ and KIKD measurements from the “all of Binding MOAD” and “nonredundant MOAD” sets. **f.** shows the affinity value distribution on the refined set of PDBbind-v2020 that only contains the measurement of KIKD. All of these datasets produce approximately normal distributions with their values.

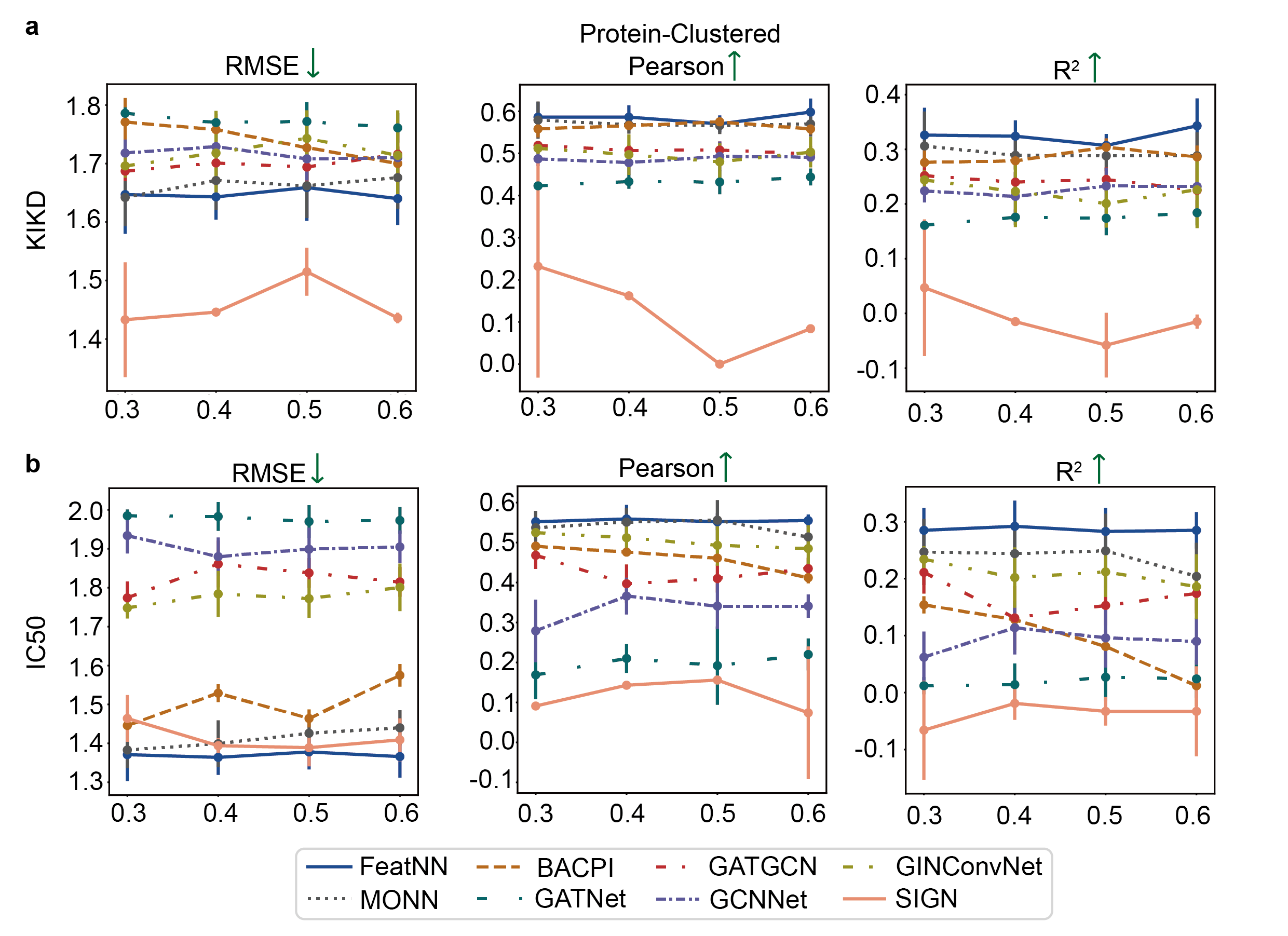

**Supplementary Fig. 3** Evaluation performances on the datasets generated from PDBbind with the protein-cluster strategy. **a.** Performance evaluated on the dataset generated from PDBbind with KIKD measurements. **b.** Performance evaluated on the dataset generated from PDBbind with IC_50_ measurements. Performance results are plotted as the mean values and standard deviations (SD) by 5-fold cross-validation with 10 independent experiments. Each point represents the mean of an independent experimental group, with error bars indicating SD. Note: the results present here were slightly different from the results reported by the original literature [32], possibly because we use PDBbind-v2020 as our benchmark database instead of PDBbind-v2016 used in their study. In addition, considering the biology means behind the data, we split the dataset into two parts ("IC50" and "KIKD" [33]) instead of simply mixing the affinity measured with "IC_50_", "K_i_", and "K_d_" together in their study. Moreover, we applied compound-cluster and protein-cluster strategies in our study to avoid data leakage caused by the biology-correlated knowledge (similarity structure or sequence in protein or compound). Thus, the results here may differ from the results in their article [32].

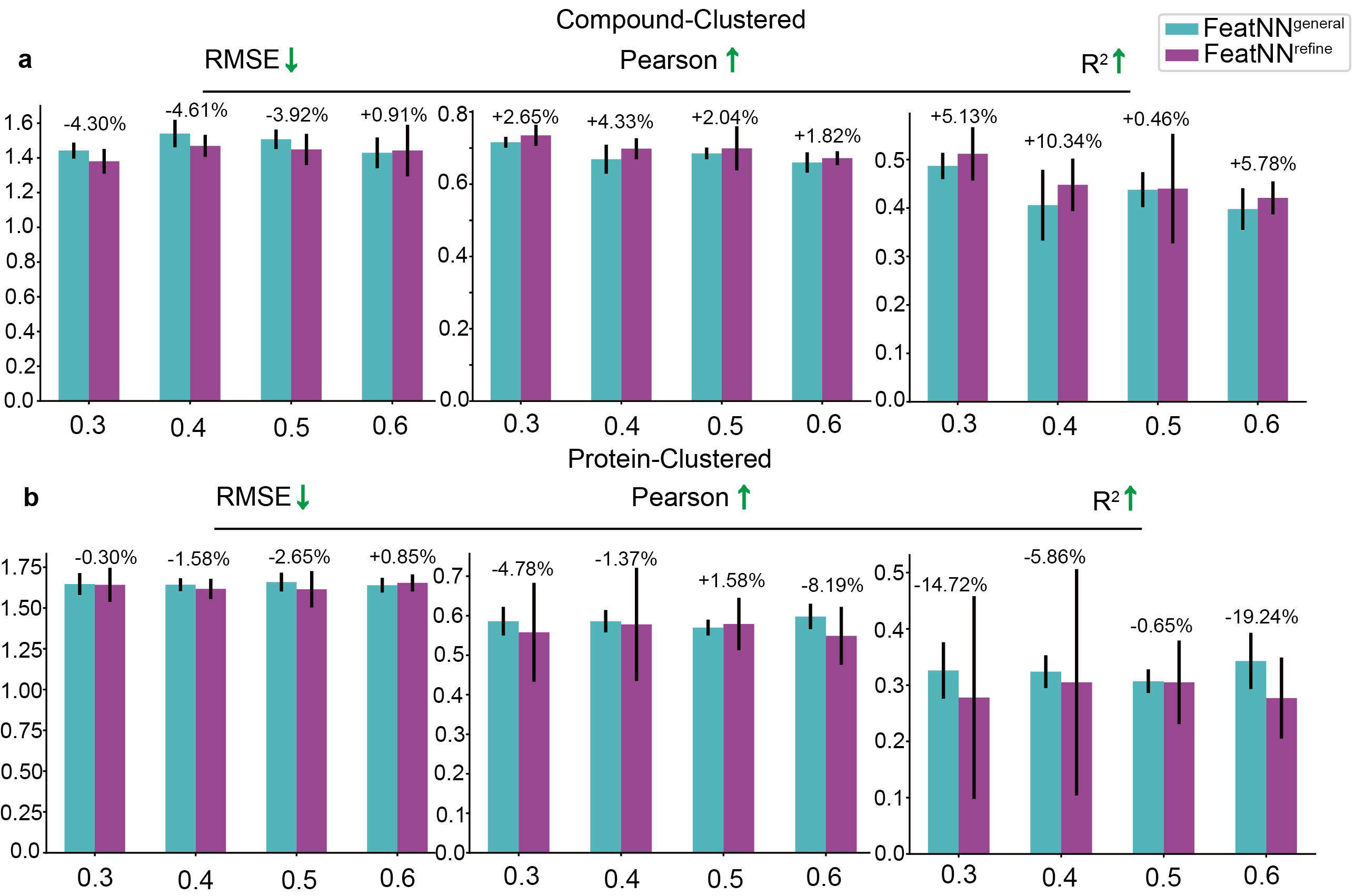

**Supplementary Fig. 4** Comparison of the performances of FeatNN on the datasets generated from the general set and refined set of PDBbind with the compound-clustered and protein-clustered strategy. **a.** Based on the compound-clustered method, FeatNN^refine^ shows improved performance compared with FeatNN^general^, possibly because high-quality structural information is introduced into the training process. **b.** However, the performance of FeatNN^refine^ based on the protein-clustered method is much worse than that of FeatNN^general^. The performance of FeatNN^refine^ declined strongly, particularly at the threshold of 0.6 (which means that less similar proteins will appear during the training process). This result may be obtained because in the training process, both the amount of data and the diversity of protein information are more important than data quality [7]. The detailed data can be found in Supplementary Table 4. Note: the text on each group bar indicates the difference between the performance of the model trained on the refined dataset and the performance of the same model trained on the general dataset. FeatNN^refine^ indicates the FeatNN trained and tested on the datasets generated from the refined set of PDBbind. FeatNN^general^ indicates the FeatNN trained and tested on the datasets generated from the general set of PDBbind. Performance results are plotted as the mean values and standard deviations (SD) obtained by 5-fold cross-validation with 5 independent experiments. Each bar represents the mean of an experimental group, with error bars indicating the SD.

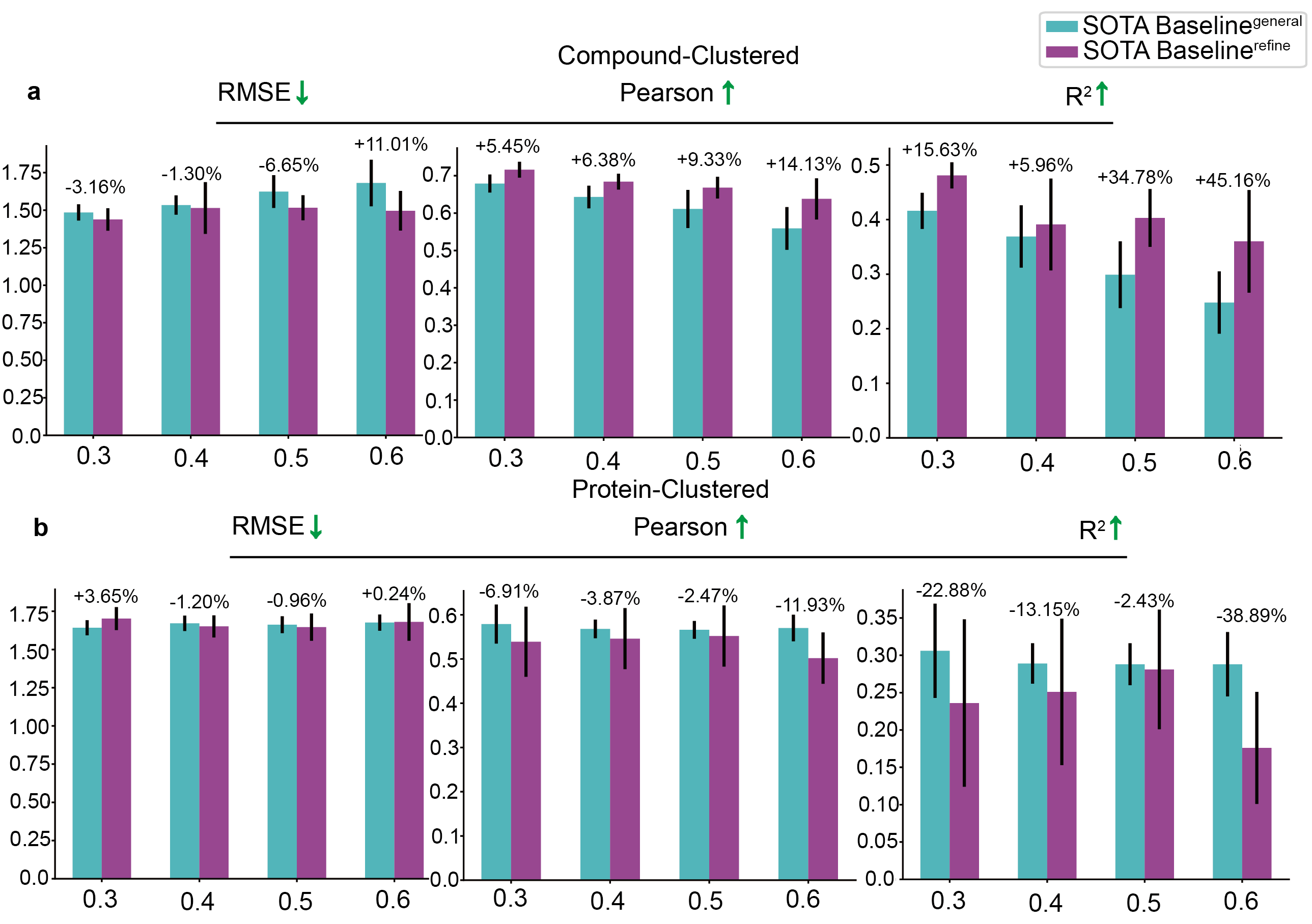

**Supplementary Fig. 5** Comparison of the performances of the SOTA baseline (MONN) on datasets generated from the general set and refined set of PDBbind with the compound-clustered and protein-clustered strategy. **a.** he performances of the SOTA baseline^refine^ based on the compound-clustered method is more or less improved with the SOTA baseline^general^, which is similar to the FeatNN^refine^ results in Supplementary Fig. 4a. **b.** The performances of the SOTA baseline^refine^ based on the protein-clustered method are much worse compared with the SOTA baseline^general^. Additionally, the performance of the SOTA baseline^refine^’ declined significantly at the threshold of 0.6, supporting the hypothesis and result shown in Supplementary Fig. 4b. The detailed data can be found in Supplementary Table 5. Note: The text on each group bar indicates the difference between the performance of the model trained on the refined dataset and the performance of the same model trained on the general dataset. SOTA Baseline^refine^ indicates the SOTA baseline trained and tested on the datasets generated from the refined set of PDBbind. SOTA Baseline^general^ indicates the SOTA baseline trained and tested on the datasets generated from the general set of PDBbind. Performance results are plotted as the mean values and standard deviations (SD) obtained by 5-fold cross-validation with 5 independent experiments. Each bar represents the mean of an experimental group, with error bars indicating the SD.

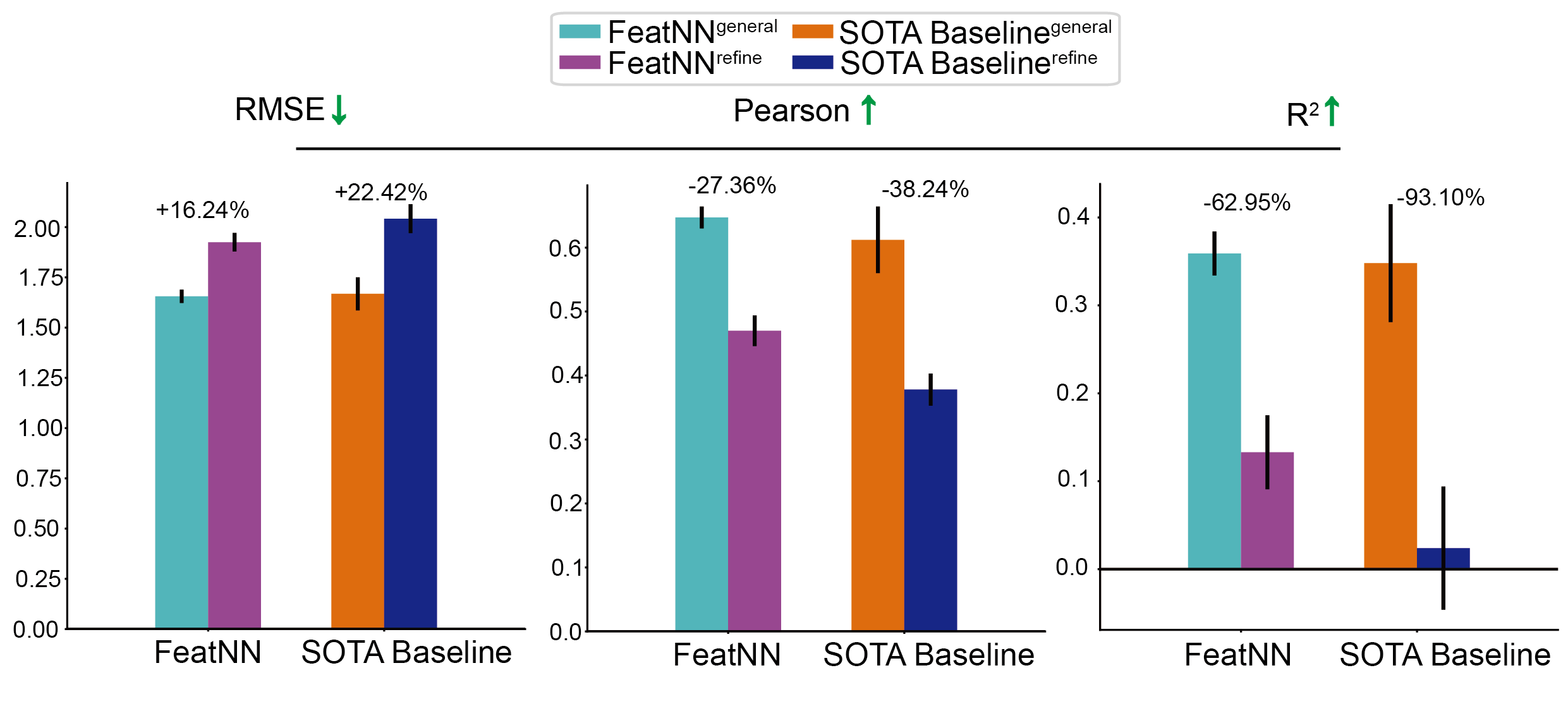

**Supplementary Fig. 6** Comparison of the generalization performance of FeatNN^general^ versus FeatNN^refine^ and SOTA baseline^general^ versus SOTA baseline^refine^ on the dataset generated from the Binding MOAD database. The detailed data can be found in Supplementary Table 6. Note: the text on each group bar indicates the difference between the performance of the model trained on the refined dataset and the performance of the same model trained on the general dataset. FeatNN^refine^ and SOTA Baseline^refine^ indicate that these two models were trained on the datasets generated from the refined set of PDBbind. FeatNN^general^ and SOTA Baseline^general^ indicate that these two models were trained on the datasets generated from the general set of PDBbind. Performance results are plotted as the mean values and standard deviations (SD). Each bar represents the mean of an experimental group, with error bars indicating the SD.

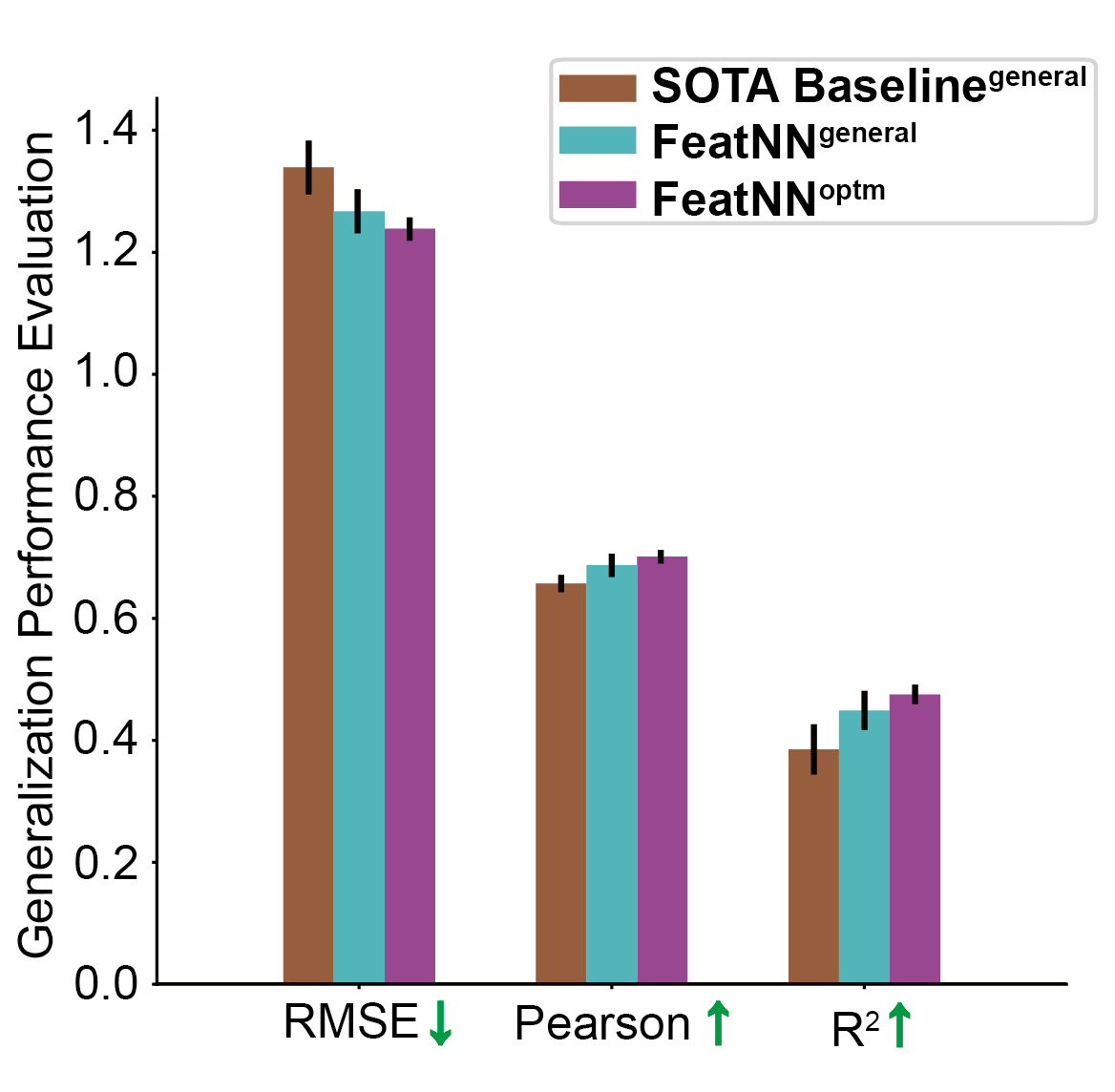

**Supplementary Fig. 7** Comparison of the generalization performances of FeatNN^general^, FeatNN^optm^ and SOTA baseline^general^ (MONN) on the dataset generated from the Binding MOAD database. Generalization of the Pearson and R^2^ of FeatNN^general^ are improved by 4.57% and 16.62% compared with the SOTA baseline^general^. FeatNN^optm^ is further improved by 2.04% and 5.79% in Pearson and R^2^ compared with FeatNN^general^. Detailed data can be found in Supplementary Table 8.

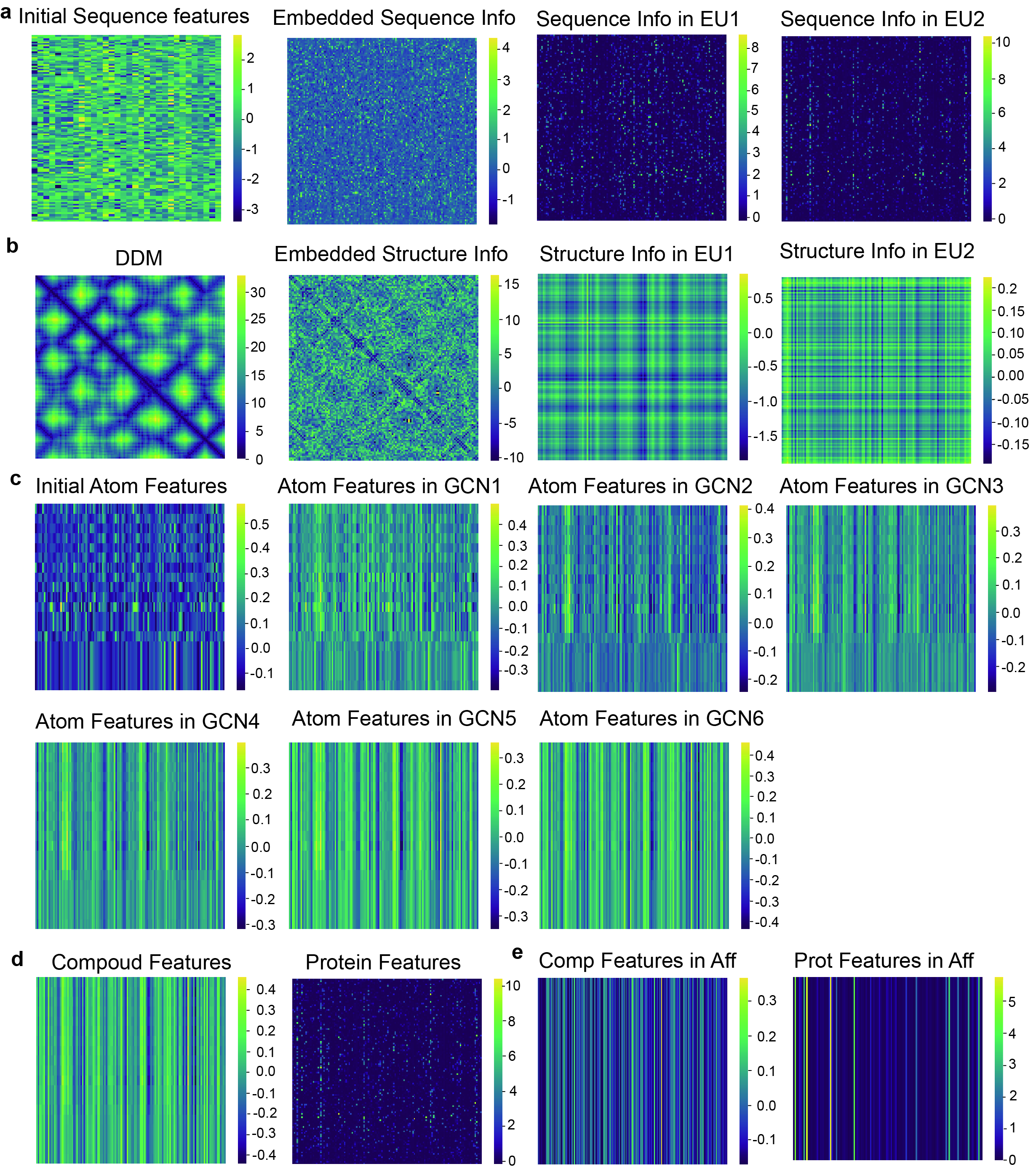

**Supplementary Fig. 8 Visualization of original features in FeatNN. a.** Sequence information in the Prot-Aggregation module and Evo-Updating module. **b.** Structure information in the Prot-Aggregation module and Evo-Updating module. **c.** Atom features in the deep GCN. **d.** Protein and compound features extracted by the protein extractor and compound extractor. **e.** Compound and protein feature interactions in the affinity learning module. Abbrev. Info: information. DDM: Discrete Distance Matrix. EU1: Evo-Updating of Layer 1. EU2: Evo-Updating of Layer 2. GCN1: GCN block of Layer 1. GCN2: GCN block of Layer 2. GCN3: GCN block of Layer 3. GCN4: GCN block of Layer 4. GCN5: GCN block of Layer 5. GCN6: GCN block of Layer 6. Aff: affinity learning module.

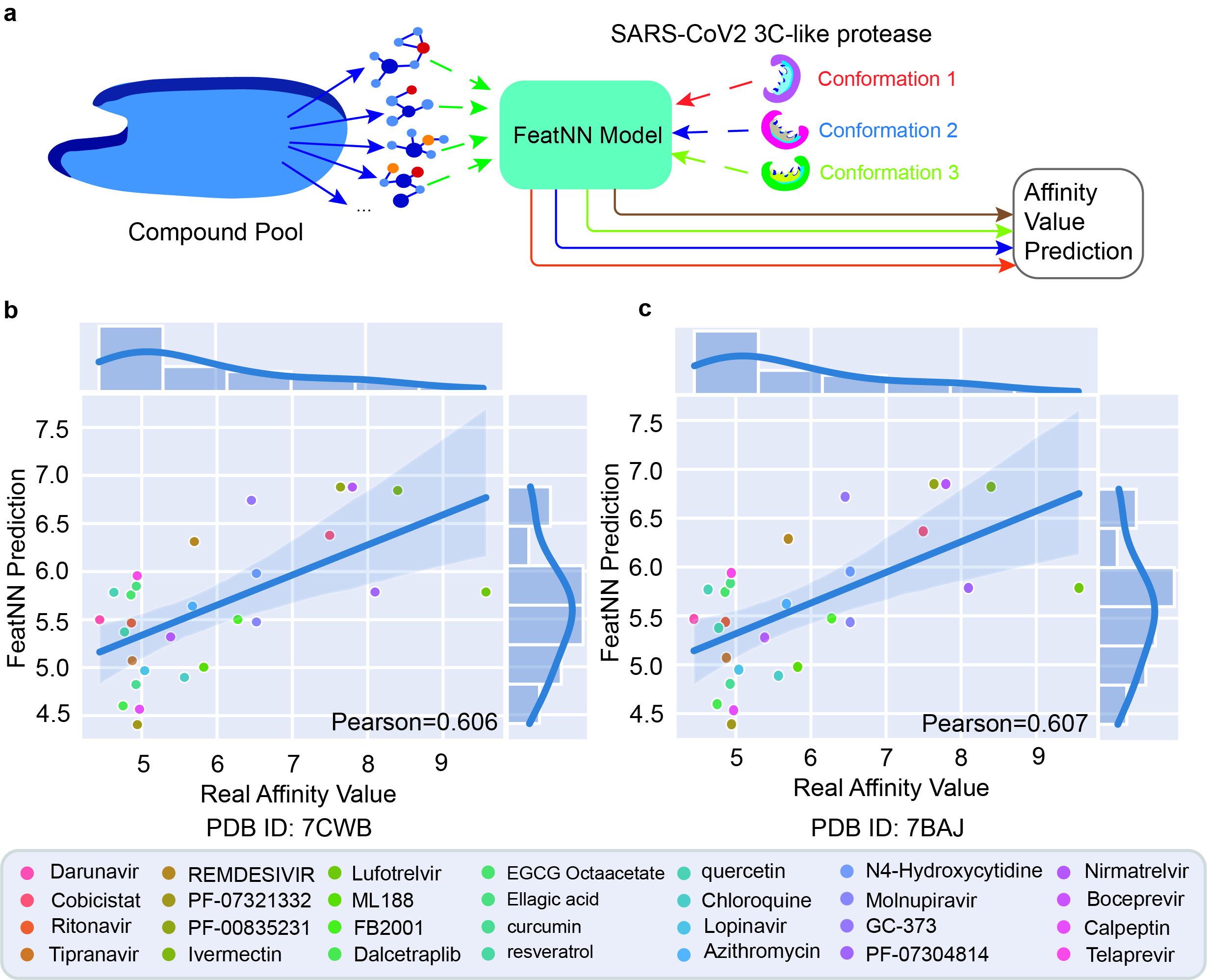

**Supplementary Fig. 9** **Affinity prediction results obtained based on receptors (SARS-CoV-2 3C-like protease) with different protein conformations**. **a.** We applied FeatNN to predict the binding affinity for 28 validation compounds and different conformations of the same target protein (SARS-CoV-2 3C-like protease, PDB-ids: 7CWC (Fig. 6b), 7CWB, 7BAJ). **b.** Affinity prediction results of 28 validation bioactive compounds (Supplementary Table 10) by FeatNN based on the conformation of 7CWB in the PDB file. **c.** Affinity prediction results of 28 validation bioactive compounds by FeatNN based on the conformation of 7BAJ in the PDB file. Each point was obtained by the average of 15 independent experiments. Note: All protein conformations were selected based on the ligand-free structure. In addition, the affinity prediction results among different protein conformations did not show significant differences.

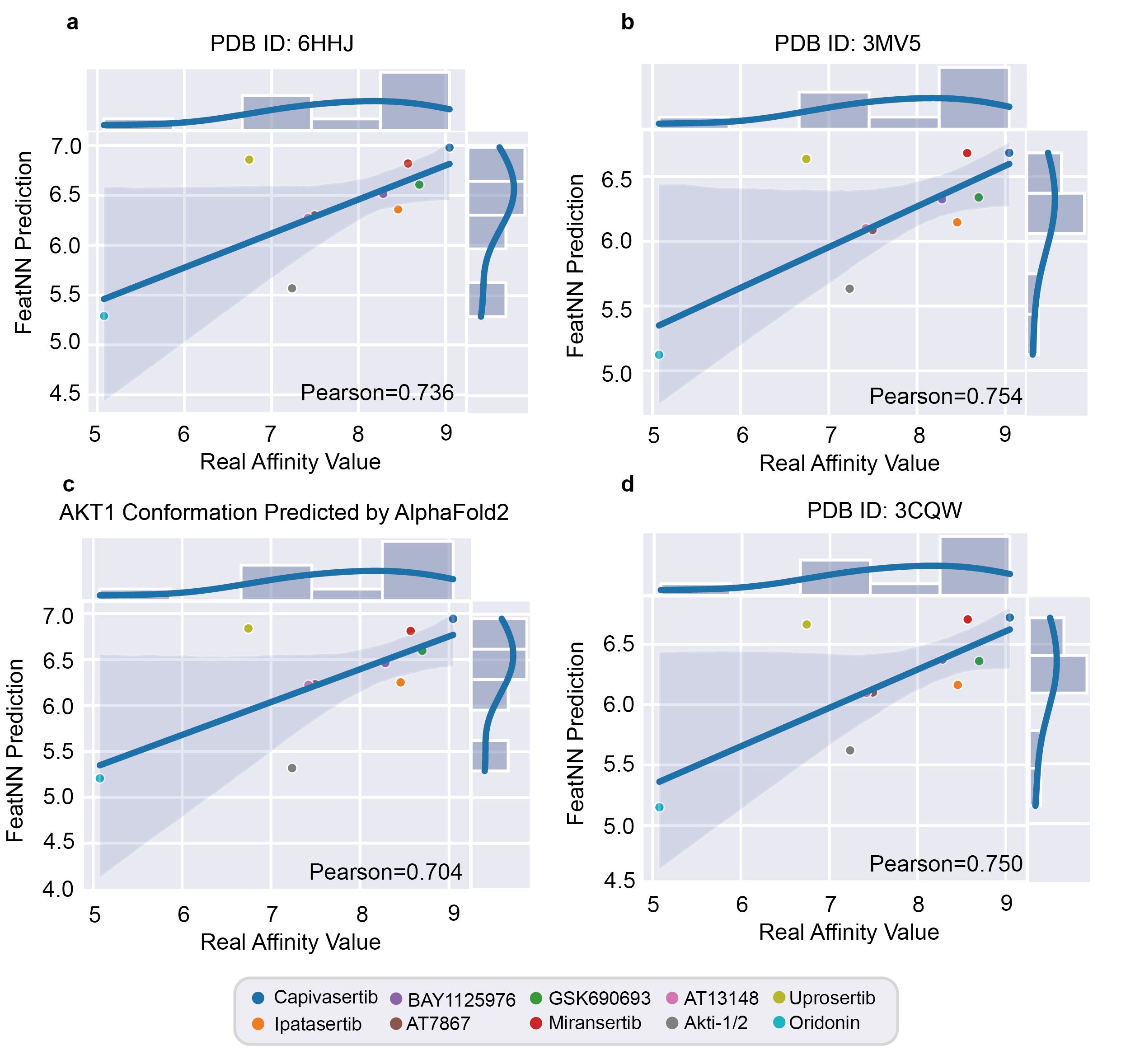

**Supplementary Fig. 10** **Affinity prediction results obtained based on receptor Akt-1 with different protein conformations.** We applied FeatNN to predict the binding affinity for 10 validation compounds (Supplementary Table 11) and different conformations of the same target protein (Akt-1, PDB-ids: 3O96 (Fig. 6d), **a.** 6HHJ, **b.** 3MV5, **d.** 3CQW and **c.** The conformation predicted by AlphaFold2 [36]). Note: We did not find the ligand-free structure in the Protein Data Bank. All protein conformations that we selected to bind with small molecules. We obtained the ligand-free structure from the prediction of AlphaFold2. In addition, the affinity prediction results among different protein conformations did not show significant differences.

#### 1.2. Influence of the number of layers with a deep graph convolution block and Evo-Updating block and the convergence rates of different models

The RMSE, R^2^, and Pearson correlation metrics are utilized to evaluate the performance of FeatNN in predicting binding affinities on the IC50 dataset generated from PDBbind. For both panels, FeatNN is evaluated under 5-fold cross-validation settings with a clustering threshold of 0.3, and the layers of the Evo-Updating block are fixed as 2. The means and SDs of the metrics over five cross-validations are shown in Supplementary Figs. 11a-c. It is clear that FeatNN performance is gradually optimized as the number of GCN layers increases (from 1 to 6 layers). The FeatNN with a deep GCN block outperforms the same model without the deep GCN block, emphasizing the importance of addressing the oversmoothing problem in the traditional GCN.

For Supplementary Fig. 11d, FeatNN is evaluated under 5-fold cross-validation settings with a clustering threshold of 0.3, and the layers of the deep GCN block are fixed as 6. The performances of the FeatNN with different numbers of layers of the Evo-Updating block are shown in Supplementary Fig. 11d.

For Supplementary Fig. 11e, we test FeatNN, MONN, BACPI, and GraphDTA on the IC_50_ dataset and evaluate them under 5-fold cross-validation settings with a clustering threshold of 0.3. We use the RMSE in each epoch to represent the convergence rate. The convergence rates of different modes are given below (Supplementary Fig. 11e).

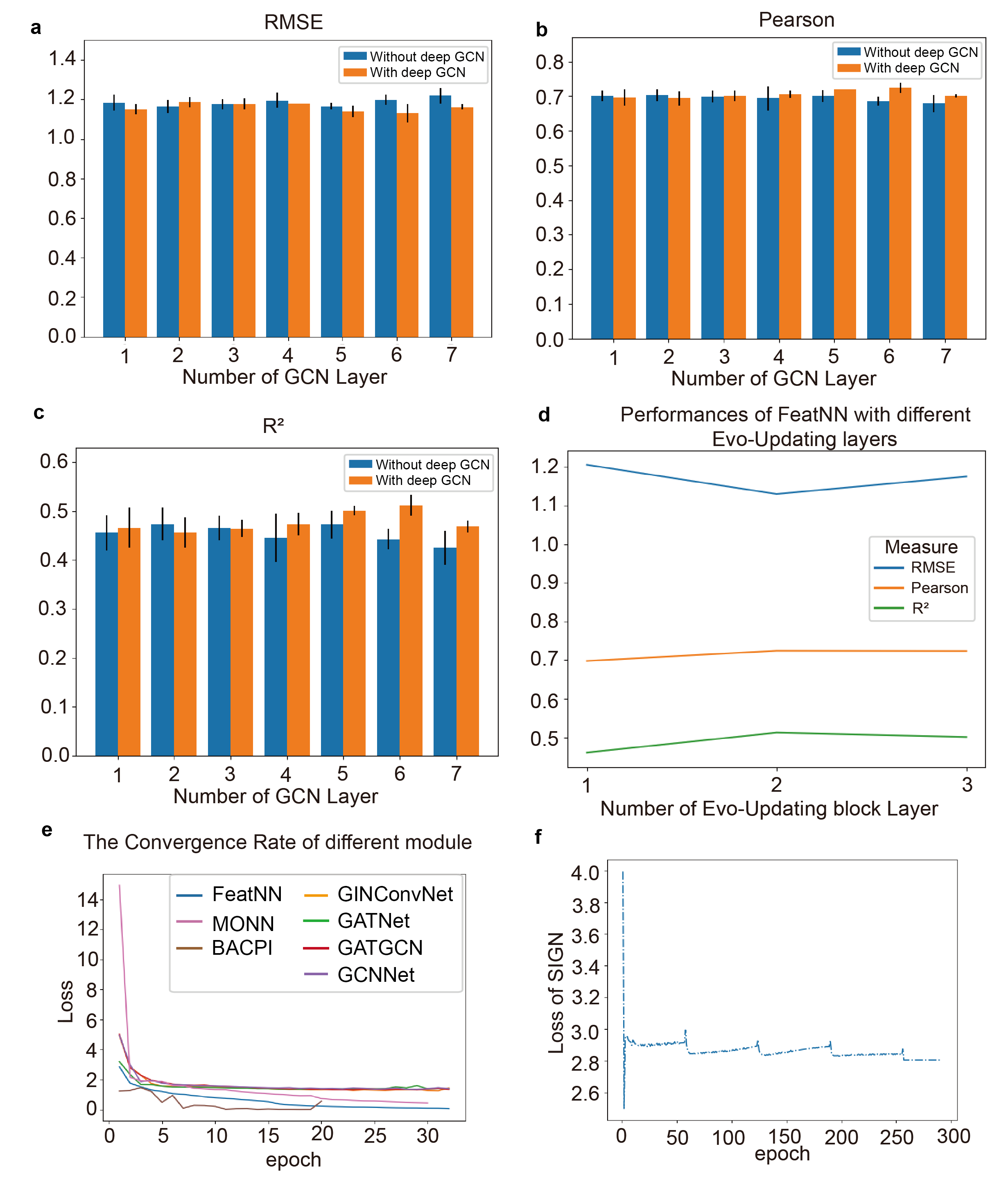

**Supplementary Fig. 11 Influence of the number of layers with a deep graph convolution block and Evo-Updating block and the convergence rates of different models. a-c.** With the deepening of the GCN module layers (from 1 layer to 6 layers), the RMSE, Pearson and R^2^ performance metrics of CPA prediction improve. The performance metrics of FeatNN with the deep GCN block are superior to those of FeatNN without a deep GCN block. **d.** Performance metrics of FeatNNs with different numbers of Evo-Updating block layers. **e.** During the training process with 32 epochs, the convergence rate of FeatNN is compared with those of the baseline models. **f.** Since 300 epochs were used for SIGN , and the value of the initial loss exceeds 1e6, we remove the outliers and separately present the result of SIGN here.

### 2. Supplementary Architecture

#### 2.1. Notation Definitions

*Linear* ($\cdot$) indicates a fully connected linear layer without an activation function. $matmul\left( \cdot\right)$ represents the multiplication operation between two tensors. $DimentionReshape(\cdot)$ indicates the dimension reshaping operation. $Embedding(\cdot)$ indicates the embedding layer based on the word embedding strategy. $concat$($\cdot$) indicates the concatenation operation between two tensors. $LayerNorm(\cdot)$ indicates the layer normalization operation on a specific channel with learnable per-channel gains and biases. $COMBINE(\cdot)$ indicates the aggregation operation based on the message passing mechanism. $dropout(\cdot)$ is the dropout regulation method. $tanh(\cdot)$, $sigmoid(\cdot)$, $Softmax\left( \cdot\right)$ and $gelu(\cdot)$ serve as the activation functions. For the definition of the calculation process, we use $\odot$ for the elementwise product and $\oplus$ for the outer sum.

#### 2.2. Block I: Compound Extractor

The deep GCN and multihead attention representation are illustrated in the compound extractor module (Supplementary Fig. 13). In the graph network, the compound information is extracted using graph representation, in which the main nodes (each atom in the compound) and the master node (the node sum of all atoms in the compound) are employed to aggregate the local and global information of the compound, respectively. A deep graph convolution unit (Supplementary Fig. 12) and a multihead attention representation strategy are used to update the main node information. The gate warp strategy interactively regulates and updates the information between the main nodes and the master node. A gated recurrent unit (GRU) is responsible for aggregating the multilayer information in the compound extractor module and updating the features of the main nodes and the master node.

In the GCN, the message passing unit gathers the information of a node's neighbors and passes it to that node for local feature updating (Fig. 1a, Supplementary Fig. 12). Here, we apply a master node to maintain the global features for nodes over long distances. This helps to mitigate the oversmoothing problem in the GCN (for a detailed explanation of the oversmoothing problem, please refer to Supplementary Note 1.1). Moreover, by applying the master node, the number of layers in the GCN can be deepened to better extract the features of compounds, thus contributing to the multihead attention representation (Supplementary Fig. 14). Finally, the local information and global information representations of compounds are jointly input into the affinity prediction module with the protein features learned from the protein extractor module, ultimately benefiting the CPA prediction process.

##### 2.2.1. Algorithm 1: Deep GCN Block

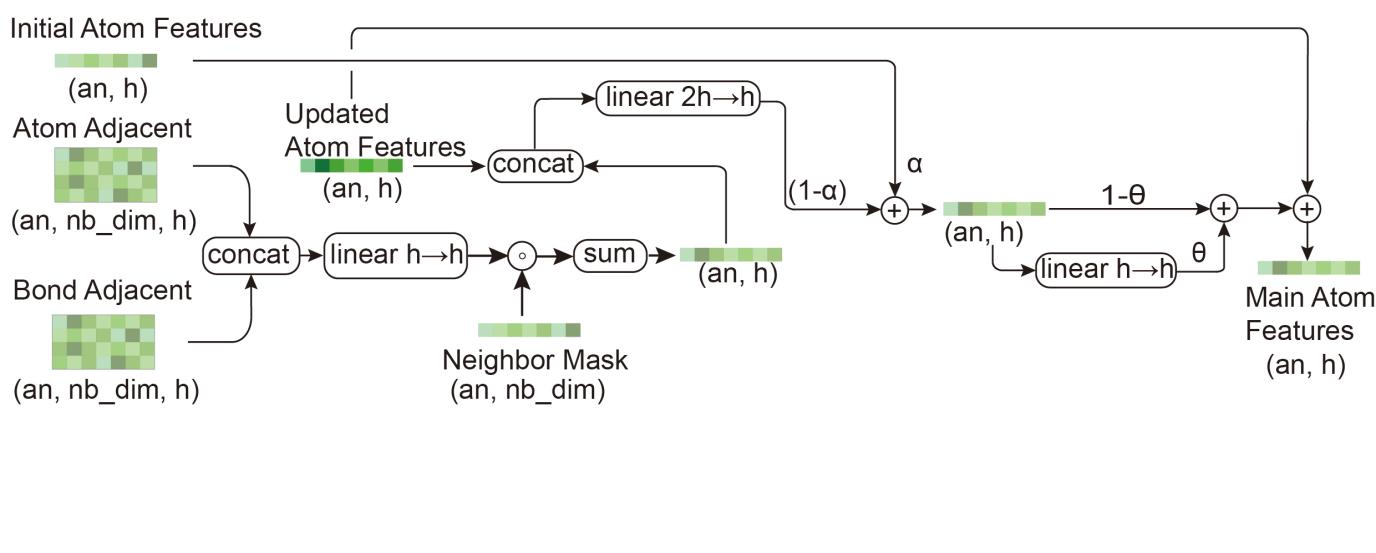

**Supplementary Fig. 12 Deep GCN block.** Atom features are combined by a message passing mechanism in a deep GCN block.

**Define:** $F_{N_{atom},h}^{vertex}$ indicates the features of the atoms in the compound, and $F_{N_{atom},nbs}^{edge}$ indicates the features of the bonds in the compound. ${Adj}_{N_{atom},nbs}^{atom}$ and ${Adj}_{N_{atom},nbs}^{bond}$ are the adjacency matrices of atoms and bonds, respectively, which are used to aggregate adjacent vertex and bond information into each atom. $F_{N_{atom},h}^{h0}$ denotes the initial features of atoms that are vital for addressing the oversmoothing problem. $theta$ and $alpha$ are hyperparameters of the residual connection in the deep GCN block.

**def** $GraphCovNN$({$F_{N_{atom},h}^{vertex}$},{$F_{N_{atom},nbs}^{edge}$},{${Adj}_{N_{atom},nbs}^{atom}$},{${Adj}_{N_{atom},nbs}^{bond}$},{$F_{N_{atom},h}^{h0}$},{$theta$},{$alpha$}):

$${ver}_{neighbor} = COMBINE (F_{N_{atom},h}^{vertex},{Adj}_{N_{atom},nbs}^{atom})$$

$${edge}_{neighbor} = COMBINE(F_{N_{atom},nbs}^{edge},{Adj}_{N_{atom},nbs}^{bond})$$

$${con}_{neighbor} = {concat}_{neighbor}({ver}_{neighbor}, {edge}_{neighbor})$$

$$neighbor\_label = gelu(Linear({con}_{neighbor}))$$

$$hi = {Linear(concat}_{h}(F_{N_{atom},h}^{vertex}, neighbor\_label))$$

$$support = (1-alpha)\odot hi+alpha\odot F_{N_{atom},h}^{h0}$$

$$output = theta\odot Linear(support)+(1-theta)\odot support$$

**return**$\boldsymbol{\to}${$F_{N_{atom},h}^{output}\}$

##### 2.2.2. Algorithm 2: Compound Extractor

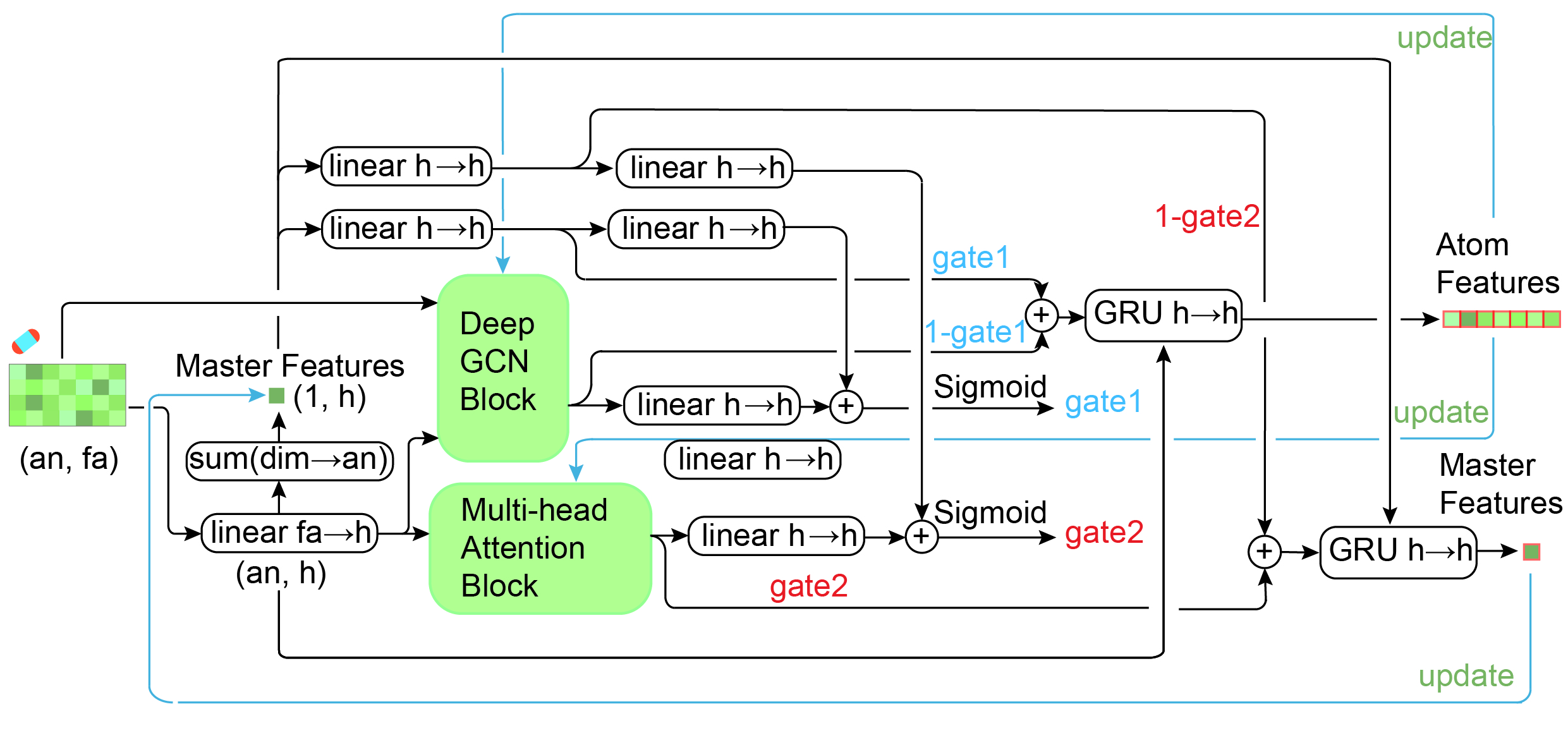

**Supplementary Fig. 13 Outlines of the compound extractor.** The deep GCN block and multihead attention block function form the core of the compound extractor.

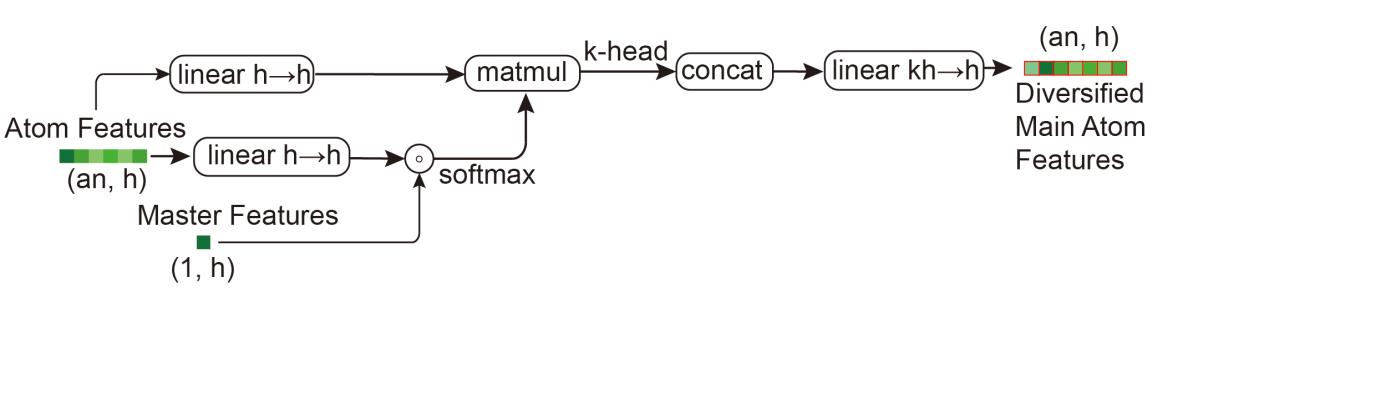

**Supplementary Fig. 14 Multihead attention block in the compound extractor.** A multihead attention block is applied to enhance the diversification of atom features.

**Define:** The definitions of $F_{N_{atom},h}^{vertex}$, $F_{N_{atom},nbs}^{edge},F_{N_{atom},nbs}^{edge}$, ${Adj}_{N_{atom},nbs}^{atom}$ and ${Adj}_{N_{atom},nbs}^{bond}$ are the same as those in Algorithm 1. In particular, ${F_{1,h}^{master}}_{l_{c}}$ and ${F_{N_{atom},h}^{atom}}_{l_{c}}$ indicate the master features (the sum over all atom features in the compound) and the atom features extracted from the GCN’s $l_{c}th$ layer. ${F_{1,h}^{master}}_{l_{0}}$ and ${F_{N_{atom},h}^{atom}}_{l_{0}}$ indicate the initial states of the master and atom features. ${mask}_{N_{atom}}^{vertex}$ indicates the mask matrix of the vertex in the compound graph.

**def** $CompExtractor$({$F_{N_{atom},h}^{vertex}$},{$F_{N_{atom},nbs}^{edge}$},{${Adj}_{N_{atom},nbs}^{atom}$},{${Adj}_{N_{atom},nbs}^{bond}$},{${mask}_{N_{atom}}^{vertex}$}):

$${F_{N_{atom},h}^{atom}}_{l_{0}} = gelu(Linear(F_{N_{atom},h}^{vertex}))$$

$$F_{N_{atom},h}^{h0} = {F_{N_{atom},h}^{atom}}_{l_{0}}$$

$${F_{1,h}^{master}}_{l_{0}} = sum({F_{N_{atom},h}^{atom}}_{l_{0}}\odot{mask}_{N_{atom}}^{vertex})$$

**for**$l_{c}\in[l_{0},\ldots. N_{Comp}]:$

**for**$k \in[0,\ldots. head\_num]:$

$$main\_vertex = tanh(Linear({F_{N_{atom},h}^{atom}}_{l_{c}-1}))$$

$$vertex=Linear(main\_vertex\odot{F_{1,h}^{master}}_{l_{c}-1})$$

$$attention\_score=softmax(vertex+{mask}_{N_{atom}}^{vertex})$$

$$k\_head\_atom\_to\_master=bmm(attention\_score,Linear({F_{N_{atom},h}^{atom}}_{l_{c}-1}))$$

**if** k $==$ 0:

$$m\_atom\_to\_master=k\_head\_atom\_to\_master$$

**else**:

$$m\_atom\_to\_master=concat(m\_atom\_to\_master, k\_head\_atom\_to\_master)$$

**end if**

$$atom\_to\_master=tanh(Linear(m\_atom\_to\_master))$$

$$atom\_feat=dropout({F_{N_{atom},h}^{atom}}_{l_{c}-1})$$

$${vert}_{agg}=GraphCovNN(atom\_feat, F_{N_{atom},nbs}^{edge},{Adj}_{N_{atom},nbs}^{atom},{Adj}_{N_{atom},nbs}^{bond}, F_{N_{atom},h}^{h0},theta, alpha)$$

$$master\_to\_atom=gelu(Linear({F_{1,h}^{master}}_{l_{c}-1}))$$

$$master\_agg=gelu(Linear({F_{1,h}^{master}}_{l_{c}-1}))$$

$${gate}_{atom}=sigmoid(Linear({vert}_{agg})+Linear(master\_to\_atom))$$

$$updated\_atom=(1-{gate}_{atom})\odot{vert}_{agg}+{gate}_{atom}\odot master\_to\_atom$$

$${F_{N_{atom},h}^{atom}}_{l_{c}}=GRU(updated\_atom,{F_{N_{atom},h}^{atom}}_{l_{c}-1})$$

$${gate}_{master}=sigmoid(Linear(master\_self)+Linear(atom\_to\_master))$$

$$updated\_master=(1-{gate}_{master})\odot master\_agg+{gate}_{master}\odot atom\_to\_maste$$

$${F_{1,h}^{master}}_{l_{c}}=GRU(updated\_master, {F_{1,h}^{master}}_{l_{c}-1})$$

**end for**

**end for**

**return**$\boldsymbol{\to}\boldsymbol{\{}{F_{N_{atom},h}^{atom}}_{N_{Comp}}\},\{{F_{1,h}^{master}}_{N_{Comp}}\}$

#### 2.3. Block II: Protein Extractor Module

Most importantly, the direct introduction of the 3D structures of proteins may drastically increase the computational costs of our model. The continuous Euclidean distance information between protein residues in the traditional distance matrix is difficult to discriminate within a small scope. In this study, the protein distance matrix is discretely encoded, and its continuous values are divided into 40 mapping intervals that conform to a normal distribution in statistics. Between 3.25 Å and 50.75 Å, the distance matrix is mapped to 38 intervals with equal distances and widths (1.25 Å per unit). Two additional intervals are added to store any larger distances (when the distances between residues are greater than 50.75 Å) and smaller distances (when the distances between residues are less than 3.25 Å). Therefore, the computational cost is greatly reduced. Furthermore, the sequence information and torsion angle information of the protein are introduced in the protein extractor module, and the DDM and protein residue sequence information are further characterized.

Many traditional networks can only update one type of data source at a time, while a multimodal mechanism can learn more comprehensive information from a variety of data sources. In contrast with previous work, we innovatively aggregate the sequence and structure features of the proteins with the protein aggregation unit (Prot-Aggregation, Supplementary Fig. 17). The torsion matrix is aggregated into sequence features through the linear mapping and Hadamard product operation in the protein aggregation unit. A mechanism employed by the evolutionary updating block (Evo-Updating) can interactively update these two properties. The Prot-Aggregation block and Evo-Updating block jointly construct the backbone of the protein extractor module (Supplementary Fig. 15). The DDM updates sequence features by summation over its columns (Supplementary Fig. 18).

Message communication from the evolving DDM to the sequence features in the Evo-Updating unit (Supplementary Fig. 18) is enabled by an enormous amount of matrix multiplications that serves as the core of the protein encoder module (Supplementary Fig. 16). The embedded distance matrix (embedded DM) is transformed into distance vectors that possess the same shapes as the sequence features through column sum and row sum operations. A merging matrix is constructed by multiplying the embedded sequence features with the distance vectors through a batch-dot-product operation, and this matrix is then added to the features of the embedded DM. The sequence features are finally renovated by the attention mechanism and gate unit updating methods. These sequence features are then evolutionarily projected to structure information through the outer sum operation and gate unit updating method. Such an intricate network architecture satisfies the requirement of multimodal pattern feature extraction, ensuring that the overall Evo-Updating unit can fully mix information regarding sequence and structure features and is sufficient for accurate affinity prediction.

##### 2.3.1. Algorithm 3: Protein Extractor

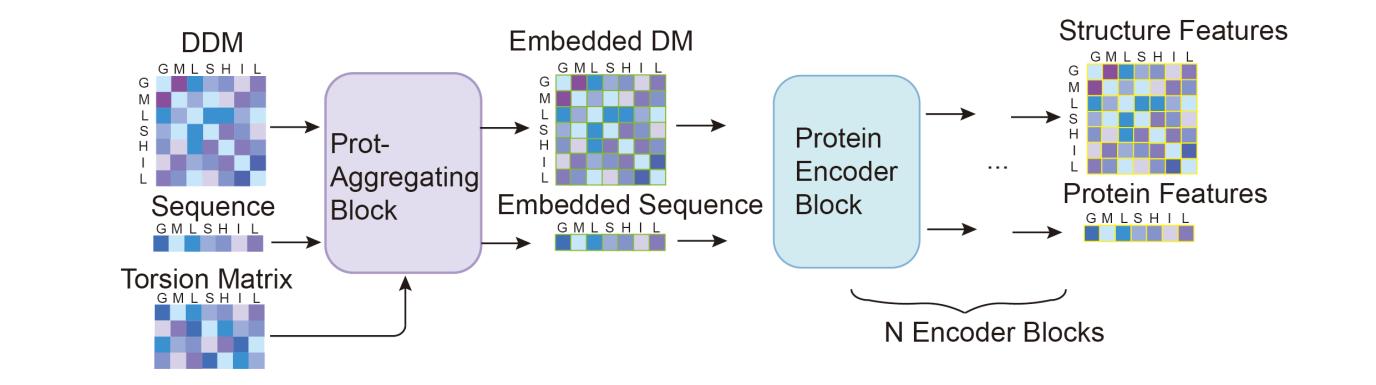

**Supplementary Fig. 15 Overview of the protein extractor.** The protein extractor consists of the Prot-Aggregation block and protein encoder block.

**Define:** ${Seq}_{init},$ ${DDM}_{init}$ and ${TorMat}_{init}$ indicate the initial information of the protein residue sequence, *DDM* and torsion matrix, respectively. ${mask}_{N_{res}}^{Seq}$ and ${mask}_{N_{res},N_{res}}^{DDM}$ are the mask matrices of the protein residue sequence, and $DDM$.

**def** *ProtExtractor*({${Seq}_{init}$},{${DDM}_{init}$},{${TorMat}_{init}$},{${mask}_{N_{res},N_{res}}^{DDM}$},{${mask}_{N_{res}}^{Seq}$}):

$${F_{N_{res}, h}^{seq}}_{init},{F_{N_{res}, e}^{DDM}}_{init}\leftarrow ProtAggregation({Seq}_{init},{DDM}_{init},{TorMat}_{init},{mask}_{N_{res},N_{res}}^{DDM},{mask}_{N_{res}}^{Seq})$$

$$F_{N_{res}, h}^{seq}, F_{N_{res}, e}^{DDM}\leftarrow ProtEncoder(F_{N_{res}, h}^{seq}, F_{N_{res}, e}^{DDM},{mask}_{N_{res}}^{Seq},{mask}_{N_{res},N_{res}}^{DDM})$$

**return** $\boldsymbol{\to}$ $\boldsymbol{\{}F_{N_{res}, h}^{seq}\},\{F_{N_{res}, e}^{DDM}\}$

##### 2.3.2. Algorithm 4: Protein Encoder

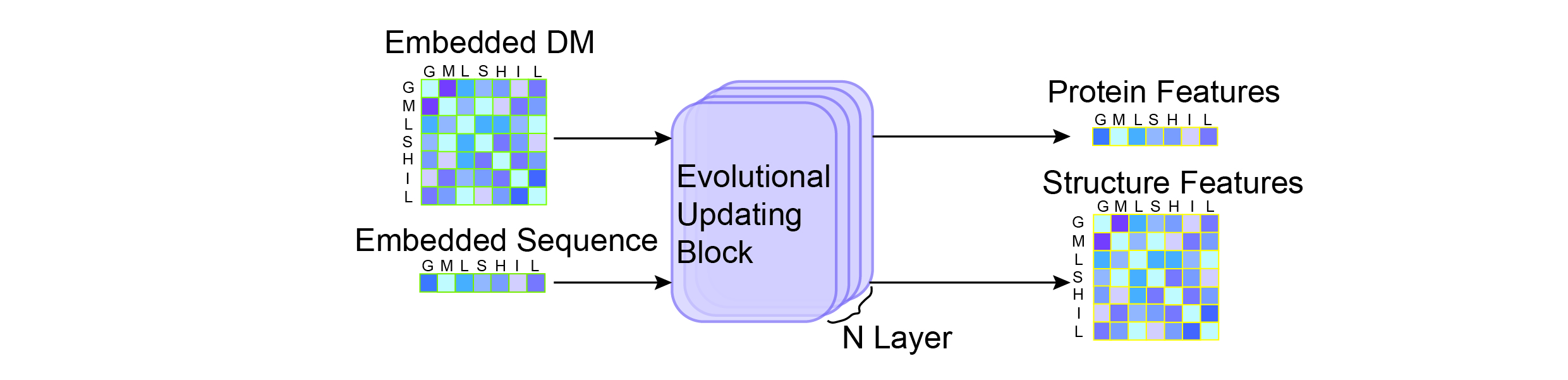

**Supplementary Fig. 16 Architecture of the protein encoder.** With the Evo-Updating block, the protein encoder interactively updates the sequence and structure features.

**Define:** $F_{N_{res}, h}^{seq}$ and $F_{N_{res}, e}^{DDM}$ are the features of the residue sequence and *DDM* embedded by the Prot-Aggregation Algorithm, and ${mask}_{N_{res}}^{Seq}$ and ${mask}_{N_{res},N_{res}}^{DDM}$ are the mask matrices of the protein residue sequence and $DDM$, respectively. ${F_{N_{res}, h}^{seq}}_{l_{Encoder}}$ and ${F_{N_{res}, e}^{DDM}}_{l_{Encoder}}$indicate the DDM and sequence features extracted from the $l_{Encoder}th$ layer of the protein encoder.

**def** *ProtEncoder*({$F_{N_{res}, h}^{seq}$},{$F_{N_{res}, e}^{DDM}\},\{{mask}_{N_{res}}^{Seq}\},\{{mask}_{N_{res},N_{res}}^{DDM}$}):

**for all** $l_{Encoder}\in[init, 1,2,...,N_{Prot}] do:$

$${{Seq}_{N_{res}, h}^{final}}_{l_{Encoder}}, {{DDM}_{N_{res}, e}^{final}}_{l_{Encoder}}\leftarrow EvoUpdating({F_{N_{res}, h}^{seq}}_{l_{Encoder}-1},{F_{N_{res}, e}^{DDM}}_{l_{Encoder}-1},{mask}_{N_{res}}^{Seq},{mask}_{N_{res},N_{res}}^{DDM})$$

**end for**

**return** $\boldsymbol{\to}$ {${{Seq}_{N_{res}, h}^{final}}_{N_{Prot}}$}, {${{DDM}_{N_{res}, e}^{final}}_{N_{Prot}}$}

##### 2.3.3. Algorithm 5: Prot-Aggregation

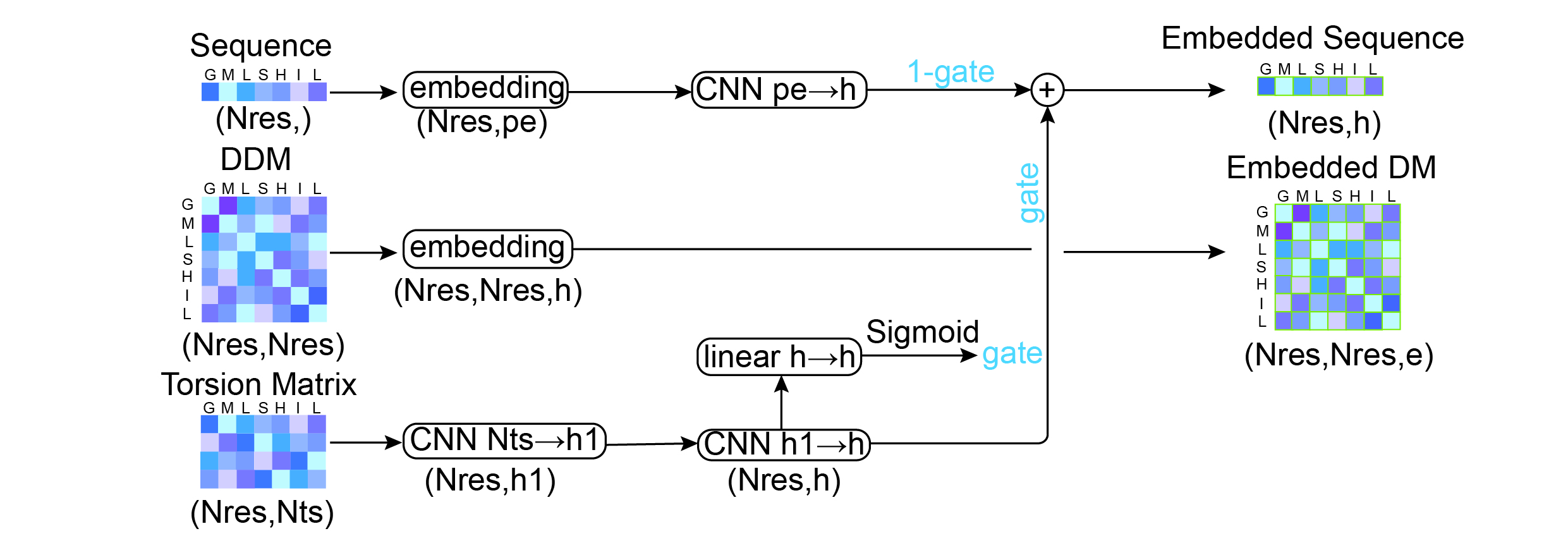

**Supplementary Fig. 17 Prot-aggregation block.** Based on the input raw protein data, the sequence features and structure features are embedded and aggregated in the Prot-Aggregation block.

**Define:** The definitions of ${Seq}_{init},$ ${DDM}_{init}$, ${TorMat}_{init}$, ${mask}_{N_{res}}^{Seq}$ and ${mask}_{N_{res},N_{res}}^{DDM}$ are the same as those in Algorithm 3.

**def** $ProtAggregation$({${Seq}_{init}$},{${DDM}_{init}$},{${TorMat}_{init}$},{${mask}_{N_{res},N_{res}}^{DDM}$},{${mask}_{N_{res}}^{Seq}$}):

$$seq\_embed = Embedding({Seq}_{init})$$

$$seq\_features = CNN(seq\_embed\odot{mask}_{N_{res}}^{Seq})$$

$$torsion\_embed = {CNN}_{Nts\to h1}({TorMat}_{init})$$

$$torsion\_vector = {CNN}_{h1\to h}(torsion\_embed)$$

$gate = Sigmoid(Linear(torsion\_vector)$)

$${Embed}_{N_{res} ,h}^{Seq} \leftarrow gate\odot torsion\_vector+(1-gate)\odot seq\_features$$

$${Embed}_{N_{res} ,N_{res} ,e}^{DM} = Embedding({DDM}_{init})\odot{mask}_{N_{res},N_{res}}^{DDM}$$

**return** $\boldsymbol{\to}$ $\left\{ {Embed}_{N_{res} ,N_{res} ,e}^{DM} \right\},\{{Embed}_{N_{res} ,h}^{Seq}\}$

##### 2.3.4. Algorithm 6: Evo-Updating

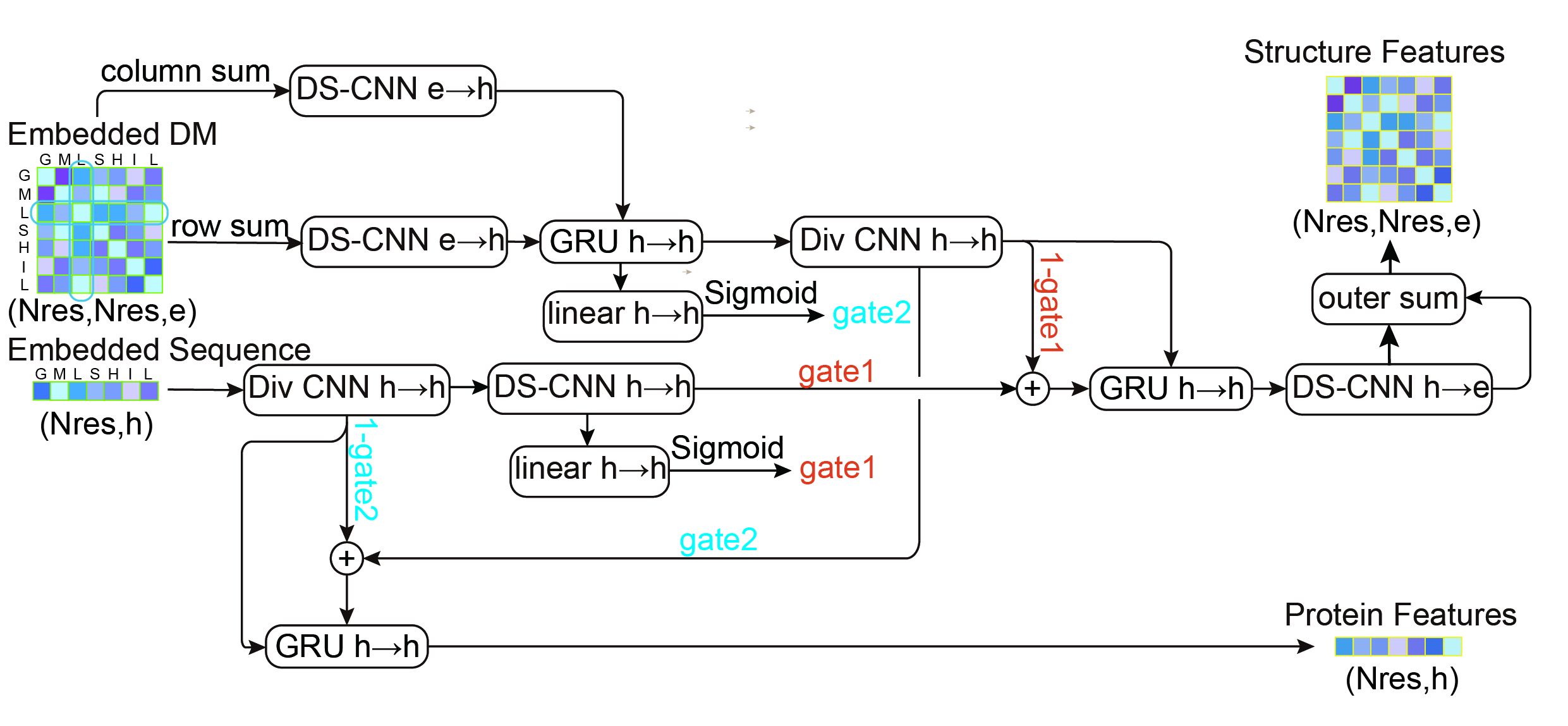

**Supplementary Fig. 18 The protein Evo-Updating block.** The protein residue sequence and structure features are coevolutionarily updated through the Evo-Updating block, so that the sequence features include structure information and forcing the structure features to contain sequence information. Abbrev: DS-CNN: Depthwise Separable Convolution Neural Network, Div CNN: Diversification Convolution neural network.

**Define:** $F_{N_{res}, h}^{seq}$ and $F_{N_{res}, e}^{DDM}$ are the features of the residue sequence and *DDM* embedded by the Prot-Aggregation algorithm. ${mask}_{N_{res}}^{Seq}$ and ${mask}_{N_{res},N_{res}}^{DDM}$ are the mask matrices of the protein residue sequence and $DDM$, respectively.

**def** *EvoUpdating*({$F_{N_{res}, h}^{seq}$},{$F_{N_{res}, e}^{DDM}\},\{{mask}_{N_{res}}^{Seq}\},\{{mask}_{N_{res},N_{res}}^{DDM}$}):

$$PairKey1 = DeepSparseCNN(RowSum(F_{N_{res}, e}^{DDM}))$$

$$PairKey2 = DeepSparseCNN(ColumnSum(F_{N_{res}, e}^{DDM}))$$

$$PairKey2 = DeepSparseCNN(ColumnSum(F_{N_{res}, e}^{DDM}))$$

$$MixKey = GRU(PairKey1, PairKey2)$$

$$Struct\_Features = DivCNN(MixKey)$$

$$Seq\_Features = DivCNN(F_{N_{res}, h}^{seq})$$

$$Seq2Struct = DeepSparseCNN(F_{N_{res}, h}^{seq})$$

$$SeqGate = Sigmoid(Linear(Seq2Struct))$$

$$StructGate = Sigmoid(Linear(MixKey))$$

$$Seq2Struct\_Vector = SeqGate\odot Seq2Struct+(1-SeqGate)\odot Struct\_Features$$

$$Struct\_Vector = GRU(Seq2Struct\_Vector, Struct\_Features)$$

$$Struct2Seq\_Mapping = DeepSparseCNN(Struct\_Vector)$$

$${F_{N_{res}, e}^{DistMat}}_{output} \leftarrow Struct2Seq\_Mapping\oplus Struct2Seq\_Mapping$$

$$Seq\_Vector = StructGate\odot Struct\_Features+(1-StructGate)\odot Seq\_Features$$

${F_{N_{res}, h}^{sequence}}_{output} = GRU$*(*$Seq\_Vector$*,* $Seq\_Features$*)*$\odot{mask}_{N_{res}}^{Seq}$

**return** $\boldsymbol{\to}$ {${F_{N_{res}, h}^{sequence}}_{output}$},$\{{F_{N_{res}, e}^{DistMat}}_{output}\}$

##### 2.3.5. Algorithm 7: Div CNN

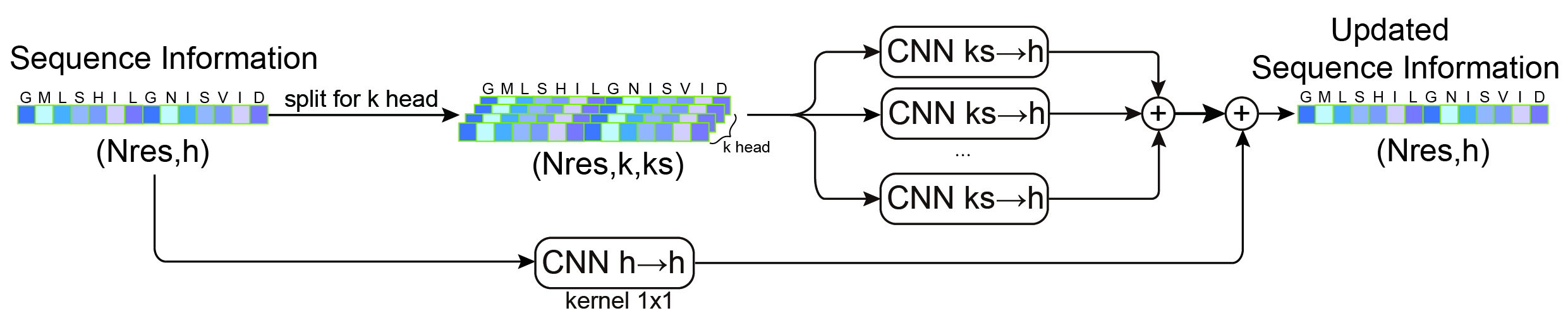

**Supplementary Fig. 19 The diversification convolution neural network (Div CNN) block.** Div CNN is used to enhance the diversification of structure features and sequence features with the multihead mechanism.

**def** *DivCNN* (*x*):

$$x0 = {CNN}_{kernel=1}(x)$$

\# x (seq, hidden_size) → $x_{k-head}$(seq, k, head_size)

\# where k is the number of heads, and head_size = hidden_size/k

$$x_{k-head} = TransposeForScores(x)$$

$$x_{total} = \sum_{n=k-head} {CNN}_{head\_size\to hidden\_size}(x_{k-head})+x0$$

**return** $\boldsymbol{\to}$ {$x_{total}$}

##### 2.3.6. Algorithm 8: TransposeForScores

**def** *TransposeForScores({input}):*

*# input dimension:*$(N_{res}/N_{atom}, h)$

\# output dimension:$(k, N_{res}/N_{atom}, ks)$

$$output = DimentionReshape(input)$$

**return** $\boldsymbol{\to}$ {$output$}

#### 2.4. Block III: Affinity Learning Module

Based on an end-to-end architecture, the protein and compound features extracted from the upstream model (included in the protein extractor and compound extractor) are fed into the affinity learning module (Supplementary Fig. 20. The mapping information between proteins and compounds is constructed as a pairwise matrix through the matrix multiplication operation to achieve feature aggregation between proteins and compounds, enabling the fitting and learning of the potential interaction information between the proteins and the compounds. Finally, the CPA predictions are given.

##### 2.4.1. Algorithm 9: Affinity Prediction

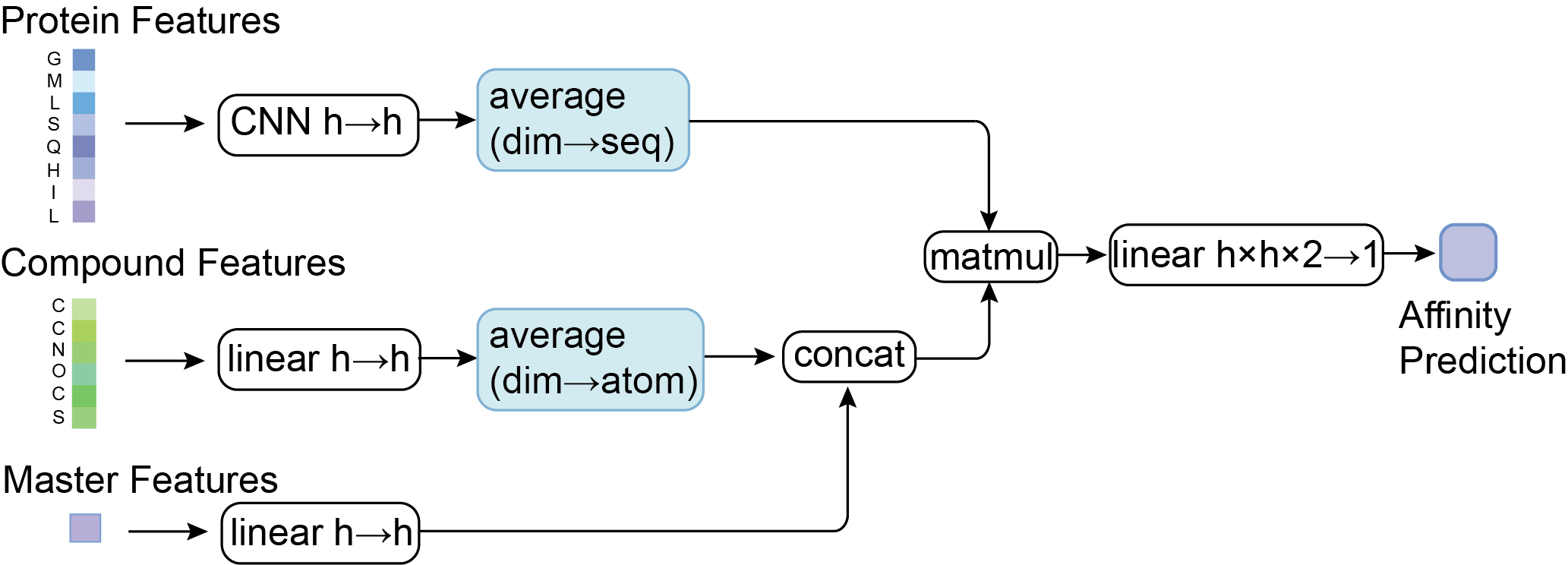

**Supplementary Fig. 20 Architecture of the affinity learning module.**

**Define** $F_{N_{atom}, h}^{Compound}$ and $F_{1, h}^{Master}$ as the compound atom features and master features extracted from the compound extractor algorithm, respectively. $F_{N_{res}, h}^{Protein}$ denotes the protein features extracted from the protein extractor algorithm that contain both the sequence and structure information of the protein. ${mask}_{N_{res}}^{Seq}$ and ${mask}_{N_{atom}}^{vertex}$ are the mask matrices of the protein residue sequence and the vertex in the compound graph.

**def** *AffinityPrediction*({$F_{N_{atom}, h}^{Compound}\},\{F_{N_{res}, h}^{Protein}\}, \{F_{1, h}^{Master}\},\{{mask}_{N_{res}}^{Seq}\},\{{mask}_{N_{atom}}^{vertex}\}$):

\# Inputs projections

${feature}_{init}^{Compound}=Linear$($F_{N_{atom}, h}^{Compound}$)

${feature}_{init}^{Protein}=CNN$($F_{N_{res}, h}^{Protein}$)

${feature}^{Master Node}=Linear$($F_{1, h}^{Master}$)

$${feature}_{final}^{Compound}\leftarrow(\sum_{a=N_{atom}} {{feature}_{init}^{Compound}}_{a}\odot{mask}_{a}^{vertex})/\sum_{a=N_{atom}} {mask}_{N_{atom}}^{vertex}$$

$${feature}_{final}^{Protein}\leftarrow(\sum_{s=N_{res}} {{feature}_{init}^{Protein}}_{s}\odot{mask}_{s}^{Seq})/\sum_{s=N_{res}} {mask}_{N_{res}}^{Seq}$$

$${feature}_{mixture}^{Compound}={concat}_{h}({feature}_{final}^{Compound},{feature}^{Master Node})$$

\# Output projection

$${Affinity}_{Prediction}=Linear(matmul({feature}_{mixture}^{Compound}, {feature}_{final}^{Protein}))$$

**return**$\to\left\{ {Affinity}_{Prediction} \right\}$

### 3. Supplementary Methods

**3.1 Calculation details for the discrete distance matrix and torsion angle matrix**

To calculate the distance between the residues and construct the discrete distance matrix (DDM) in each protein, we followed the steps listed below:

1. The 3D position of each residue in a protein was represented by the amino acids’ beta carbon position for all amino acids except glycine because glycine does not have beta carbon; therefore, for glycine the position of its alpha carbon is used instead;
2. Then, based on the represented 3D position of each residue in a protein, we calculated the Euclidean distance between each residue to construct a distance matrix (N_res_×N_res_, where N_res_ is the residue number in the protein) of the protein;
3. The distance between every two residues was discretized into 40 bins: the number tokens from 1 to 38 represent 38 bins of equal width between 3.25 Å and 50.75 Å, 0 represents distances smaller than 3.25 Å, and 39 represents distances larger than 50.75 Å.

Ultimately, a discrete distance matrix with lower storage and calculation requirements was constructed.

The torsion angle matrix was calculated through the following steps:

1. We first calculated the ψ (the torsion between Cα-C) and Φ (the torsion between N-Cα) angles in each residue;
2. Then, sine and cosine functions were applied to encode the torsion angles of ψ and Φ to accurately represent the torsion information of each protein;

Ultimately, a torsion angle matrix with the dimensions of N_res_×4 was constructed.

#### 3.2 Parameter Settings of the FeatNN

In this work, for fast and convenient calculation, we utilized 6 layers of deep GCN blocks and 2 layers of Evo-Updating blocks. The hidden size in the entire architecture was set as 128. The number of attention heads in the deep GCN blocks and the Evo-Updating blocks was 4. The detailed parameter settings can be found in Supplementary Table 2.

**3.3 Details of Dataset Construction from PDBbind, BindingDB and Binding MOAD**

The datasets constructed from the general set of PDBbind contain the CPA values with the K_i_, K_d_, and IC_50_ measurements between drugs and proteins, while datasets constructed from the refined set of PDBbind only contain the CPA values with the measurements of K_i_ and K_d_. BindingDB is rich in IC_50_ measured values (more than 500 thousand data), while the collections of the measurement values based on K_i_ and K_d_ are significantly smaller (40 thousand K_i_ measurements and 28 thousand K_d_ measurements were recorded). In this paper, to construct large datasets from BindingDB, we only selected the measured IC_50_ values to generate training data. To test the generalization ability of the models, we constructed new datasets from the Binding MOAD database and excluded the complexes that appeared in the datasets (training, validation, and test datasets) constructed from PDBbind. For a fair comparison of the generalization ability, we limit the datasets constructed from Binding MOAD with the measurement of IC_50_ and KIKD (K_i_ and K_d_) to the same amount of data. Thus, we constructed the dataset with IC_50_ and KIKD measurements from the “all of Binding MOAD” and “nonredundant MOAD” sets in the Binding MOAD database.

**3.4 Molecular Similarity Calculation**

Molecular structures were represented by 1024-dimensional binary Morgan fingerprints with radii of 2, while the Tanimoto coefficient was utilized to measure molecular similarities. Finally, the compounds in the dataset, according to similarity thresholds from 0.3 to 0.6 (with a step of 0.1), and similar compounds were grouped into the same dataset (training, valid or test set) to prevent data leakage.

**3.5 Homologous Protein Calculation**

The homology between proteins was quantified by multisequence alignment (MSA) methods, and based on the thresholds from 0.3 to 0.6 (with a step of 0.1), the obtained similarity scores were applied to divide the homologous proteins into the same subset to ensure that similar proteins did not appear in the same dataset (training, valid or test set), which is similar to the method in Note 3.3.

**3.6 Generalization evaluation on Binding MOAD**

All generalization testing processes are evaluated on the dataset constructed from Binding MOAD. We selected all models trained on the refined set and general set of PDBbind-v2020 to investigate the differences in the generalization ability (Supplementary Fig. 6) when trained with different amounts or qualities of data. We further tested all of the generalization abilities of FeatNN^optm^ (Supplementary Fig. 7) to prove the effectiveness of the pretraining strategy that utilized the high- and low-quality data from PDBbind and BindingDB, respectively.

**3.7 Details of Optimization with a Pretraining Strategy**

FeatNN was pretrained for 32 epochs on the datasets generated by BindingDB and used as the initial fine-tuning model. Then, we froze the parameters of the compound extractor and trained (fine-tuned) for 30 epochs on the protein extractor and affinity learning module with the training dataset generated with the measurement of IC_50_ (because the BindingDB dataset that we constructed here only has the affinity values calculated from the IC_50_ data) based on the general set of PDBbind-v2020.

**3.8 Details of the Ablation Experiment**

We used the IC_50_ dataset constructed from PDBbind’s general set to generate the training datasets for the module ablation experiment. The other dataset generation steps and model parameter settings were the same as those used to train FeatNN on the benchmark datasets generated from the general set of PDBbind in the main text.

In the model architecture modification step, we directly deleted or replaced the module to be ablated with a simple linear layer. Finally, the RMSE, Pearson coefficient and R^2^ were selected to compare and evaluate the comprehensive performance of these models.

4. **Full Algorithm Details**

The pseudocodes for each module are available in the supplementary methods.

**Notations for the Operations Between Vectors and Matrix**

The definitions of operations and variables are listed as follows. We use $\oplus$ for the outer sum, $\odot$ for the elementwise product, namely, the Hadamard product, $\sigma\left( \cdot\right)$ for the sigmoid activation function $\sigma(x)=1/(1+e^{-x})$, $tanh(\cdot)$ for the tanh activation function $tanh(x)=(e^{x}-e^{-x})/(e^{x}+e^{-x})$, and $f(\cdot)$ for the Gaussian error linear unit (GELU) activation function $GELU(x)=0.5x(1+erf(\frac{x}{\sqrt{2}}))$, where $erf\left( \cdot\right)$ serves as the Gaussian error function, $erf\left( x \right)=\frac{2}{\sqrt{\pi}}\int_{0}^{x} e^{{-t}^{2}}dt$, and $softmax(x_{i})$ is used for the softmax function $exp(x_{i})/\sum_{i} exp(x_{i})$.

**4.1. Compound Extractor Module**

In this study, a graph representation of a compound is utilized to describe the specific correlation between its atom features and bond features. Given a graph representation $\{V, E\}$, vertices and edges are used to represent atom and bond featuresin the compound, respectively. More specifically,

$\left\{ V \right\}=\left\{ \text{element name},\text{aromatic type},\text{vertex degree},"atom valence" \right\}, in which$the features are encoded by a one-hot-encoding strategy and then are concatenated into an all-one vector as ${\{F_{i}^{atom}\in R^{h}\}}_{i=1}^{N_{a}}$ for each atom. Similarly, $\left\{ E \right\}=\left\{ "bond type","\mathrm{shape}" \right\}$ is also applied, obtaining the embedded bond feature vector as ${\{F_{j}^{bond}\in R^{h}\}}_{j=1}^{N_{b}}$, where $i=1,2,\ldots,N_{a}$, $j=1,2,\ldots, N_{b}$, $h$ is the dimensionality of the hidden size, $N_{a}$ is the number of compound atoms, and $N_{b}$ is the number of protein residues. Original atom features are defined as $F^{0}\in R^{N_{a}\times h}$, and master node features are defined as summaries of atom features, that is, $F^{master}=\sum_{i=1}^{N_{a}} F_{i}^{atom}$. Considering that there are $l_{c}$ graph convolution layers where $l_{c}=1,2,...,l_{comp}$ and $k_{c}$ attention heads where $k_{c}=1,2,...,k_{comp}$, $l_{comp}$ is the total number of graph convolution layers, and $k_{comp}$ is the total number of compound feature attention heads. The atom features, bond features, and master features in the $l_{c}th$ layer are defined as $F_{atom}^{l_{c}}$, $F_{bond}^{l_{c}}$and $F_{master}^{l_{c}}$, respectively, and the variables $V$ with $k_{c}$ heads are defined as $V^{k_{c}}$. For example, $F_{atom}^{l_{c},k_{c}}$ represents the atom features in the $l_{c}th$ layer of the GCN with $k_{c}$ heads. For a detailed description, see Supplementary Session 2.2.

**4.1.1 Multihead Attention Block**

Main vertex (atom) features are obtained with a multihead attention mechanism and the elementwise product operation. The main vertex features are updated as $v_{comain}^{l_{c}}$ in each layer:

$$v_{tmain}^{l_{c},k_{c}}=softmax(W_{vm}^{l_{c},k_{c}}(tanh(W_{vmain}^{l_{c},k_{c}}F_{atom}^{l_{c}-1})\odot F_{master}^{l_{c}-1}))\boldsymbol{\otimes}W_{ms}^{l_{c},k_{c}}F_{atom}^{l_{c}-1}$$

$$v_{main}^{l_{c}}=tanh({W_{cmat}^{l_{c}}[v_{tmain}^{l_{c},k_{c}}]}_{k_{c}})$$

$$v_{comain}^{l_{c}}=dropout(F_{atom}^{l_{c}-1})$$

where $W_{vmain}^{l_{c},k_{c}}\in R^{h\times h}$, $W_{vm}^{l_{c},k_{c}}\in R^{h\times h}$, $W_{ms}^{l_{c},k_{c}}\in R^{h\times h}$, and ${[\cdot]}_{k_{c}}$ indicates the integration of the information from multihead attention. A detailed description and the pseudocode are provided in Supplementary Section 2.2.2.

**4.1.2 Deep GCN**

The atom features are sequentially updated using a message passing unit and a graph warp unit at each iteration of the GCN.

$${mt}_{main}^{l_{c}}={W_{lu2}^{l_{c}}[v_{comain}^{l_{c}},\sum_{v_{k}\in Neighbor(v_{i})} f(W_{ln}^{l_{c}}{[v_{comain}^{l_{c}},F_{bond}^{l_{c}}]}_{h+h+bn})]}_{h+h}$$

where $bn$ is the shape or size of bond neighbors, ${[\cdot, \cdot]}_{m}$ indicates the concatenation operation on different dimensions, $W_{lu2}^{l_{c}}\in R^{2h\times h}$, and $W_{ln}^{l_{c}}\in R^{h+(h+bn)}$.

To avoid the oversmoothing problem in the graph convolution process, we use the initial vertex features $F^{0}$ as the identity information and the residual connection pathway:

$$r^{l_{c}}=(1-\alpha){mt}_{main}^{l_{c}}+\alpha F^{0}$$

$$v_{comp}^{l_{c}}=\theta W_{fu}^{l_{c}}r^{l_{c}}+(1-\theta)r^{l_{c}}$$

where $W_{fu}^{l_{c}}\in R^{h\times h}$, and both $\alpha$ and $\theta$ are hyperparameters.

Next, ${ht}_{master}^{l_{c}}=W_{mas}^{l_{c}}F_{master}^{l_{c}-1}$, ${ht}_{mas2m}^{l_{c}}=W_{mas2m}^{l_{c}}F_{master}^{l_{c}-1}$ is defined. Both the main vertex and master node features can be mutually updated through the graph warp unit and $GRU$ layers, and $GRUs$ are used to determine the proportions of the main vertex and master node features updated at layer $l_{c}$.

$$g_{main}^{l_{c}}=\sigma(W_{zm1}^{l_{c}}v_{comp}^{l_{c}}+W_{zm2}^{l_{c}}{ht}_{mas2m}^{l_{c}})$$

$${ht}_{main}^{l_{c}}=g_{main}^{l_{c}}{ht}_{mas2m}^{l_{c}}+(1-g_{main}^{l_{c}})v_{comp}^{l_{c}}$$

$$v_{comain}^{l_{c}}={GRU}_{main}({ht}_{main}^{l_{c}},v_{comain}^{l_{c}-1})$$

With the same process, the master node features are also updated as $v_{comaster}^{l_{c}}$ in each graph convolution layer, that is,

$$g_{master}^{l_{c}}=\sigma(W_{zs1}^{l_{c}}{ht}_{master}^{l_{c}}+W_{zs2}^{l_{c}}v_{main}^{l_{c}})$$

$${ht}_{master}^{l_{c}}=g_{master}^{l_{c}}v_{main}^{l_{c}}+(1-g_{master}^{l_{c}}){ht}_{master}^{l_{c}}$$

$$v_{comaster}^{l_{c}}={GRU}_{master}({ht}_{master}^{l_{c}},v_{comaster}^{l_{c}-1})$$

where $W_{mas2m}^{l_{c}}\in R^{h\times h}$, $W_{mas}^{l_{c}}\in R^{h\times h}$, $W_{zs1}^{l_{c}}\in R^{h\times h}$, and $W_{zs2}^{l_{c}}\in R^{h\times h}$.

After the iterations of the deep GCN block, the final features of the main vertex and master features are obtained as $v_{comain}^{l_{comp}}$ and $v_{comaster}^{l_{comp}}$ that are defined above as $F^{fatom} and F^{fmaster}$. A detailed description and pseudocode are provided in Supplementary Section 2.2.1.

**4.2. Protein Extractor Module**

**4.2.1 Protein Aggregation Module**

Sequence and distance features are embedded through a word embedding strategy, and torsion features are embedded through a linear layer. The protein embedding module takes sequence features ${\{F_{n}^{seq}\in R^{h}\}}_{n=1}^{N_{res}}$, the DDM ${\{\boldsymbol{F}_{n}^{DDM}\in R^{e}\}}_{n=1}^{N_{res}\times N_{res}}$ and the torsion matrix ${\{\boldsymbol{F}_{n}^{TM}\in R^{h}\}}_{n=1}^{N_{res}}$ of proteins as input data. In addition, $N_{ts}$ is the initial torsion dimension, $h$ is the hidden size, $e$ is the embedding size, $k$ is the number of attention heads and $k_{h}$ is the hidden size of the attention heads, where $k_{h}=h/k$. A linear layer is used to update these features, that is,

$$l_{n}^{seq}={CNN}_{e\to h}^{seq}F_{n}^{seq}$$

$$\boldsymbol{E}_{i,j}^{DDM}=I_{n}F_{n}^{DDM}$$

$$l_{n}^{TM}={CNN}_{e\to h}^{tor2}f\left( {CNN}_{N_{ts}\to e}^{tor1}\boldsymbol{F}_{n}^{TM} \right)$$

where $I_{n}$ is the identity matrix, $N_{res}$ is the length of the amino acid in each protein, $N_{ts}$ is the initial dimensionality of the torsion size,$h$ is the hidden size, $m$ is the kernel size $e$ is the embedding size, and $pe$ is the preembedding size. All ${CNN}_{e\to h}^{seq}, {CNN}_{N_{ts}\to e}^{tor1}$ and ${CNN}_{e\to h}^{tor2}$ retain the width and height of the input matrix but change the feature dimensions with specific kernel sizes and padding sizes.

The aggregation of the protein sequence, distance and torsion features together is a novel strategy for use prior to the extraction of protein features.

$torgate=\sigma(W_{gt}l_{n}^{TM}$)

$$\boldsymbol{E}_{n}^{seq}=torgate\odot l_{n}^{TM}+(1-torgate)\odot l_{n}^{seq}$$

where $W_{gt}\in R^{h\times h}$.

Ultimately, the embedded sequence vector ${\{E_{n}^{seq}\in R^{h}\}}_{n=1}^{N_{res}}$ and embedded DDM${\{\boldsymbol{E}_{i,j}^{DDM}\in R^{e}\}}_{i=1, j=1}^{N_{res}\times N_{res}}$ are obtained from the protein embedding block. A detailed description and pseudocode are provided in Supplementary Section 2.3.3.

**4.2.2 Evo-Updating Module**

We use an evolutionary updating strategy to update the sequence and structure features in the Evo-Updating model block by combining the information derived from the protein embedding block.

The sequence features ${\{E_{n}^{seq}\in R^{h}\}}_{n=1}^{N_{res}}$ and structure features ${\{\boldsymbol{E}_{i, j}^{DDM}\in R^{e}\}}_{i=1, j=1}^{{N_{res}\times N}_{res}}$ are embedded via the protein aggregation algorithm, and $i, j$ are the row and column of the embedded DDM, respectively. We define the input features in layer $l_{p}$ as $F_{seq}^{l_{p}-1}$ and $\boldsymbol{DDM}_{i,j}^{l_{p}-1}$, where $l_{p}=1,2, \ldots,L_{p}$.

$${key}_{mix}^{l_{p}}=GRU({{CNN}_{e\to h}^{row}}^{l_{p}}\sum_{i=1}^{n} \boldsymbol{DDM}_{i,j}^{l_{p}-1}, {{CNN}_{e\to h}^{column}}^{l_{p}}\sum_{j=1}^{n} \boldsymbol{DDM}_{i,j}^{l_{p}-1})$$

$${key}_{mix}^{l_{p}, kn}=TransposeForScores({key}_{mix}^{l_{p}})$$

$${struct}^{l_{p}}=\sum_{M=1}^{k} {{CNN}_{k_{h}\to h}^{sn}}^{l_{p}, M}{key}_{mix}^{l_{p}, M}+{{CNN}_{h\to h}^{init\_res}}^{l_{p}}{key}_{mix}^{l_{p}}$$

Protein features are first processed through the gated recurrent unit (GRU) cell with row and column pooling features of $\boldsymbol{DDM}_{i,j}^{l_{p}-1}$, while all ${{CNN}_{e\to h}^{row}}^{l_{p}}$, ${{CNN}_{e\to h}^{column}}^{l_{p}}$, ${{CNN}_{kn\to h}^{sn}}^{l_{p}, kn}$ and ${{CNN}_{h\to h}^{init\_res}}^{l_{p}}$ models retain the width and height of the input matrix but change the feature dimensions with specific kernel sizes and padding sizes. In particular, $TransposeForScores()$ is an algorithm described in Supplementary Section 2.3.6. We use the outer sum operation to update and map the information derived from the sequence and use multihead attention to learn the diversified correlation of $\boldsymbol{DDM}_{i,j}^{l_{p}-1}$:

$${prot\_vec}_{seq}^{l_{p}}=f(\sum_{sn=1}^{k} {{CNN}_{k_{h}\to h}^{zwei}}^{l_{p}, sn}F_{seq}^{l_{p}-1, kn}+{{CNN}_{h\to h}^{zwei\_res}}^{l_{p}}F_{seq}^{l_{p}-1})$$

$${seq}_{initial}^{l_{p}}={{CNN}_{h\to h}^{dz1}}^{l_{p}}{prot\_vec}_{seq}^{l_{p}}$$

$${seq2struct}^{l_{p}}=f({{CNN}_{h\to h}^{dz2}}^{l_{p}}{seq}_{initial}^{l_{p}})$$

We use a gate mechanism to gather more useful information from the input features and aggregate the sequence features onto the structure features, and the GRU cell is used to aggregate both updated and initial structure features,

$$g_{seq2str}^{l_{p}}=\sigma(W_{seq2str}^{l_{p}}{seq2struct}^{l_{p}})$$

$$g_{str2seq}^{l_{p}}=\sigma(W_{str2seq}^{l_{p}}{key}_{mix}^{l_{p}})$$

$$v_{struct}^{l_{p}}=g_{seq2str}^{l_{p}}\odot{seq2struct}^{l_{p}}+(1-g_{seq2str}^{l_{p}})\odot{struct}^{l_{p}}$$

$$v_{sequence}^{l_{p}}=g_{str2seq}^{l_{p}}\odot{struct}^{l_{p}}+(1-g_{str2seq}^{l_{p}})\odot{seq}_{initial}^{l_{p}}$$

$$p_{struct}^{l_{p}}={{CNN}_{h\to e}^{map}}^{l_{p}}f(GRU(v_{struct}^{l_{p}}, {struct}^{l_{p}}))$$

$$p_{sequence}^{l_{p}}=f(GRU(v_{sequence}^{l_{p}}, {seq}_{initial}^{l_{p}}))$$

where $W_{seq2str}^{l_{p}}\in R^{h\times h} and W_{str2seq}^{l_{p}}\in R^{h\times h}$ and ${{CNN}_{k_{h}\to h}^{zwei}}^{l_{p}, sn}$, ${{CNN}_{h\to h}^{zwei\_res}}^{l_{p}}$, ${{CNN}_{h\to h}^{dz1}}^{l_{p}}$ ${{CNN}_{h\to h}^{dz2}}^{l_{p}}$ and ${{CNN}_{h\to e}^{map}}^{l_{p}}$ retain the width and height of the input matrix but change the feature dimensions with specific kernel sizes and padding sizes.

We aggregate the features with the outer sum (to create a symmetric matrix with a highly correlated DDM) and the gate. The updated features of the sequence and DDM in the $l_{p}th$ layer of the Evo-Updating block are given as $F_{seq}^{l_{p}}$ and $\boldsymbol{DDM}_{i,j}^{l_{p}}$, respectively,

$$F_{seq}^{l_{p}}=I_{n}p_{sequence}^{l_{p}}$$

$$\boldsymbol{DDM}^{l_{p}}=p_{struct}^{l_{p}}\oplus p_{struct}^{l_{p}}$$

where $I_{n}$ is the identity matrix and $\oplus$ is as the outer sum operation.

After calculating $L_{p}$ iterations of the protein encoder, we obtain the final feature representations ${\{F_{seq,i}^{l_{p}}\in R^{h}\}}_{i=1}^{N_{res}}$ and ${\{\boldsymbol{DDM}_{i,j}^{l_{p}}\in R^{e}\}}_{i=1,j=1}^{{N_{res}\times N}_{res}}$. A detailed description and pseudocode are provided in Supplementary Section 2.3.4. All specific information can be found in Supplementary Section 2.3.

**4.3. Affinity Learning Module**

The affinity learning module integrates the mutual information between the compounds and proteins during noncovalent interaction affinity prediction. Suppose we are given atom features ${\{F_{i}^{fatom}\in R^{h}\}}_{i=1}^{N_{a}}$ and master node features $\{F^{fmaster}\in R^{h}\}$ from the compound extractor, as well as protein features ${\{F_{i}^{seq}\in R^{h}\}}_{i=1}^{N_{res}}$ extracted from the protein extractor. In particular, both the compound and protein features are separately transformed into a compatible space by single linear layers, that is, $F_{atom}^{Comp}=f(W_{atom}F_{i}^{fatom})$ and $F_{seq}^{Prot}=f({CNN}_{h\to h}^{Fseq}F_{i}^{seq})$, where $i=1,2,..,N_{a}$, $j=1,2,.., N_{res}$, $W_{atom}\in R^{h\times h}$, and ${CNN}_{h\to h}^{Fseq}$ retains the width and height of the input matrix but changes the feature dimensions with specific kernel sizes and padding sizes.

The protein and compound features are eventually calculated after the$l_{aff}th$ iteration. Prior to performing affinity prediction, the feature aggregation operation between the master node features and main graph features should be considered with the help of a summation operation, that is,

$$C_{final}=\sum_{{a=N}_{a}} {F_{atom}^{Comp}}_{a}/N_{a}$$

$$C_{aggre}={[C_{final}, F^{fmaster}]}_{h+h}$$

The same operations are also utilized for the protein features, that is,

$$P_{final}=\sum_{{s=N}_{res}} {F_{seq}^{Prot}}_{s}/N_{res}$$

where ${[\cdot, \cdot]}_{m}$ indicates the concatenation operation between the hidden sizes of the main vertex features and master node features.

Finally, with a single linear mapping layer, the affinity value is calculated by vectors $C_{aggre}$ and $P_{final}$, that is,

$$affinity=W_{aff}(f(C_{aggre}P_{final}))$$

where $W_{aff}\in R^{2h^{2}\times1}$.

A detailed description and pseudocode are provided in Supplementary Section 2.4.

**4.4. Quantification and Statistical Analysis**

Evaluation Metrics

We use eight metrics that are commonly used for this problem to evaluate the prediction performance of our model. These metrics are defined as follows.

The R^2^ score, RMSE, and Pearson coefficient are often used in regression analysis. They describe the distance between the predicted values and true values. The higher the values of R2 and the Pearson coefficient are, the closer the model prediction results are to the real values. The smaller the RMSE value is, the smaller the error in the prediction value, that is, the higher the accuracy.

The RMSE is the standardized value of the MSE that is typically used as the training loss in machine learning studies. It is defined as follows:

$$RMSE\left( y,\hat{y} \right)=\sqrt{\frac{1}{n}\sum_{i=1}^{n} \left( y_{i}-\hat{y_{i}} \right)^{2}}$$

The R^2^ score is a dimensionless score describing the effectiveness of the model. It compares the output prediction to a random guess according to the average of the true values:

$$\begin{matrix} R^{2}\left( y,\hat{y} \right) & =1-\frac{SS_{residual}}{SS_{total}} \\ & =1-\frac{\sum_{i} \left( y_{i}-\hat{y_{i}} \right)^{2}}{\sum_{i} \left( y_{i}-\overline{y_{i}} \right)^{2}} \end{matrix}$$

We use a coefficient that can describe the correlation between the predicted values and true values: namely the Pearson product-moment correlation coefficient.

The Pearson correlation coefficient describes the linear correlation between two values and is defined as:

$$\begin{matrix} Pearson\left( y,\hat{y} \right) & =\frac{Cov\left( y,\hat{y} \right)}{\sigma_{y}\sigma_{\hat{y}}} \\ & =\frac{\sum_{i} \left( y_{i}-\overline{y} \right)\left( \hat{y_{i}}-\overline{\hat{y_{i}}} \right)}{\sqrt{\sum_{i} \left( y_{i}-\overline{y_{i}} \right)^{2}}\sqrt{\sum_{i} \left( \hat{y_{i}}-\overline{\hat{y_{i}}} \right)^{2}}} \end{matrix}$$

where $y_{i}$ are the prediction values, and $\hat{y_{i}}$ are the true values in the dataset, $i=$1,2...,n, where n is the total amount of the dataset.

In this paper, Pearson was selected to evaluate the accuracy of CPA prediction when predicting the affinity of 28 bioactive small molecules binding to SARS-CoV-2 3C-like protease, and the calculation and statistical method are consistent with the above description.
